# Tabula Glycine: The whole-soybean single-cell resolution transcriptome atlas

**DOI:** 10.1101/2024.07.08.602332

**Authors:** Sergio Alan Cervantes-Pérez, Sandra Thibivilliers, Sahand Amini, Julie M. Pelletier, Ian Meyer, Hengping Xu, Sutton Tennant, Pengchong Ma, Chandler M. Sprueill, Andrew D. Farmer, Jeremy E. Coate, Hilde Nelissen, Qiuming Yao, Olivier C. Martin, Erik J. Amézquita, Robert B. Goldberg, John J. Harada, Marc Libault

## Abstract

Soybean (*Glycine max*) is an essential source of protein and oil with high nutritional value for human and animal consumption. To enhance our understanding of soybean biology, it is essential to have accurate information regarding the expression of each of its 55,897 protein-coding genes. Here, we present “Tabula Glycine”, the soybean single-cell resolution transcriptome atlas. This atlas is composed of single-nucleus RNA-sequencing data of nearly 120,000 nuclei isolated from 10 different *Glycine max* organs and morphological structures comprising the entire soybean plant. These nuclei are grouped into 157 different clusters based on their transcriptomic profiles. Among genes, the pattern of activity of transcription factor genes is sufficient to define most cell types and their organ/morphological structure of origin, suggesting that transcription factors are key determinants of cell identity and function. This unprecedented level of resolution makes the Tabula Glycine a unique resource for the plant and soybean communities.

## Introduction

Soybean (*Glycine max*) is an economically important legume crop, producing ∼25% of the proteins and 30% of the oil used for livestock and human consumption [1, 2]. Additionally, soybean has a positive ecological footprint on agriculture with its capability to symbiotically interact with nitrogen-fixing soil bacteria (i.e., *Bradyrhizobium diazoefficiens*) [3]. This interaction minimizes the application of inorganic nitrogen fertilizers, supporting sustainable agricultural practices. Considering the current predictions of world population growth, the demand for soybean products should double by 2050 [4]. To reach this demand, soybean biologists need to maximize soybean yield and enhance other important traits (e.g., oil and protein content and quality, biotic and abiotic stress resistance, and biological nitrogen fixation). In addition to ongoing breeding programs, the emergence of synthetic biology strategies and genome editing technology [5, 6] allow researchers to finely regulate the activity of selected genes to improve a specific trait. However, to reach its full potential, there is a need to gain deeper knowledge of gene function, expression patterns, and the regulatory mechanisms controlling gene activity. As described below, our understanding of soybean gene activity at the scale of the entire plant is currently limited to organ-level averages, and we lack expression data at single cell-or cell-type resolution.

In 2010, building on the completion of the soybean genome sequence [7], two whole-organ transcriptome atlases were generated to measure the transcriptional profiles of the soybean genes across the entire plant [8, 9]. Concomitantly to the release of these transcriptome atlases, several online resources such as SoyBase [10], SoyXpress [11], SoyDB [12]], SoyKB [13], and ePlant Soybean [14] were released allowing researchers to access the expression profile of their genes of interest. While valuable, the current soybean transcriptome atlases do not take into consideration the multicellular complexity of each soybean organ/morphological structure, referred to as “organ” in the rest of the manuscript, and the fact that each cell type composing the plant is characterized by a unique transcriptomic profile.

Plant single-cell transcriptomics, a technology that recently emerged to decipher the transcriptomic profile of all and even rare cell types [15], represents an attractive solution to precisely characterize the transcriptional profile of each cell type composing the soybean plant. First extensively applied to the Arabidopsis root system, researchers used isolated protoplasts as an input to single-cell RNA-seq experiments [15]. However, protoplast-based single-cell RNA-seq technology has several limitations, especially when applied to non-model species [16]. First, the breadth of cell types analyzed depends on the digestion of the cell wall. Considering the differential composition of the cell wall between species, organs, and cell types (e.g., suberization of the exodermis of the tomato root [17] or of the endodermis layer of the Arabidopsis root [18]), the implementation of plant single-cell transcriptomics on isolated protoplasts would require the optimization of the cocktail of enzymes for each plant organ and developmental stages. Second, the transcriptomic information collected from isolated protoplasts includes transcripts synthesized in response to protoplastization [19–21]. Third, the bursting of fragile protoplasts during the early stages of the cDNA library construction is another limitation of their use. In addition to losing the cells, protoplast bursting also generates significant transcriptional noise.

To overcome these limitations and develop a high-quality single-cell resolution transcriptome atlas of the soybean plant, we applied single-nucleus RNA-seq (sNucRNA-seq) using the droplet-based 10x Genomics technology on 10 different organs of the soybean plant. We report the transcriptome of almost 120,000 soybean nuclei isolated from 10 organs clustered into 157 groups. Our analysis reveals that, on average, less than 400 nuclei per cluster are sufficient to cover over 95% of the transcriptome of a specific soybean cell type, a reflection of the quality and depth of the transcriptomic information collected. A focus on the activity of transcription factor genes (TFs) reveals that their co-expression is critical to control the biology of specific organs and cell types. In this manuscript, we present Tabula Glycine, the first single-cell resolution and comprehensive transcriptomic resource of the soybean plant, an atlas that covers all the organs of this crop.

## Results

### Creation of Tabula Glycine, the single cell resolution soybean transcriptome atlas

To capture the transcriptomes of most of the cell types and cell states composing the soybean plant and create a comprehensive single-cell resolution soybean transcriptome atlas, we applied sNucRNA-seq technology to at least two and at most three independent biological replicates from ten representative organs of the soybean plant. In addition to our previously published root tip and nodule atlases [22], Tabula Glycine also includes the single cell transcriptome atlases of the shoot apical meristem, unifoliate leaf, trifoliate leaf, floral bud, green pod without seeds and seeds collected at different developmental stages (i.e., at the heart, cotyledon, and mid-maturation stages; Fig.1A). A total of 140,728 nuclei were analyzed. Upon a stringent quality check and filtration of the processed nuclei (see Methods), we selected 116,525 high-quality nuclei for further analysis (∼83% of nuclei recovery; Fig.S1 and [22]). The number of analyzed nuclei per organ ranges from 4,703 for green pods (two replicates) to 20,154 for seeds at their mid-maturation stage (three replicates) (Fig.S2; Table.S1). The average number of high-quality nuclei per organ is 11,652 with, on average, 2,919 unique molecular identifiers (UMIs; Fig.1B) and 1,931 expressed genes detected per nucleus (Fig.1C). Upon integration, an average of 42,903 protein-coding genes were found expressed per organ (Table.S1) with a total of 51,770 expressed protein-coding genes across the entire Tabula Glycine atlas (i.e., 92.6% of the 55,897 predicted protein-coding genes).

**Figure 1.**
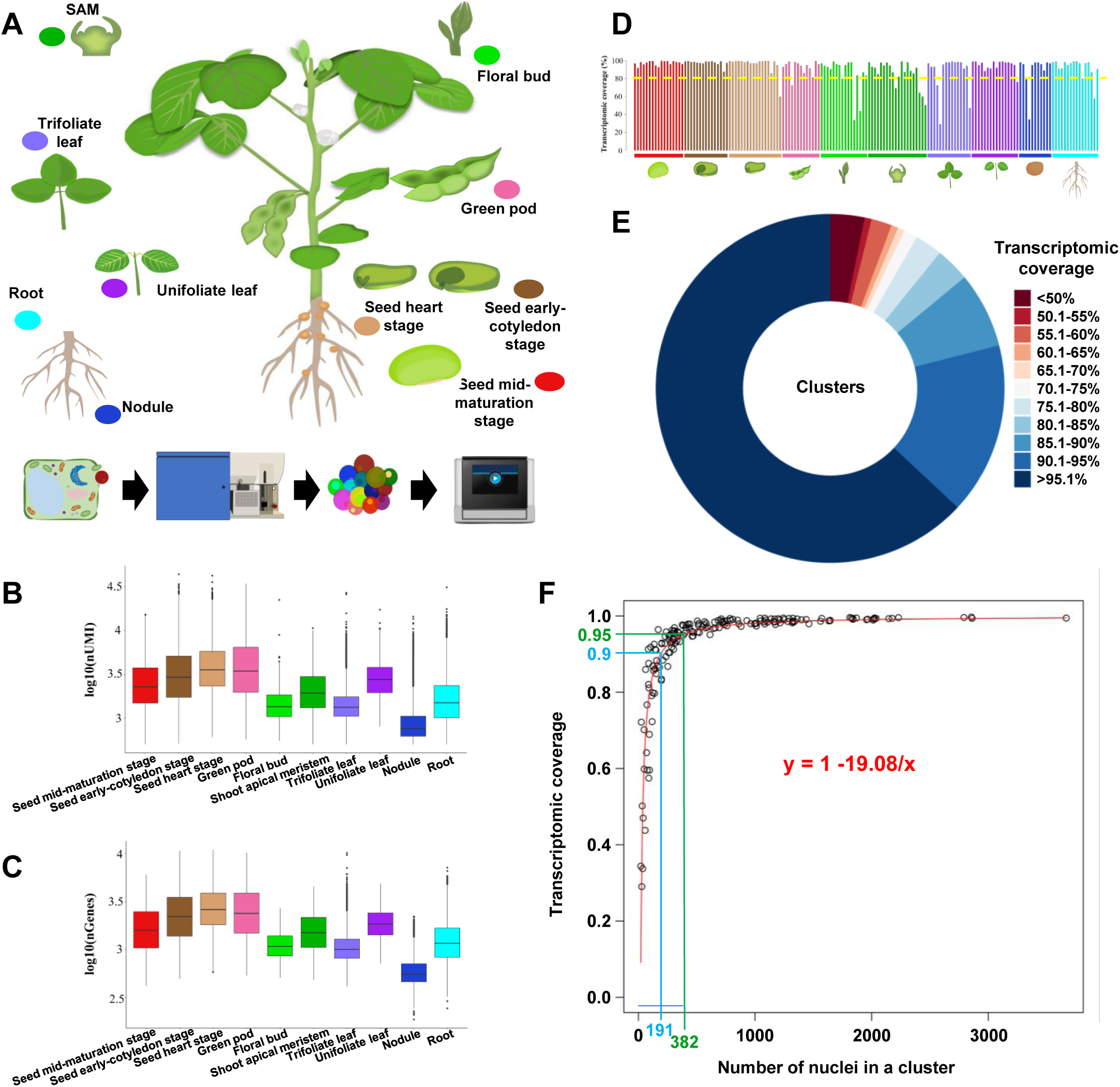
Establishment of Tabula Glycine, the soybean single-nucleus transcriptome atlas. **A.** The soybean cell atlas is generated from 10 collected tissues, including several seed developmental stages (i.e., heart, early cotyledon, and mid-maturation stages), green pods, flower bud, shoot apical meristem (SAM), leaves (trifoliate and unifoliate leaves), nodule and root (see joint manuscript). Bottom: Schematic representation of the experimental design used to generate single-nucleus RNA-seq (sNucRNA-seq) libraries using the 10X Genomics platform. The soybean nuclei were sorted to ensure their purification. **B and C**. Number of Unique Molecular Identifiers (UMI; **B**) and expressed genes per nucleus (**C**) in each tissue composing Tabula Glycine. **D.** Percentage of transcriptomic coverage of the 157 different cell clusters composing the soybean atlas using Shannon’s entropy measure. **E.** Distribution of the 157 Tabula Glycine clusters based on the percentage of their transcriptomic coverage. **F.** Estimation of the transcriptomic coverage of Tabula Glycine based on the number of cells using the entire soybean cell atlas by fitting a Pareto cumulative distribution function.

To estimate the transcriptional depth of Tabula Glycine and its quality compared to existing bulk RNA-seq resources, we conducted a comparative analysis between the pseudo-bulk single nucleus transcriptome of four organs (i.e., green pod, flower, shoot apical meristem, unifoliate leaves) and the previously published transcriptomes of whole soybean organs [9] (Fig.S3). In addition to confirming the expression of most of the genes previously reported using whole-organ RNA-seq technology [9] (93.4 to 95.3% of the genes found expressed in the whole-organ transcriptomes were confirmed in Tabula Glycine), the sNucRNA-seq dataset led to the identification of thousands of newly expressed genes (from 5,104 to 8,250 genes in the SAM and green pod, respectively; Fig.S3). This result suggests that, in addition to increasing the resolution of existing organ-based transcriptome atlases, Tabula Glycine also provides enhanced coverage of the transcriptome of each soybean organ.

The independent clustering of 10 soybean organs/seed developmental stages revealed a total of 157 cell clusters that vary from 21 nuclei for the “sclereid layer” cluster of the nodule [22] to 3,668 nuclei for cluster #14 of the seed at its mid-maturation stage (Table.S2). The number of biological entities per cluster, their nature (i.e., cells *vs.* nucleus; [18, 23]), and the depth of sequencing of the single cell/nucleus RNA-seq libraries [24], all impact the saturation rate of the transcriptome of a cluster. Therefore, we estimated the transcriptomic coverage of each of the 157 soybean clusters by calculating Shannon’s entropy using the Unique Molecular Identifier (UMI) matrix per gene (See Methods; Fig.1D). We found high transcriptomic coverage (>80%) for 140 clusters of the 157 Tabula Glycine clusters (89% of the clusters; Fig.1E; Fig.S4; Table.S2). Among them, 100 clusters are characterized by their very high transcriptomic coverage (>95%; Table.S2; Fig.1E). Considering the 17 remaining clusters characterized by a transcriptomic coverage below 80%, twelve clusters are composed of less than 100 nuclei (Table.S2). To estimate the number of nuclei needed to properly cover the transcriptome of a cluster, we first plotted the transcriptomic coverage percentage of the 157 Tabula Glycine clusters *versus* the number of nuclei composing each cluster (Fig.1F). By noticing that a large transcriptomic coverage is obtained with a modest number of nuclei across all clusters, we modeled this coverage fraction to nuclei number with a Pareto cumulative distribution function (i.e., coverage fraction = 1 – 19.08/nuclei# in the cluster; Fig.1F). Based on this equation, we estimate that 191 and 382 nuclei are needed to cover 90% and 95% of the transcriptome of a soybean cluster, respectively (Fig.1F). We assume that these values will fluctuate based on the plant species considered for a single-cell RNA-seq study, the nature of the organs analyzed such as the level of endoreduplication of the cells, its response to environmental stresses, and, according to our experience, based on the quality of the nuclear suspension used to generate the sNucRNA-seq libraries. Nevertheless, for the present data and considering our previous study and the work of others showing that the nuclear and cellular transcriptomes at the single-cell level are both highly correlated with whole organ transcriptomes [18, 23], we conclude that Tabula Glycine is a resource that provides accurate transcriptomic information for most of its 157 clusters.

### Cell type annotation of the Tabula Glycine clusters

In a previous study, we used Molecular Cartography™ technology to annotate all the 16 and 11 soybean root and nodule cell clusters [22], respectively. To functionally annotate the remaining 130 cell clusters identified from the 8 remaining organs composing Tabula Glycine, we implemented multiple approaches. For the clusters of the soybean canopy (i.e., the shoot apical meristem, unifoliate and trifoliate leaves, floral buds, and green pods), we analyzed the expression of soybean genes orthologous to *Arabidopsis thaliana* cell type marker genes assuming that their cell-type transcriptional specificity is extensively conserved. Our previous work conducted on the expression of root marker genes between *A. thaliana* and *Medicago truncatula* supports this assumption [24]. Accordingly, upon identifying 25,790 pairs of orthologous soybean-*A. thaliana* genes (Table.S3), we analyzed the expression patterns of the soybean genes orthologous to Arabidopsis cell-type-specific marker genes (Table.S4). This approach revealed many soybean orthologs to Arabidopsis marker genes co-expressed in the same soybean cluster (Fig.2; Figs.S5 to S7). This result supports conservation of cell-type-specificities between the expression of soybean and Arabidopsis orthologous marker genes.

**Figure 2.**
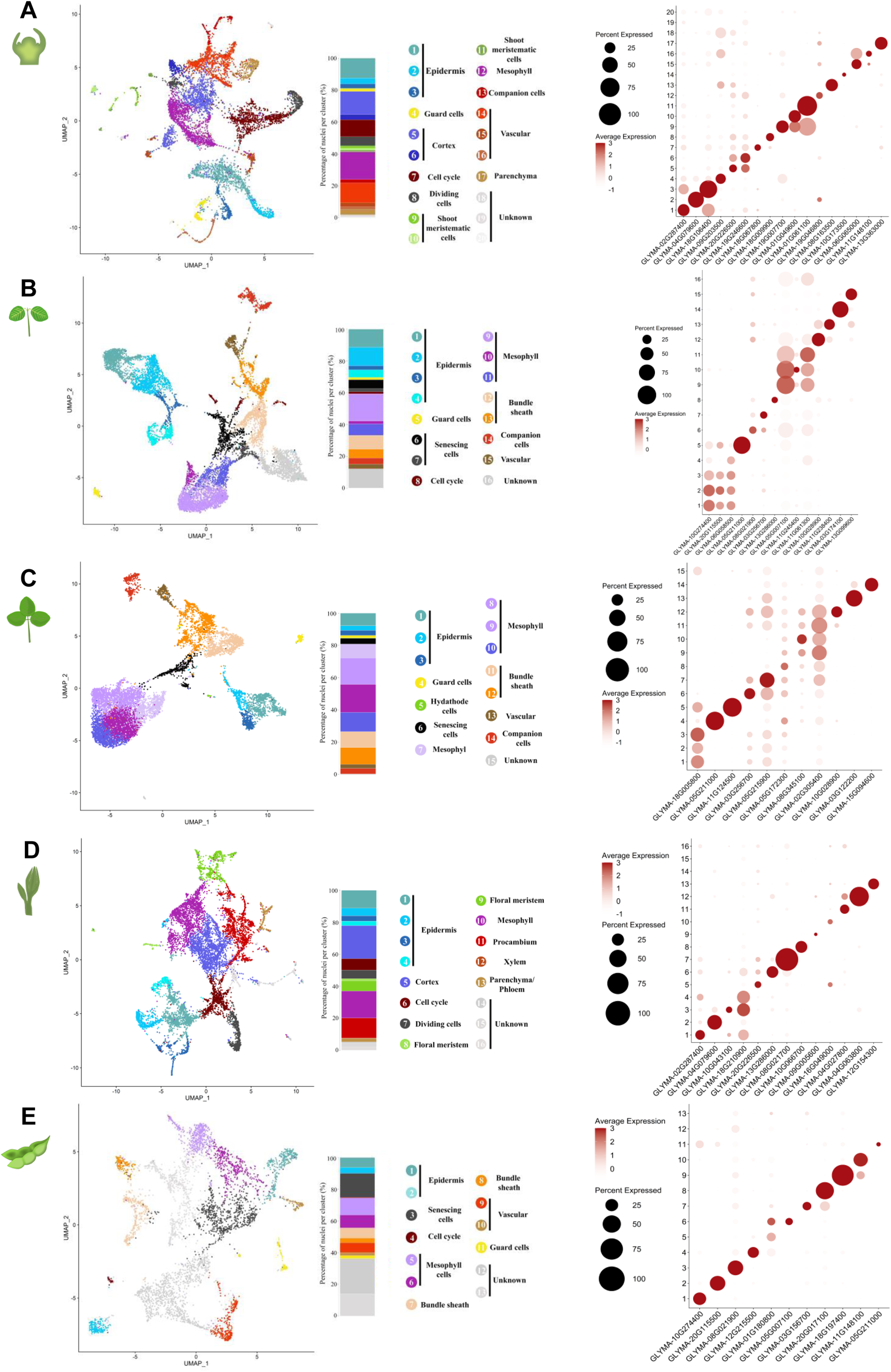
Functional annotation of the sNucRNA-seq clusters from the soybean canopy using ortholog genes. **A-E.** Left: UMAP plots of the shoot apical meristem (**A.**), unifoliate leaf (**B.**), trifoliate leaf (**C.**), floral bud (**D.**), and pod (**E.**) colored by distinct cell-cluster according to their transcriptomic profiles. Center: Stacked plots of the percentage of nuclei contributing to each cell cluster in a UMAP. Right: Dotplots of the expression pattern of a subset of cell-type specific gene markers used to functionally annotate the clusters of the soybean SAM (**A.**), unifoliate leaf (**B.**), trifoliate leaf (**C.**), floral bud (**D.**), and pod (**E.**) (see Figures S5 to S7 for details). The percentage of nuclei expressing the gene of interest (circle size) and the mean expression (circle color) of the genes are shown.

In several instances, we noticed that the same set of soybean marker genes can be used to annotate the same cell types of the unifoliate leaves, trifoliate leaves, and green pods, a plant organ sharing similar cellular features to leaves. For example, the epidermal cell clusters were annotated based on the expression patterns of soybean orthologs to the *A. thaliana CUT1* (AT1G68530) and *LPTG1* (AT1G27950) genes [25, 26] (Table.S4, highlighted green cells; Fig.2; Figs.S5 to S7). In the same organs, the mesophyll cells were functionally annotated based on the activity of soybean genes associated with the photosynthetic complex [e.g., *LHCA6* (AT1G19150) [27], *PSBY* (AT1G67740), *LHCB2.1* (AT2G05100), *PSBO2* (AT3G50820), and *RBCS2B* (AT5G38420) (Table.S4, highlighted blue cells)] and *EPFL9*, a gene also known as *STOMAGEN* (AT4G12970) and expressed in mesophyll cells to induce the formation of stomata (Table.S4, highlighted yellow cells) (Fig.2; Figs.S5 to S7). Similarly, to identify the cell clusters associated with the vasculature or the guard cells, we looked for the expression of ion and sugar transporter genes (Table.S4, highlighted purple cells), and genes orthologous to the *A. thaliana* guard cell marker genes *FMA* (AT3G24140) and *MPK5* (AT4G11330) and others [28], respectively (Table.S4, highlighted orange cells (Fig.2; Figs.S5 to S7). The expression patterns of the soybean orthologs to *A. thaliana WRKY46* (AT2G46400) and *WRKY53* (AT4G23810), genes promoting senescence [29–32], serve to annotate clusters #6 and 7, cluster #6, and cluster #3 as the “senescing cells” clusters in the soybean unifoliate leaves, trifoliate leaves, and green pod UMAPs, respectively (Table.S4, highlighted light blue cells; Fig.2; Figs.S5 to S7). Finally, soybean orthologs to *A. thaliana* genes controlling cellular proliferation and histone modification were used to annotate the clusters composed of cells engaged in the cell cycle. Using these marker genes and many others, we were able to annotate 15, 14, and 11 of the 16, 15, and 13 clusters composing the unifoliate leaves, trifoliate leaves, and green pod UMAPs (90.9% of annotated clusters; Fig.2B, C, and E). We applied a similar strategy to annotate 13 and 17 of the 16 and 20 clusters composing the floral bud and shoot apical meristem UMAPs, respectively (Table.S4, Fig.2; Figs.S5 and S6).

To annotate the soybean seed clusters, we adopted a different strategy. First, we identified cluster-specific genes for each cluster composing the three seed developmental stages represented in Tabula Glycine (i.e., heart, cotyledon, and mid-maturation stages). Then, we corroborated their cell-type specific expression upon mining the single-cell transcriptomic resource generated by Drs. Goldberg and Harada using Laser Capture Microdissection (LCM) technology (GEO accession number-GSE116036) and RNA-seq dataset from the seed-filling cells [33, 34] (Table.S4). This strategy allowed us to functionally annotate 15, 14, and 13 of the 18, 15, and 17 clusters composing the UMAP of the soybean seed heart, cotyledon, and mid-maturation stages, respectively (Fig.3).

**Figure 3.**
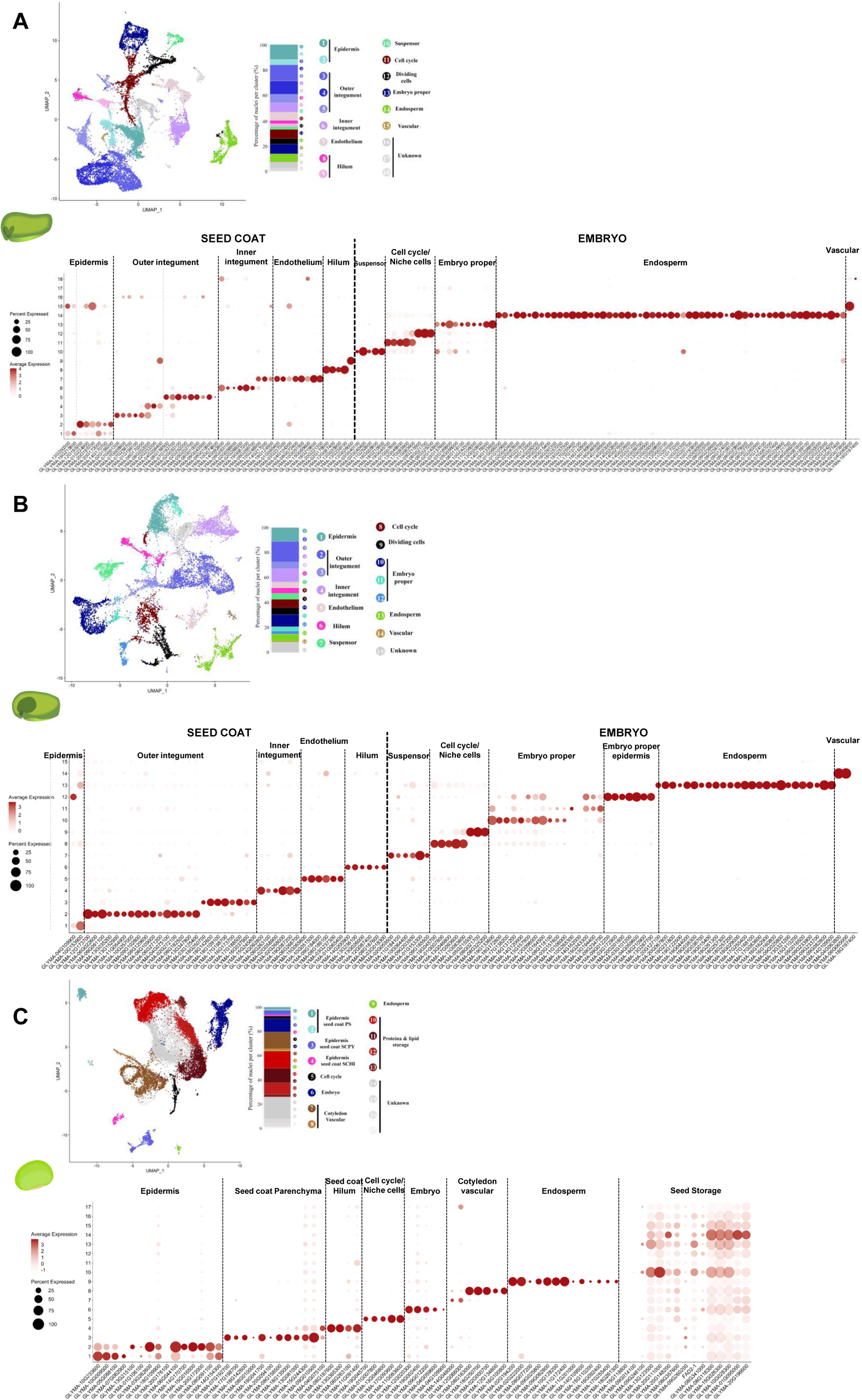
Functional annotation of the sNucRNA-seq clusters from the soybean seed using Laser Capture Microdissection RNA-Seq datasets. Top Left panel: UMAP and stacked plots of the percentage of nuclei contributing to the seed at the heart (**A.**), early cotyledon (**B.**), and mid-maturation stages (**C.**); Top right panel: schematic representation of the different tissues composing the seed at the heart (**A.**) and early cotyledon stages (**B.**); Bottom panel: Dotplots of the expression pattern of seed cell-type specific marker genes identified by LCM to support the functional annotation of the clusters of the seed heart, early cotyledon, and mid-maturation stages. The percentage of nuclei expressing the gene of interest (circle size) and the mean expression (circle color) of the genes are shown.

To better assess changes in the cellular composition and transcriptomic programs occurring during seed development, we also generated a unified UMAP of the soybean seed (Fig.S8A). The 19 clusters of this unified UMAP were annotated using the seed marker genes of the seed cell types as defined by LCM (Fig.S8B). This integrated analysis revealed striking differences in the transcriptome and cellular population of the seed during its development. During its early developmental stages, the soybean seed is represented by cells composing the epidermis and the outer and inner integuments. At the mid-maturation stage, the embryonic and storage parenchyma cells are over-represented (i.e., clusters #10 to 13 represent over 70 % of the cells transcriptionally analyzed; Fig.S8C and D). At the transcriptional level, the induction of the expression of the genes controlling the biosynthesis of cupins, seed storage proteins, and oleosins, proteins involved in the production of oil bodies, and genes encoding albumin [35–37] start as soon as the cotyledon stage of the seed (Fig.S8B, clusters #10 to 13, red square). Later during the development of the seed, their expression is extended to all the cells composing the mid-maturation seed including in the embryo (Fig.S8B). The co-induction of the expression of the oleosin-, cupin-, and protein storage-encoding genes in all the cell types composing the seed is not artefactual (e.g., the result of ambient noise) because such ubiquitous transcriptional pattern does not apply to cell-type marker genes of the soybean seed mid-maturation genes. As a result, and considering the annotation of the soybean root and nodule clusters [22], we confidently annotated 139 of the 157 clusters composing Tabula Glycine (88.5%).

### Evaluating the transcriptomic heterogeneity of the soybean cell types

Tabula Glycine allows us to evaluate the transcriptomic heterogeneity between soybean cell types. First, we assessed the transcriptomic breath of the 157 soybean clusters by analyzing the distribution of the numbers of UMIs (i.e., transcripts) (Fig.4A) and expressed genes per cluster (Fig.4B). These numbers vary between organs, potentially as the result of the differential transcriptomic activity of an organ (i.e., developing seed and green pods show higher transcriptomic activities than other organs) or as the result of technical bias during library construction. More interestingly, we observed variations in the number of UMIs and expressed genes between the clusters within the same organ suggesting differential transcriptomic activities between the cell types composing an organ. For instance, the endosperm clusters of the soybean seed at the heart and cotyledon stages (clusters #14 and 13, light brown and brown arrows, respectively, Fig.4A and B), as well as the Rhizobia-infected cell clusters of the soybean nodule (clusters F and G; Fig.4A and B, dark blue arrows, [22]), show over 1.5-fold increases in the number of UMIs per nucleus (Table.S2, green cells) compared to the average number of UMIs in the organ considered (Table.S2, black cells). Accompanying this increase in transcript number per nucleus, we noticed an increase in the number of expressed genes per nucleus (Fig.4B). We assume that the higher transcriptional activity of these clusters results from events of endoreduplication that have been reported in the seed endosperm and the Rhizobia-infected cells of the nodule. Detecting a larger population of transcripts per cell supports the enhanced detection of expressed genes per nucleus [38, 39]. Our findings support a previous work that mentioned the impact of endoreduplication on gene activity at the single cell-type level [40].

**Figure 4.**
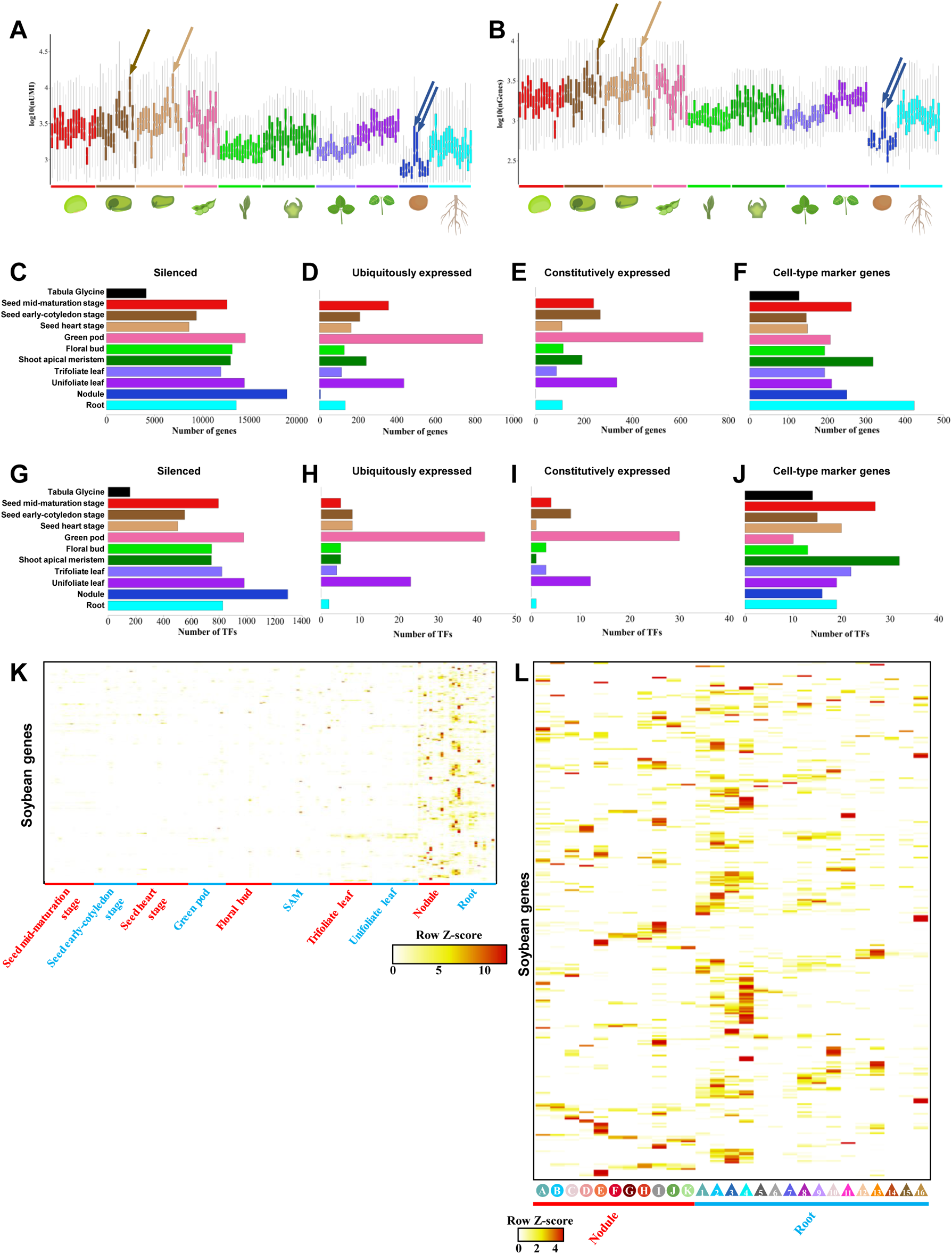
Differential analysis of gene activity between soybean cell types. Distribution of the number of UMIs (**A.**) and expressed genes per nucleus (**B.**) across the 157 cell clusters composing Tabula Glycine. Arrows highlight the cell types characterized by their endoreduplication and higher transcriptomic activities. **C-J.** Numbers of soybean silent (**C, G**), ubiquitously (**D, H**), constitutively expressed (**E, I**), and marker genes (**F, J**) for all the soybean genes (**C-F**) and the soybean TF genes (**G-J**) across 10 different soybean organs and in the entire Tabula Glycine, **K.** Heatmap of the transcriptional activity of 318 soybean genes considered as preferentially expressed in the soybean root and nodule (Moisseyev et al., 2020) (y-axis) for each of the 157 cell clusters (x axis). **L.** Magnification of this heat map to the 11 nodule and 16 root clusters.

To further reveal the unique transcriptomic activity of the 55,897 soybean protein-coding genes across soybean cell types, we conducted a comprehensive analysis of their expression (Table.S5). Considering the central role of TF-coding genes in controlling genetic programs, we also specifically analyzed the expression profile of 3,726 soybean TF genes (Table.S6). Accordingly, we categorized the soybean genes into 4 different groups based on their expression profile: 1) the silenced genes (i.e., not expressed in Tabula Glycine; Fig.4C and G); 2) the ubiquitously expressed genes (i.e., expressed in all the cell clusters with a transcriptomic activity detected in at least 20% of the nuclei composing the cluster; Fig.4D and H); 3) the constitutively expressed genes (i.e., subpopulation of ubiquitous genes characterized by less than 4-fold change of expression between the clusters where the gene is the most and less active; Fig.4E and I); and 4) the cell-type marker genes (i.e., genes expressed in at least 20% of the nuclei composing a cluster, with a minimum relative expression of 0.1 UMI, and with at least a 20-fold change of expression between the top two clusters where the gene is the most expressed; Fig.4F and J). Based on these parameters and considering Tabula Glycine as a whole, we identified 4,127 silenced genes (i.e., 7.4% of the 55,897 protein-coding genes; Fig.4C), including 157 TFs (i.e., 4.2% of the 3,747 predicted soybean TFs; Fig.4G). However, we did not identify any genes ubiquitously and, *a fortiori*, any constitutively expressed at the level of the entire soybean plant; likely a consequence of the stringency of our criteria and the partial coverage of the transcriptome of a few soybean clusters (Fig.1E). Therefore, to identify the most constitutively expressed genes in Tabula Glycine, we calculated the coefficient of variation of expression of all the soybean genes (Table.S7). Unexpectedly, among the 2,000 genes with the lowest coefficient of variation, we did not identify the popular *SKIP16*, *UKN1*, and *UKN2* reference genes [41–43] suggesting that these three commonly used reference genes at the organ level are differentially expressed at the cell type level. Mining Tabula Glycine, we found that the expression of these three genes is mostly restricted to a few cell types in specific organs. For instance, *SKIP16* is expressed in clusters #6 and 7 of the trifoliate and unifoliate leaves, *UKN1* is most expressed in clusters #12 and 14 of the floral bud while *UKN2* is most transcribed in clusters 2, 11, and 12 of the green pods (Fig.S9). Therefore, we propose a new set of 20 genes with the highest transcriptional stability across soybean cell types as new reference soybean genes (Fig.S10). We expect that their promoter sequences could be used to ubiquitously express transgenes in the soybean plant.

Our analysis also revealed 127 cell-type specific marker genes based on their preferential activity in one out of the 157 soybean clusters (Fig.4F). These genes are distributed across 19 cell types (Table.S5, “Whole plant markers” sheet) and represent another resource for the soybean community to preferentially express a transgene in one of these 19 soybean cell types. Among them, three cell types are over-represented: the nodule cluster D (i.e., sclereid layer; 26 marker genes), the root cluster #3 (i.e., root hair cell; 47 marker genes), and the root cluster #11 (i.e., endodermis; 20 marker genes) [22]; likely a reflection of the unique function of these cell types (Table.S5, “Whole plant markers” sheet). It is important to note that the representation of similar cell types isolated from different organs (e.g., phloem and xylem cells or the leaf mesophyll cells) might prevent the identification of additional valuable cell-type marker genes. Therefore, a more targeted exploration of Tabula Glycine might reveal additional interesting cell-type marker candidate genes. As another reflection of the uses of Tabula Glycine, we also reevaluated the expression pattern of 318 soybean genes identified as preferentially expressed in the soybean root system [44]. While we confirmed the preferential activity of these genes in the root and the nodule, we noticed their differential expression between clusters (Figs.4K and L). We also observed their strong expression in a limited number of cell types from the plant canopy suggesting the use of these putative “root” promoter sequences to express a transgene might also regulate the activity of genes in other cell types potentially leading to unexpected phenotypic responses. Therefore, Tabula Glycine gives a novel perspective on the activity of organ-specific genes by providing information on their cell-type specificity.

### The expression landscape of transcription factor genes to define cell type identity

Conceptual and experimental frameworks regarding the role of transcription factors in defining a cell type and controlling its biology have been developed in animal species. Notably, the combinatorial activation of TFs was described by Yao et al (2023) as a code to determine a cell type [45]. We assume that a unique combination of active TFs, which is synonymous with the core regulatory circuit in animal and human cells as defined by Almeida et al. (2021) and Arendt et al. (2016) [46, 47], would also apply to define the biology of plant cell types. Therefore, taking advantage of Tabula Glycine, we conducted a comprehensive analysis of the expression patterns of the soybean TFs.

First, to estimate the transcriptional diversity of the soybean 3,569 TFs [48] and its impact on organ and cell-type identities, we conducted an unbiased co-expression analysis of the soybean TFs across the 157 soybean clusters. Among these 3,569 TF genes, we analyzed the differential expression of the 2,335 TF genes narrowly expressed across the 157 cell clusters (i.e., Tau score > 0.9). We generated a cell lineage tree based on the co-expression pattern of TF genes (Fig.5A) and considered both the family membership of the 2,335 TF genes (Fig.5A, x-axis) and the cell-type annotation or organ of origin of the 157 cell clusters (Fig.5A, left and right y-axes, respectively).

**Figure 5.**
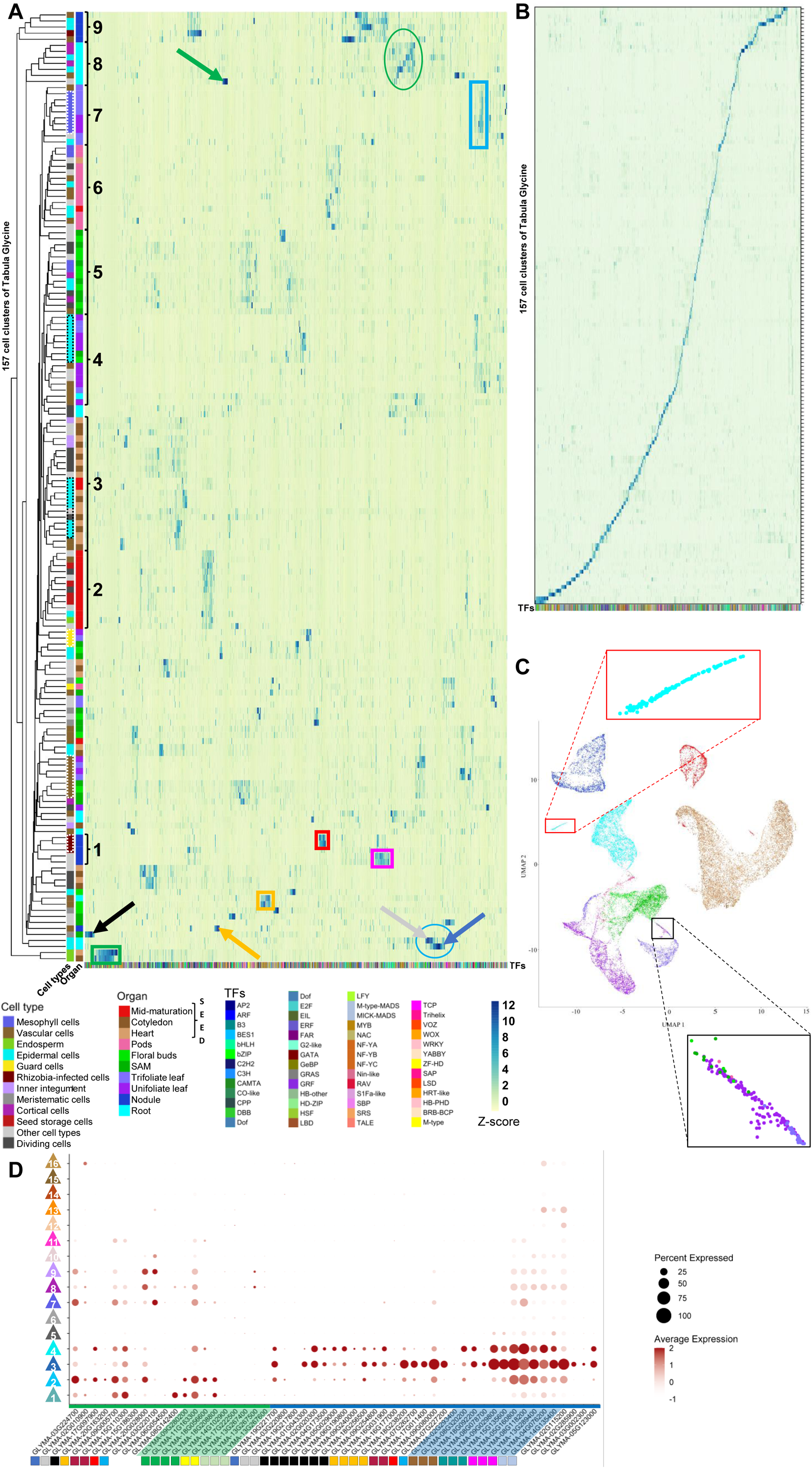
Co-expression analysis of the soybean TF genes. **A and B.** Heatmaps of co-expression of soybean TF genes (x-axis) across the 157 cell clusters of Tabula Glycine (y-axis). In **A.**, the 157 clusters are organized in a cell lineage tree (y-axis) based on the co-expression pattern of TF genes (x-axis). The functional annotation (e.g., epidermal, vascular, dividing, mesophyll, inner integument, cortical, storage, endosperm, rhizobia-infected, guard, meristematic cells, etc.) and organ of origin of the 157 cell clusters are highlighted with a color code (left and right y-axes, respectively). Branches of this trer are highlighted with black brackets and dashed boxes when considering the “organ of origin” and the “functional annotation” of the clusters (y-axes). Groups of co-expressed TF are highlighted with colored arrows and boxes. In **B.** the 157 clusters are rearranged while the order of the co-expressed TF genes is kept highlighting the sets of co-expressed TF specific to each cluster. **C.** UMAP plot of the epidermal cells of the soybean plant. Sub-populations of cells of interest are highlighted via zoom-in boxes. **D.** Dotplots of the expression pattern of co-expressed WRKY TFs and over-represented in two distinct groups of root clusters (i.e., the green and blue lines refer to the WRKYs identified among the set of co-expressed TF highlighted in the green and blue circles in Fig.5A, respectively). When identified, groups of duplicated WRKYs are highlighted by squares of the same color. WRKYs highlighted in the green and blue parallelograms are duplicated WRKYs that are strictly restricted to the same group of co-expressed TFs. The percentage of nuclei expressing the gene of interest (circle size) and the mean expression (circle color) of the genes are shown.

Our analyses reveal hundreds of groups of co-expressed TFs in one specific cluster [e.g., the root hair cells (blue arrow), the root cap (gray arrow), the root endoderm (green arrow), the sclereid layer of the nodule (orange arrow), or the meristematic cells of the shoot apex (black arrow)] or shared between several clusters with similar functional annotations [e.g., the infected (red box) and uninfected cells of the nodule (pink box), the mesophyll cells of the unifoliate and trifoliate leaves (blue box), the xylem cells (orange box), the endosperm of the developing seed (green box), or the lipid and protein storage cells in the developing seed (black box)] (Fig.5A, Table.S8 for details). Looking at the distribution of these populations of co-expressed TFs, we noticed that in most cases, they are unique to a cell cluster (Fig.5B). We assume that these cell type-specific co-expressed TFs likely synergistically activate and repress transcriptomic programs to regulate the biology of functionally specialized cell types. Hence, the combinatorial activation of TFs could serve as a code to control the biology of plant cells and, *a fortiori*, as a molecular marker of plant cell identity. Identifying the same sets of co-expressed TFs between cell clusters from different organs but with similar functional annotation (Fig.5A, colored boxes) also demonstrates the quality of the transcriptional information shared in Tabula Glycine.

To better assess the biological impact of the combinatorial activation of TFs of the soybean cell type identity, we analyzed the distribution of the functional annotation and organ of origin of the 157 Tabula Glycine clusters in the cell lineage tree (Fig.5A, y-axes). We found that the tree is mostly organized based on the organ of origin of the clusters (black brackets labeled 1 to 9 on the right y-axis legend, Fig.5A). This observation suggests that the cells’ TF transcriptional patterns are strongly associated with the organ’s function. To a lesser extent, branches of this tree are also organized based on the functional annotations of the cell clusters (white and black dashed boxes on the left y-axis legend, Fig.5A). We noticed that the latter include cell types characterized by highly specialized biological function such as the rhizobia infected cells of the nodule, the vascular cells of the canopy, the guard cells, or the mesophyll cells (Fig.5A, left y-axis).

In contrast, the epidermal cells of various soybean organs are distributed along the entire cell lineage tree (cyan color in the “cell types” y-axis) with two areas of concentration that include the epidermal cell of the developing seed and the leaf (i.e., see the black dashed boxes on the left y-axis). This result suggests that the epidermal cells, despite their common functional annotation, also have organ-specific transcriptomic signatures. To verify this hypothesis, we generated an integrated UMAP based on the transcriptome of all epidermal-labeled nuclei (Fig.5C) and observed that the epidermal nuclei were clustered based on their organ origin further supporting the organ’s primary role in the transcriptome of plant cells. Notably, we noticed three main clusters of epidermal cells from the nodule and the root (Fig.5C, blue and dark blue clusters), the canopy (Fig.5C, green and purple clusters), and the soybean seed (Fig.5C, red and brown clusters). Among these clusters, we noticed two interesting sub-populations. The first sub-population is composed of nuclei isolated from different organs of the canopy (Fig.5C, black box) suggesting a strong conservation of the transcriptome between these epidermal cells. The second subpopulation is composed of a subset of the root epidermal cells (Fig.5C; red box). Based on the expression of marker genes (Figs.S11; [22]), we functionally annotated the first and second sub-populations as the guard cells and the root hair cells, respectively. Our data suggest that the unique function of these two cell types is reflected in their unique transcriptomic profiles conserved across different organs in the case of the guard cells. This statement is further supported by the identity of the co-expressed TFs. When considering the set of 70 TFs co-expressed in the root hairs (Fig.5A; blue arrow; Table.S8, bold characters), we identified 16 bHLH TFs that include seven soybean genes (i.e., Glyma.04G045300, Glyma.10G257400, Glyma.10G257500, Glyma.14G088400, Glyma.17G236100, Glyma.20G133600, and Glyma.20G133700) orthologous to *AtRSL2* (*Root Hair Defective 6-Like2*) *and AtRSL4*, genes promoting root hair cell growth [49, 50]. Two other *bHLH* genes, Glyma.17G075200 and Glyma.02G202600, are orthologous to the *AtLRL1/2/3* (*Lotus japonicus* Roothairless1-LIKE 1/2/3) genes, genes controlling root hair development. As a note, *AtLRL3* has been characterized as a direct target of RHD6 TF [51]. In the guard cells, among a group of 30 co-expressed TFs in the uni-and trifoliate leaves, we identified two soybean genes (i.e., Glyma.03G006600 and Glyma.19G119300) orthologous to *AtMYB60*, a central player controlling stomata opening [52, 53]. Considering that the biological samples were collected during daytime, it is not surprising to identify these *AtMYB6*0 orthologs among the set of leaf guard cell-specific co-expressed TF genes. Taken together, our results support the concept that a set of co-expressed TFs can be used as a molecular marker of organ and cell type identity especially when considering specialized cell types. This suggests that shared cell-type-specific co-expressed TFs between different organs play a fundamental role in controlling the basal function of a plant cell type like the core regulatory circuit in animal cells.

Our TF co-expression analysis also allowed us to estimate the distribution of TFs according to their family membership in groups of cell-type-specific co-expressed TFs (Fig.5A, x-axis). We hypothesize that the over-representation of TF families in a specific group would reflect their central regulatory role in controlling the biology of a cell type. Accordingly, we compared the representation of TF families in major groups of co-expressed TFs versus their representation in the soybean genome. Several TF families were over-represented in a few groups of co-expressed TFs. For instance, *ZF-HD* and *M-type-MADS* were overrepresented among the TFs co-expressed in the endosperm of developing seeds (adjusted p-value = 1.12e^-13^ and 4.88e^-08^, respectively). This observation concurs with previous studies in *Arabidopsis thaliana*, which showed that the overexpression of *AtZHD1* positively affects seed size [54] and the high transcriptional activity of *M-type-MADS* genes in the developing endosperm of the Arabidopsis seed and, more broadly, in the endosperm of many flowering plants [55]. For instance, among these MADS TFs, Glyma.18G052800 is orthologous to *AtAGL80*, a gene controlling endosperm development [56], and *AtAGL36*, a gene characterized by its imprinting to maintain its activity in the Arabidopsis seed endosperm [57]. Other genes include Glyma.20G136500, an ortholog to *AtAGL62* that has been reported to be essential for the suppression of cellularization and the promotion of cell proliferation during endosperm development [58], and Glyma.13G086400, an ortholog of *AtAGL66* and *AtAGL67* that regulate seed germination [59]. In the nodule cells entering in senescence, we observed the over-representation of *ERF* and *C2H2* TFs (adjusted p-value = 7.87e^-19^ and 3.01e^-08^, respectively). Previous studies showed that the expression of several soybean *ERF* and *C2H2* genes such as *RSD* (*REGULATOR OF SYMBIOSOME DIFFERENTIATION*) is critical to trigger the senescence of the legume nodule [60–62]. *NAC* and *MYB* TFs are also over-represented but in the set of TFs co-expressed in the xylem (adjusted p-value = 9.12e^-10^ and 9.12e^-10^, respectively). Such an observation is well supported by the functional characterization of several *VND* genes (*VASCULAR NAC DOMAIN*) that regulate the deposition of cell walls in the plant vascular tissues and control the differentiation of xylem vessel elements [63]. Similarly, several *MYB* TFs control the deposition of cell wall components such as lignin in the plant vascular tissues including the xylem [64–66].

Two other groups of co-expressed TFs are over-represented in *WRKY* TFs (Fig.5A, blue and green circles, adjusted p-value = 1.24e^-07^ and 2.54e^-28^, respectively). The first group (i.e., group A; Fig.5D, green line) is composed of 20 *WRKY* genes broadly co-expressed in the epidermal and cortical, and to a lesser extent in the phloem cells of the root, while the second group (i.e., group B; Fig.5D, blue line) is composed of 33 *WRKY* genes almost specifically co-expressed in the root epidermal cells including the root hair cells. Hypothesizing that several of the genes in and between groups might be evolutionarily related as the result of small-scale and whole genome duplications, we conducted a comprehensive analysis of their evolutionary relationship and evaluated their level of transcriptional conservation and divergence (see Methods for details). Among these 53 *WRKY* TFs, only 7 genes do not share an evolutionary relationship (Fig.5D, unlabeled genes). The remaining 46 TFs are distributed into 14 sets of duplicated genes (Fig.5D, colored squares). Among them, 9 and 11 WRKYs from groups A and B are duplicated in their respective groups suggesting the conservation of their expression profile and function upon duplication (Fig.5D, genes highlighted with a green and blue background, respectively). The remaining 8 and 18 WRKY genes from groups A and B, respectively belong to 7 sets of duplicated genes. The partitioning of these duplicated *WRKYs* in two different groups of co-expressed genes suggests occurrences of sub-or neo-functionalization upon duplication. Altogether, our co-expression analysis reveals groups of plant TFs that likely cooperate to control specific genetic programs that are critical in regulating cell-type function. Our analysis also highlights the evolutionary relationship of TFs in groups of co-expressed TFs as another avenue to estimate their functional redundancy, and the sub-or neo-functionalization between the members of TF families. We propose that such analysis would support the establishment of new functional genomic strategies where the knock-out or overexpression of multiple genes would lead to more drastic and informative phenotypes.

### A cellular transcriptome most reflects the biological function of a cell

We extended our analysis to the entire transcriptome of the soybean cell types by hypothesizing that, among 157 Tabula Glycine clusters, those with the same cell type annotations might share, at least to some extent, a core transcriptome independent of their organ of origin. Upon performing a dimensionality reduction by UMAP embedding, we identified 29 clusters (Fig.6A, Video.S1), drastically reducing the complexity of the 157 soybean clusters previously mentioned (Fig.1) and suggesting that different populations of nuclei share common sets of transcriptomic programs (Table.S9). Among them, clusters #1, 4, 5, and to a lesser extent, 13 and 20 show the most distinct transcriptomic signatures according to our PCA analysis (Fig.6B). Quantitatively, cluster #25 is characterized by the largest population of UMIs and expressed genes compared to other clusters (Fig.6C and D). The analysis of the nuclei composing this cluster revealed that cluster #25 of the integrated seed UMAP is mostly composed of seed endosperm cells.

**Figure 6.**
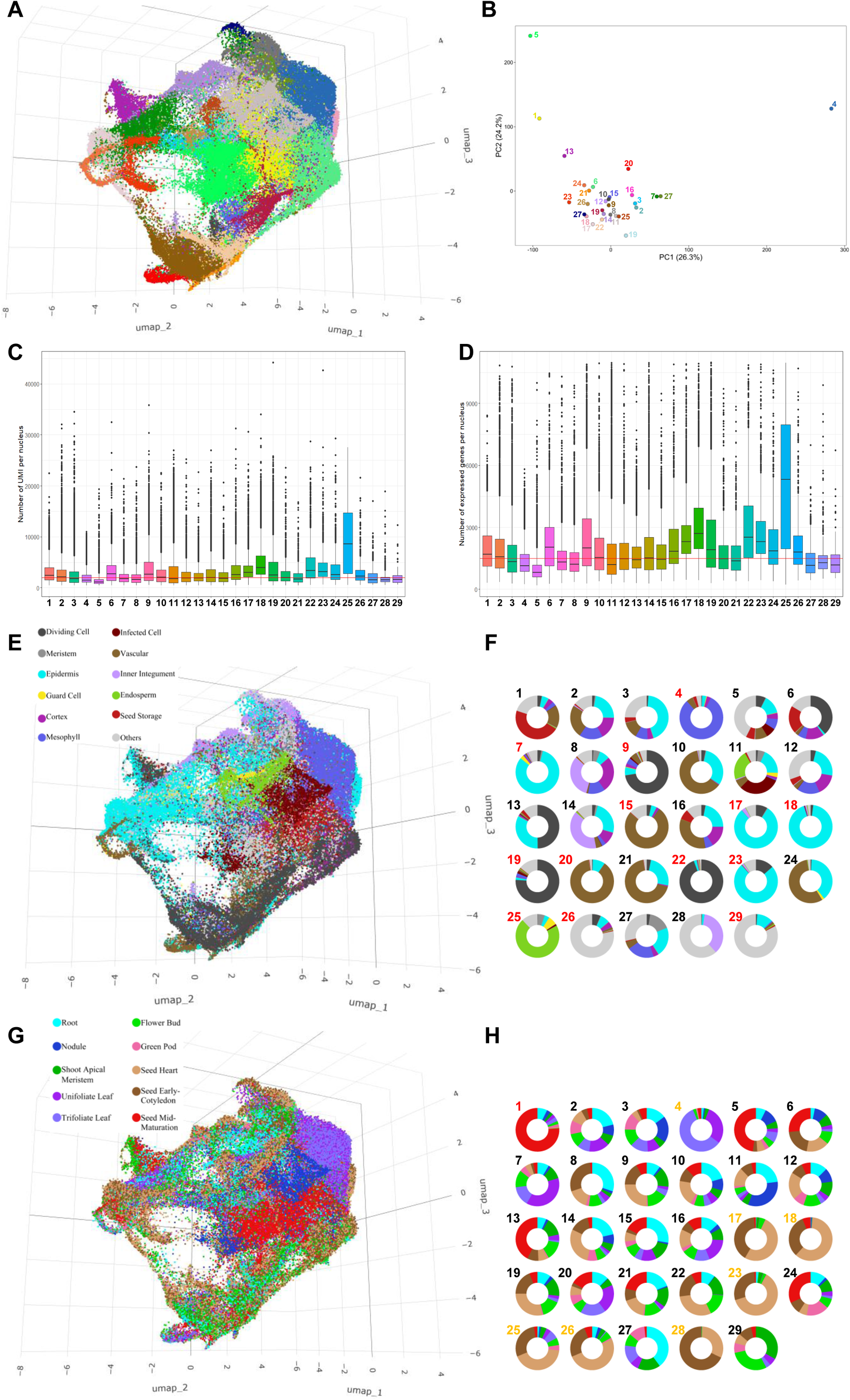
Integrated Tabula Glycine UMAP reveals the preferential clustering of the soybean nuclei based on their functional annotation. A. Integrated 3D UMAP plot according to the transcriptomic profile of 116,525 nuclei isolated from 10 different soybean organs. This clustering led to the 29 clusters. B. Principal component analysis of the transcriptome of the 29 clusters of the integrated Tabula Glycine UMAP. C and D. Number of Unique Molecular Identifiers (UMI; C) and expressed genes per nucleus (D) in each cluster composing Integrated Tabula Glycine UMAP. E to H. Labels of the integrated Tabula Glycine 3D UMAPs (E and G) and distributions of the nuclei in each cluster (F and H) are given according to the functional annotation of the nuclei (E and F) or their organ of origin (G and H). Clusters represented by over >70% of nuclei sharing the same cell type identity (F) or organ of origin (H) are highlighted in red. Due to the cellular redundancy existing between the unifoliate and trifoliate leaves, and the seeds at the heart and cotyledon stages, we highlight in orange the clusters composed by >70% of nuclei sharing the same cell type identity when conjointly considering unifoliate and trifoliate leaves, and seeds at the heart and cotyledon stages.

To annotate these 29 integrated clusters, we analyzed the 10x barcodes of each nucleus. Then, similarly to our analysis of the co-expression of TFs (Fig.5), we considered both the cell-type identity (Fig.6E and F) and the organ of origin (Fig.6G and H) to annotate each of the 29 clusters. We observed the overrepresentation of nuclei (>70% in a cluster) sharing the same cell type identity in 13 out of the 29 clusters (Fig.6F, clusters highlighted in red). Among them, the dividing, epidermal, mesophyll, and vascular cells from various organs are often clustered together, likely a reflection of their functional specialization. Contrastingly, only 1 cluster is composed of over 70% of nuclei from the same organ (i.e., the seed at the mid-maturation stage Fig.6H, cluster number highlighted in red). This number increases to 8 clusters when conjointly considering unifoliate and trifoliate leaves, and seeds at the heart and cotyledon stages (Fig.6H, cluster numbers highlighted in orange).

The impact of the transcriptome on cell function was further revealed upon integration of these two analyses. For instance, when considering clusters #7, 9, 15, 19, 20, 22, and 29 that are mostly composed of nuclei sharing the same cell type annotation (Fig.6F), their organ of origin is quite diverse as reflected by the mosaic pattern of the “organ” donut plots for these clusters (Fig.6H). This result suggests that the transcriptomic profile of these cell types shares similar properties independently of their organ of origin. In other words, these cells share a common core transcriptome, likely a signature of the biological function.

To further explore the biological significance of the co-clustering of different cell types from various soybean organs based on their transcriptional profile, we identified the GO terms most enriched for each of the 29 clusters (Fig.S12). The integration of this analysis with the allocation of cell-type and organ identity (Fig.6F and H) allowed us to functionally annotate most of the 29 clusters. Among the clusters characterized by their cellular function, clusters #15 and 21 are composed of the xylem and phloem cells from different soybean organs, respectively. When considering the soybean leaf, cluster #7 is mostly composed of leaf epidermal cells engaged in cutin and wax biosynthesis. Cluster #4 is also a leaf-specific cluster, but of the mesophyll based on its enrichment in photosynthetic GO terms. From the seed, clusters #1, 5, and 13, that are most represented by nuclei from the seed mid-maturation stage nuclei, are characterized by the activity of genes controlling fatty acid biosynthesis, the storage of fatty acids and proteins, and seed maturation. Clusters #9, 19, and 22 are the active mitotic and meiotic cell clusters composed of nuclei isolated from developing seed, shoot apical meristem, and floral buds. Based on the over-representation of cells from the seed engaged in cell division and the enrichment of GO term controlling cell polarity, we assume that cluster #6 is composed of dividing cells including embryonic cells.

Other clusters are characterized by their molecular process. Noticeably, cluster #25 is unique based on its high transcriptional activity (Fig.6C and D) further supported by several GO terms associated with gene regulation notably at the epigenetic level. Considering that this cluster is composed of nuclei from developing seeds, we assume that this intense transcriptional activity is needed to support the differentiation processes occurring in developing seeds. While clusters #3 and 8 did not show any specific cell type-or organ-identity enrichment, the GO analysis revealed the presence of cells engaged in the cell death program, stresses, and microbial symbiosis for cluster #3, and in amino acid catabolic processes for clusters #8.

Finally, another population of clusters is characterized by their metabolomic and catabolic activities. For instance, clusters #18 and 28 are characterized by amino acid catabolism, specifically in the developing seeds. Cluster #11, a cluster enriched in nodule and root nuclei, shows a strong regulation in inosine monophosphate biosynthesis, a metabolomic pathway used by the soybean plant to synthesize purine to assimilate the atmospheric nitrogen fixed by its symbiotic bacterial partner. Clusters #12 and 24 are characterized by cell wall biosynthetic processes. Another example is cluster #29 that is most represented by floral bud and shoot apical meristem nuclei and characterized by the biosynthesis of GDP-mannose, a structural carbohydrate, ascorbic acid, a regulator of cell division, and lactone. Several clusters are also specialized in the synthesis of specialized metabolites. For instance, cluster #16, an epidermis cell cluster, is involved in the biosynthesis of terpenoids and flavonoids. These compounds play a central role in the interaction between plants and their biotic environment. Another class of flavonoid compounds, proanthocyanidins, is synthesized by the cells composing clusters #17 and 23. This family of metabolites has antibacterial and antioxidant properties [67, 68]. These clusters, as well as cluster 14, are also involved in the synthesis of phenylpropanoids that also trigger plant defense responses.

## Discussion

Recent advances in single-cell genomics methods have enabled the generation of single-cell transcriptome atlases of different animal models such as *C. elegans* [69], mouse [70], human [71], and Drosophila [72], providing strong resources for research at cell resolution. In plants, single-cell transcriptome atlases are often limited to one organ [73, 74]. Here, we present Tabula Glycine, the first crop single-cell transcriptome atlas. This resource reveals new genomic properties of the soybean plant, a major source of oil and proteins for human and animal consumption, including cell-type resolution transcriptomic signatures that will enable enhanced functional genomic and synthetic biology strategies, the combinatorial expression of TFs as a major contributor of cell and organ identity, and a comprehensive analysis of the lineage of the soybean cells based on their transcriptomic profiles.

### A new resource for the plant science community

We expect that Tabula Glycine will quickly become a resource for the soybean and plant science communities. The deep coverage of the transcriptome of most of the cell types composing Tabula Glycine (Fig.1F and evidence from the quantification of the expression of almost all the soybean TF genes, genes known to be expressed at low levels) and the functional annotation of many of the soybean clusters strongly support the quality of Tabula Glycine. Notably, considering soybean as a non-model species, the functional annotation of each cell type composing these organs was a challenging task (Figs.2 and 3). However, the use of different strategies such as Molecular Cartography^TM^ to annotate the soybean root and nodule cell types [22], and the analysis of the expression of soybean orthologs to *A. thaliana* and *M. truncatula* marker genes led to the assignment of more than 80% of the Tabula Glycine cell clusters. Considering the broad range of organs analyzed, the large biological context of Tabula Glycine offers an opportunity to better understand the genetic programs governing the biology of each soybean cell type. For instance, we reveal that the expression patterns of the soybean TFs are sufficient to define most, if not all, the soybean cell types through the specific combination of co-regulated TFs (Fig.5B).

Hence, in addition to precisely establishing the transcriptional profile of candidate genes for functional genomic studies, Tabula Glycine will enable discoveries and applications that will serve soybean genomicists, cell biologists, evolutionary biologists, and the plant science community. For instance, and as described in this manuscript, we refined the set of reference and cell-type marker genes at the single-cell resolution. This information will impact the decision process of scientists when selecting promoters to drive the expression of transgenes in the most biologically relevant cell types. It will also help to estimate the functional redundancies-notably those resulting from small-scale and whole-genome duplications-and cooperation between genes in the same cell to support the establishment of functional genomic strategies that will drive the most significant phenotypic changes. For instance, our analysis of the co-expression of two different populations of WRKY TFs activated in two different sets of root cells revealed likely occurrences of both functional redundancy and neo-or sub-functionalization. Considering the evolutionary features of the soybean genome (i.e., soybean is an allotetraploid that results from two rounds of whole genome duplications [7]), Tabula Glycine will enhance our understanding of the evolution of the expression and co-expression of the soybean genes. We also expect Tabula Glycine to validate and enhance predicted gene networks providing an unprecedented resolution. As a demonstration of this potential, we recently created a gene co-expression network of the soybean nodule cells infected by their nitrogen-fixing symbiotic symbiont, *B. diazoefficens* [22]. Therefore, we expect that Tabula Glycine will support the design of high-resolution functional genomic approaches to enhance specific soybean traits such as protein and oil yield and composition, biological nitrogen fixation to support sustainable agricultural practices, plant organ development, etc.

### Enhancing current transcriptomic resources to support soybean biology

Tabula Glycine is the first high-resolution atlas developed from multiple organs for a major crop species. The broad spectrum of organs represented in Tabula Glycine (i.e., from the mature nodule infected by the nitrogen-fixing bacteria *B. diazoefficiens* to the seed at the mid-maturation stage) gives a unique perspective on the differential use of the genome by the 157 cell clusters analyzed in this study. This statement is supported by the analysis of the co-expression patterns of TF genes in 10 different organs that revealed cell-type-specific sets of co-expressed TFs. We assume these TFs act together to drive important genetic programs in various cell types. Our integrated analysis of the entire Tabula Glycine reveals that the same cell types isolated from different organs share, at least to some extent, a common core transcriptome (Fig.6). Hence, the characterization of the antagonistic and synergistic roles of TF genes and the identification of the genes under their control in the context of the epigenetic environment would further support our understanding of the regulatory mechanisms of plant gene expression. It also offers an opportunity to modify and control the biology of specific plant cell types.

### Perspectives

Tabula Glycine is a first step forward in our understanding of the differential use of the soybean genome by individual plant cells. Revealing the dynamic changes in gene expression occurring in each cell composing the plant during its development, in response to individual or combinatorial environmental stresses, and/or upon genetic perturbations is the logical next step. Accessing such information will allow the capture of transcriptomic changes, and the generation of cell-type-specific transcriptomic trajectories. Another avenue of expansion for Tabula Glycine consists in the integration of several single cell resolution-omics datasets (e.g., RNA and ATAC-seq) in order to reveal the mechanisms controlling gene activity. The use of high-throughput single-cell-omics technologies will be key to generate the depth of information needed to gain a systems-level understanding of the soybean plant at the single-cell level.

## Supporting information

Other supplemental figures

## Acknowledgments

We thank Ms. Rosa Angelica Bedolla-Gaxiola for her artistic work (Figure 1A). We thank Dirk Anderson of the Flow Cytometry Core Research Facility at the University of Nebraska-Lincoln Center for Biotechnology for helping with nuclei sorting. We also thank Dr. Jeffrey J. Doyle from Cornell University for his constructive discussions. This research was funded by the Nebraska Soybean Board, the NSF awards #2127485 and #1854326, and the Nebraska Research Initiative. OCM and IPS2 benefit from the support of Saclay Plant Sciences-SPS (ANR-17-EUR-0007).

## Author contribution

ML supervised the work and ensured that authorship was granted appropriately to contributors; STh generated single-cell RNA-seq libraries; SACP processed the raw single-nucleus RNA-seq data; OCM generated the prediction of transcriptomic coverage; SACP and STh generated the genome mapping; CMS, SACP, ADF, JC performed the evolutionary analysis to assign orthologous genes; SACP, STh, JMP, IM, HX, STe, HN, RBG, JJH, and ML performed functional annotation of the cell types, QY and PM created web-based tools for data visualization. SACP, STh, SA, STe, IM, HX, EJA, OM, RBG, JJH, and ML analyzed the results; SACP and ML wrote the manuscript with the contribution of all authors.

## STAR Methods

### Plant material

Soybean (*Glycine max Williams* 82) seedlings were sterilized as previously described [75]. The true leaf samples were collected from 2-week-old seedlings grown in soil in growth chamber conditions. The rest of the samples were collected from plants grown in soil (Green mix, 10% Canadian peat, 40% coarse vermiculite, 15% Masonry sand, and 5% screened topsoil) in the greenhouse (24-27°C/ 18-21°C). Fully developed trifoliate leaves (without petiole) and shoot apical meristem were harvested from 2-month-old plants. Flower bud samples were collected from R1 plants before the emergence of the petals. The heart, early-cotyledon, and mid-maturation seed samples are pooled seeds measuring 2, 5, and 10mm. Pods without their 10 mm seeds were collected from the green pod sample.

### Nuclei isolation, sNucRNA-seq library preparation, and sequencing

For nuclei isolation, soybean tissue samples were chopped and passed through 30 and 40 µm cell strainers as previously described [76]. The filtered nuclei were purified by cell sorting using FACS Aria II™ 603 cell sorter (BD Biosciences), centrifuged, and resuspended in a solution of PBS and BSA 0.5%. The sNucRNA-seq libraries were constructed following the Chromium™ Single Cell 3’ Library & Gel Bead Kit v3.1 protocol (10x Genomics) with a targeted recovery of 5,000 nuclei. The sequencing of single-and paired-indexed, paired-end libraries was performed on an Illumina™ NovaSeq 6000 platform according to the 10x Genomics recommendations. See Table.S10 for detailed information regarding these libraries.

### SNucRNA-seq data pre-processing, integration, and clustering

Each sNucRNA-seq library was processed individually using the 10x Genomics Cell Ranger software v6.1.1.0 for the demultiplexing and for the alignment against the soybean reference genome from Ensembl Plants database (i.e., Glycine_max_v2.1.52; http://ftp.ensemblgenomes.org/pub/plants/release-52/fasta/glycine_max/). The background contamination was subtracted after the alignment of the reads using SoupX [77], and doublets were filtered out using the DoubletDetection prediction method^70^. Finally, we applied a minimum threshold of 500 UMIs to remove the nuclei with lower transcriptional content. Upon normalization, the integration anchors were defined for the integrated replicates of each sample using the tool Seurat V4^71^. The dimensional reduction for the complexity of the data was performed using the UMAP method, upon selecting the top 2000 variable genes for the clustering by using the FindClusters method from Seurat V4.

### UMAP visualization

For visualization, all the sNucRNA-seq libraries for each sample were combined using the Cell Ranger aggr function from 10X Genomics and we used Loupe software from 10X Genomics to visualize the integrations.

### Comparison of soybean sNucRNA-seq and bulk RNA-seq

To evaluate the depth and sensitivity of the soybean unifoliate leaf, shoot apical meristem, pod, and floral bud single nuclei transcriptome atlases, we compared our pseudo-bulk sNucRNA-seq datasets with previously published bulk RNA-seq datasets [9]. Using the database of the legume information system (LIS), we extracted bulk expression datasets (2022/11/14; https://data.legumeinfo.org/Glycine/max/expression/Wm82.gnm2.ann1.expr.Wm82.Libault_Far mer_2010/; identifiers # SRR037381, SRR037382, SRR037383, and SRR037384 before comparing the number of expressed genes between bulk and pseudo-bulk RNA-seq libraries.

### Transcriptomic coverage in a cluster and mathematical framework

Given any cell in a cluster, we compute for each gene *i* the fraction *x_i_* it contributes to the cell’s UMI transcripts. We then average these *x_i_* over cells in the cluster, giving a pseudobulk proportion *p_i_* for each gene. To define an associated diversity index, we follow a common choice as used in ecology [78, 79], namely the Shannon entropy of the *p_i_*. Similarly, the *effective* number *E_G_* of genes expressed in the cluster can be taken to be the exponential of that quantity. Note that *E_G_* will be small if the number of cells *M* in the cluster is low whereas *E_G_* will go to some limit if *M* is increased indefinitely. That limit corresponds to a perfect specification of the transcriptomic profile, so in practice it is important to find out how far one is from that limit. Based on subsampling, we find that *E_G_* is very well fit by a Pareto distribution:

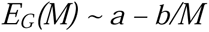

We thus define the *transcriptomic coverage* for the considered cluster with its limited number of cells as the measured value of *E_G_* divided by the (mathematically determined) value of *E_G_(M)* were *M* to become arbitrarily large.

This measure of transcriptomic coverage tells us whether the number of cells in the cluster is sufficient to approximate the “exact” transcriptomic profile. When the coverage is close to 1, adding more cells will not change the transcriptomic profile whereas when the coverage is low it is advisable to add more cells to have a better precision on that profile.

### TF gene co-expression analysis

To find the TF gene expression, we used the pseudo-bulk transcriptomic data for each cell cluster of the soybean cell atlas. The entire dataset was normalized using a Z-score test, and the most significant TF pairs were selected by applying a Tau test (Tau>0.9).

### Defining Soybean paralogs and *Arabidopsis thaliana* Orthologs

Five comparative genomics methods were used to determine orthologs between *Glycine max* and *Arabidopsis thaliana*. MCScanX was run using its MCScanX_h homology mode which relies on *a priori* gene homology as opposed to BLAST algorithm hits, although both organize homologous genes into syntenic blocks for ortholog identification [80]. Homology was provided by the protein family databases of the Legume Information Service [81]. The remaining four methods (orthologous gene families, best search hits, TribeMCL tree-based gene families, and ortholog through colinear regions) are included in the orthology databases of plant genomics platform PLAZA 5.0 [82–84]. These were used to validate the predicted orthologs from MCScanX, meaning an orthologous gene pair was included if it was supported by MCScanX and at least one of the four tools from PLAZA. Paralogs in the soybean genome were assigned to duplication mechanism as follows: Syntenic paralogs (as identified by MCScanX) were assumed to be the result of whole genome duplication (WGD). These were assigned to the papilionoid WGD if the median Ks for all gene pairs in the syntenic block was greater than 0.40 and to the Glycine-specific WGD if the median Ks was less than or equal to 0.40. Non-syntenic paralogs were considered tandem duplicates if they are on the same chromosome with no non-paralogous genes between them, proximal duplicates if they are on the same chromosome with 1-10 non-paralogous genes between them, and transposed duplicates if there are more than 10 non-paralogous duplicates between them or they are on different chromosomes.

## Data availability

The sNucRNA-seq data generated in this study have been deposited with the National Center for Biotechnology Information (NCBI) bioproject numbers PRJNA938968, PRJNA983388, and the GEO numbers GSE226149, GSE234864.

Any additional information required to reanalyze the data reported in this paper must be requested from the lead contact.

