## Supplementary material for "Tabula Glycine: The whole-soybean single-cell resolution transcriptome atlas": FigS1

### Slide 1
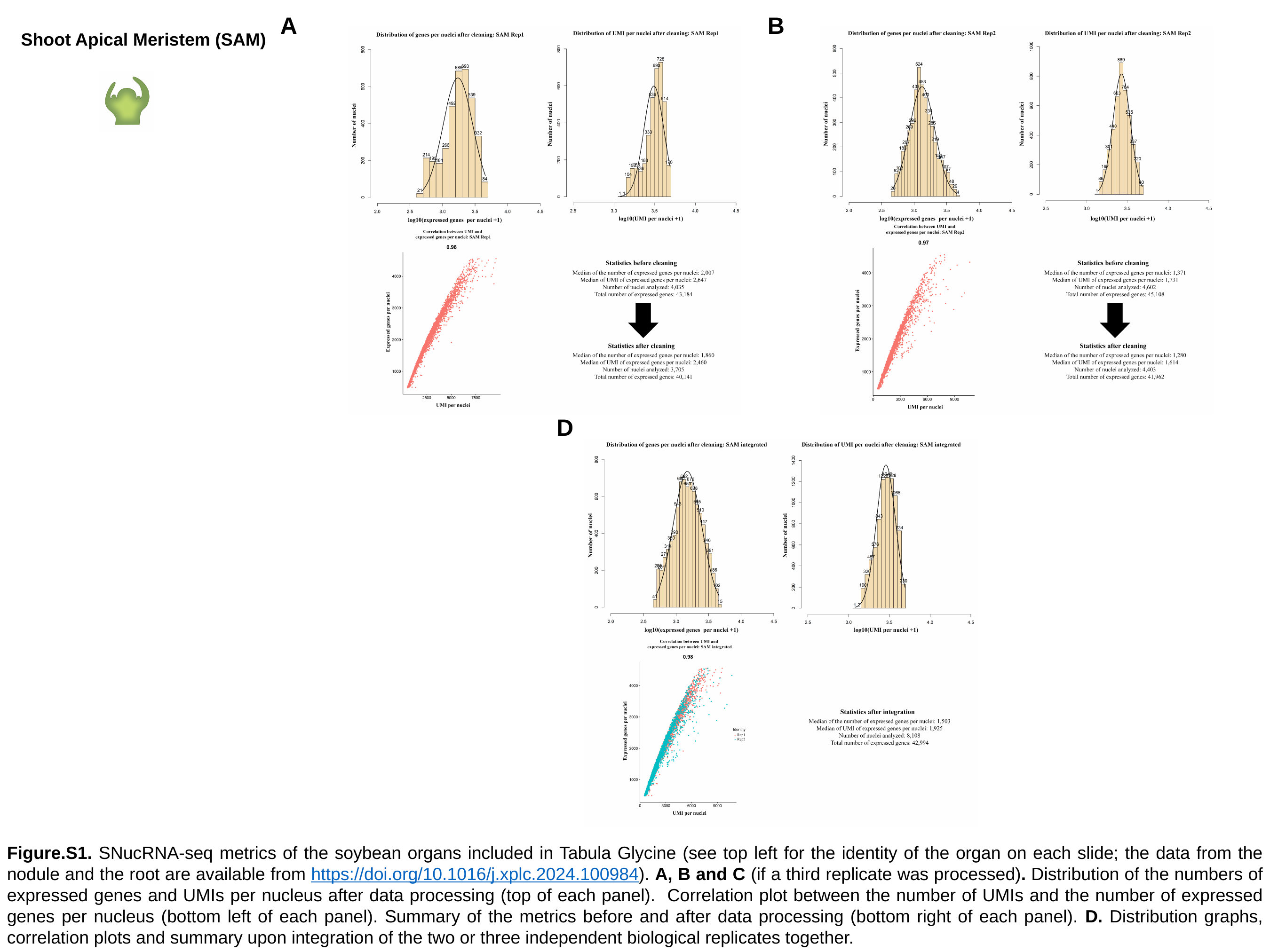

A
B
D
Shoot Apical Meristem (SAM)
Figure.S1. SNucRNA-seq metrics of the soybean organs included in Tabula Glycine (see top left for the identity of the organ on each slide; the data from the nodule and the root are available from https://doi.org/10.1016/j.xplc.2024.100984). A, B and C (if a third replicate was processed). Distribution of the numbers of expressed genes and UMIs per nucleus after data processing (top of each panel). Correlation plot between the number of UMIs and the number of expressed genes per nucleus (bottom left of each panel). Summary of the metrics before and after data processing (bottom right of each panel). D. Distribution graphs, correlation plots and summary upon integration of the two or three independent biological replicates together.

### Slide 2
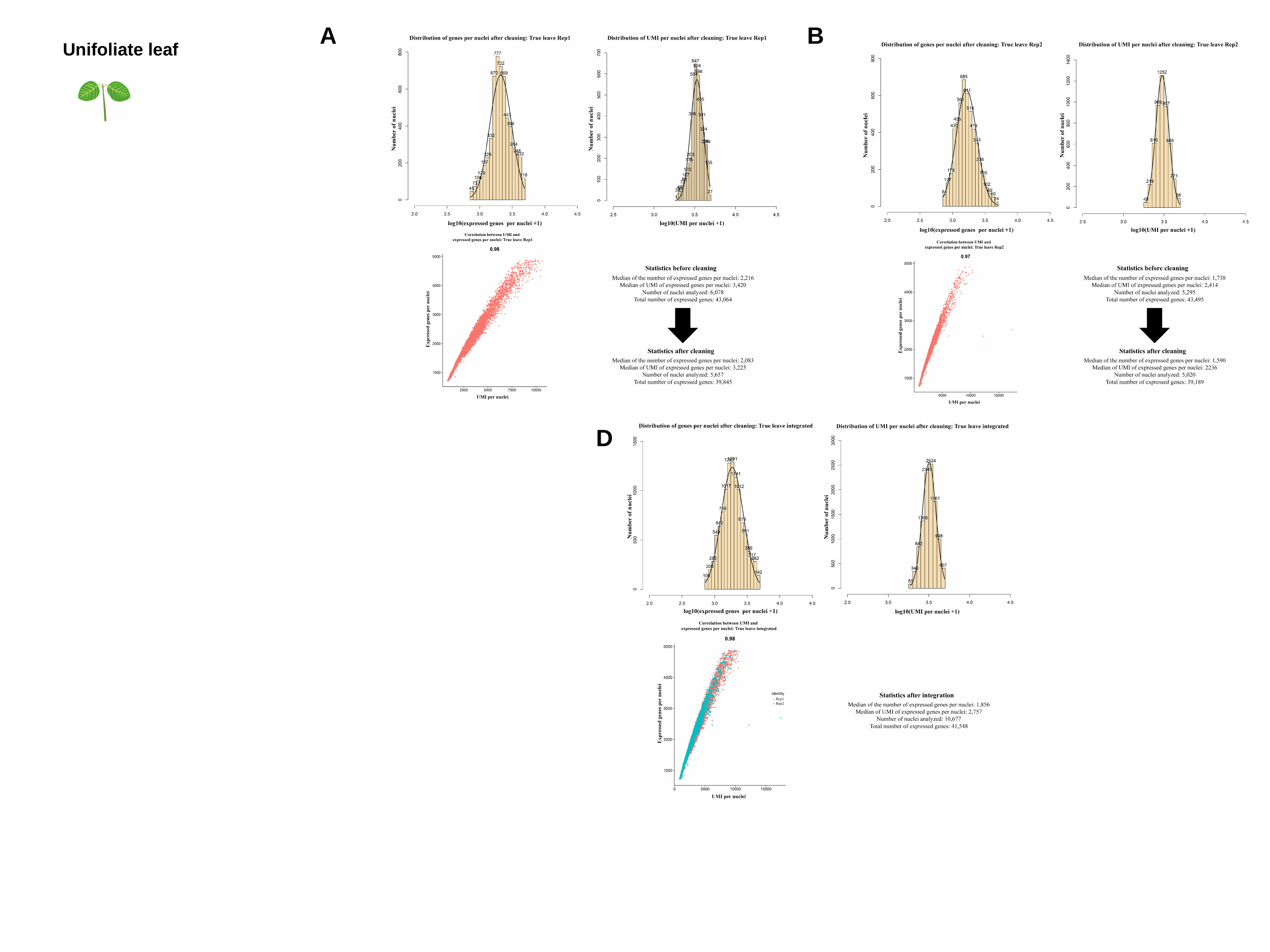

A
B
D
Unifoliate leaf

### Slide 3
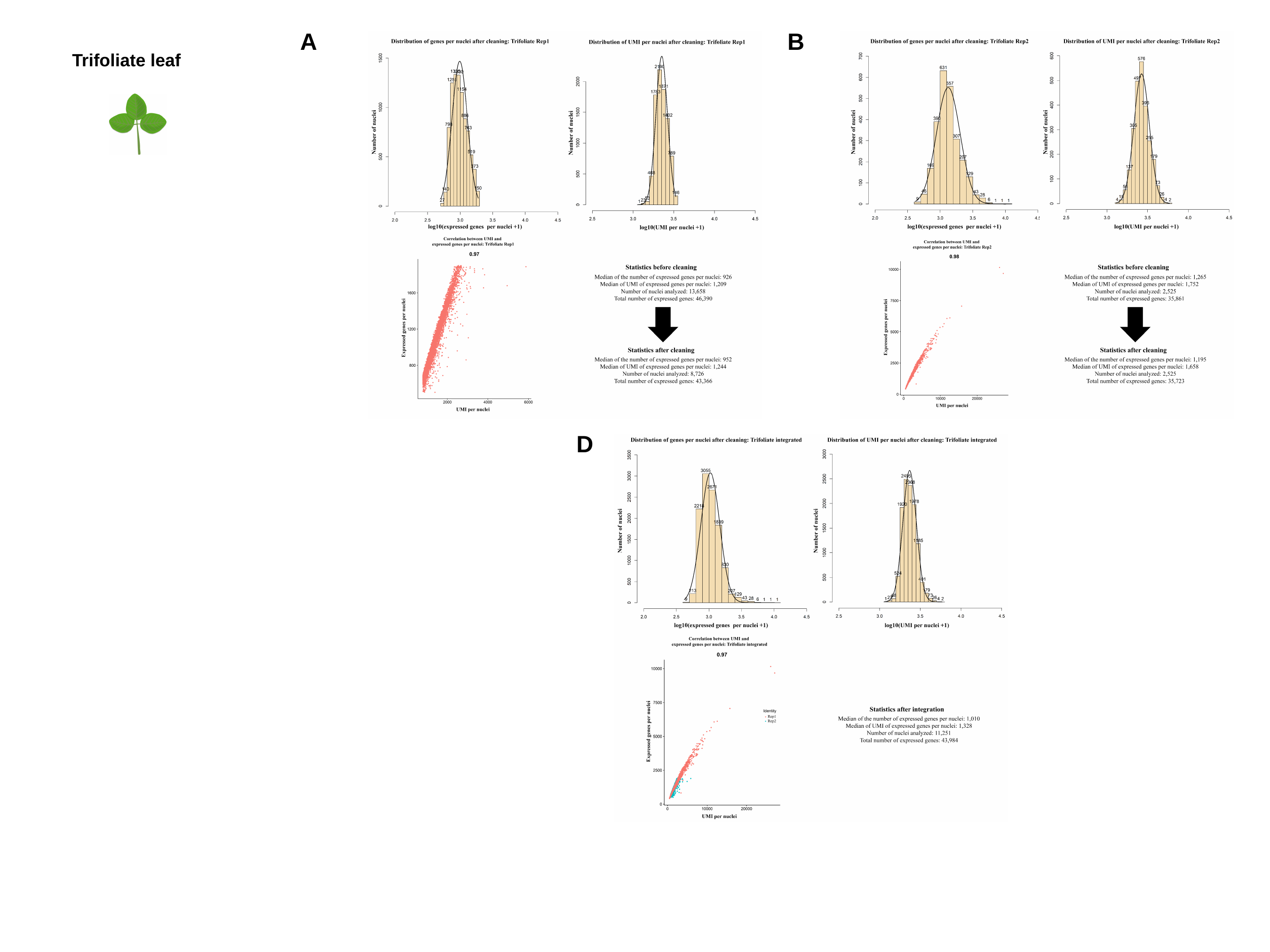

A
B
D
Trifoliate leaf

### Slide 4
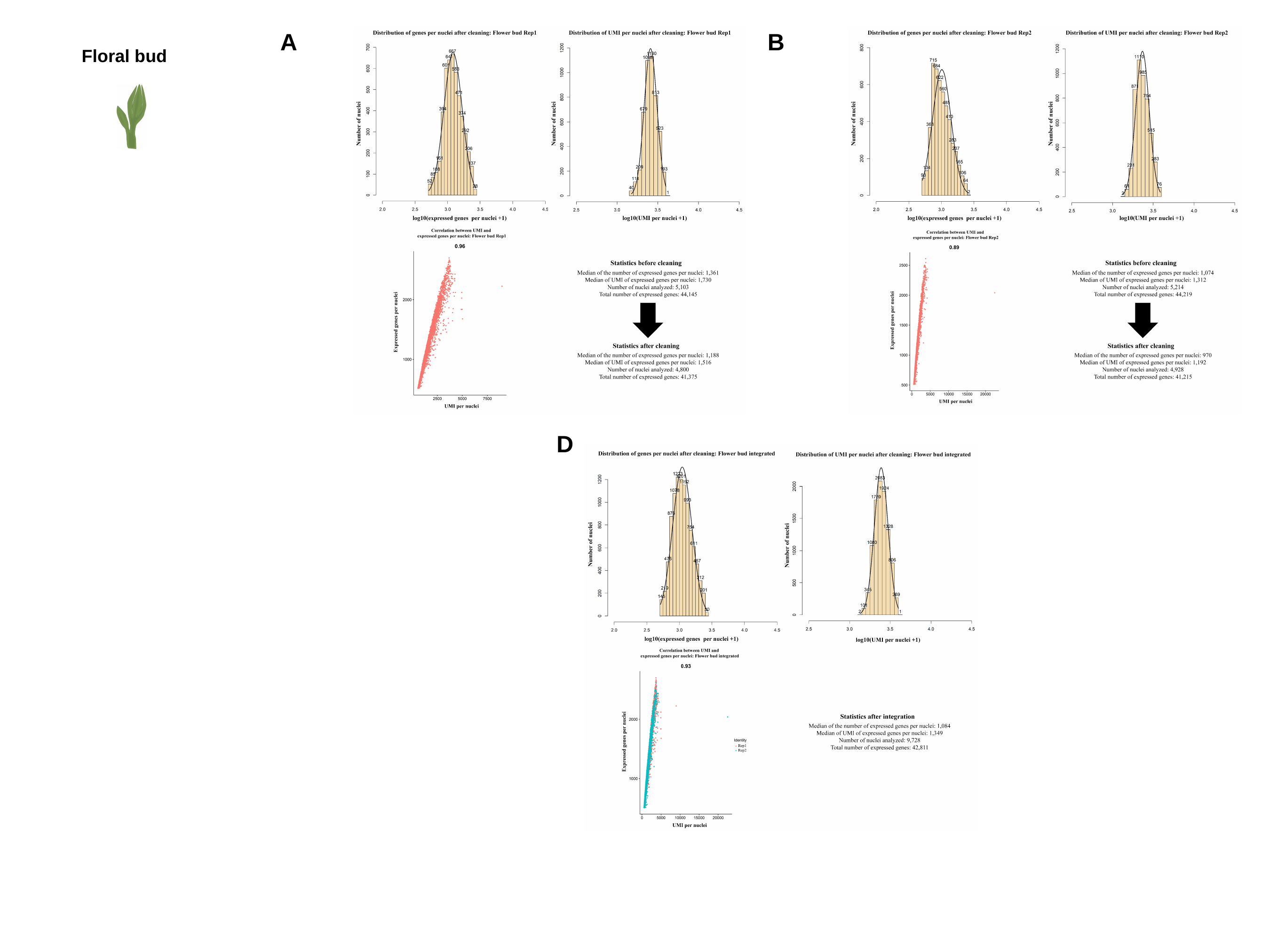

A
B
D
Floral bud

### Slide 5
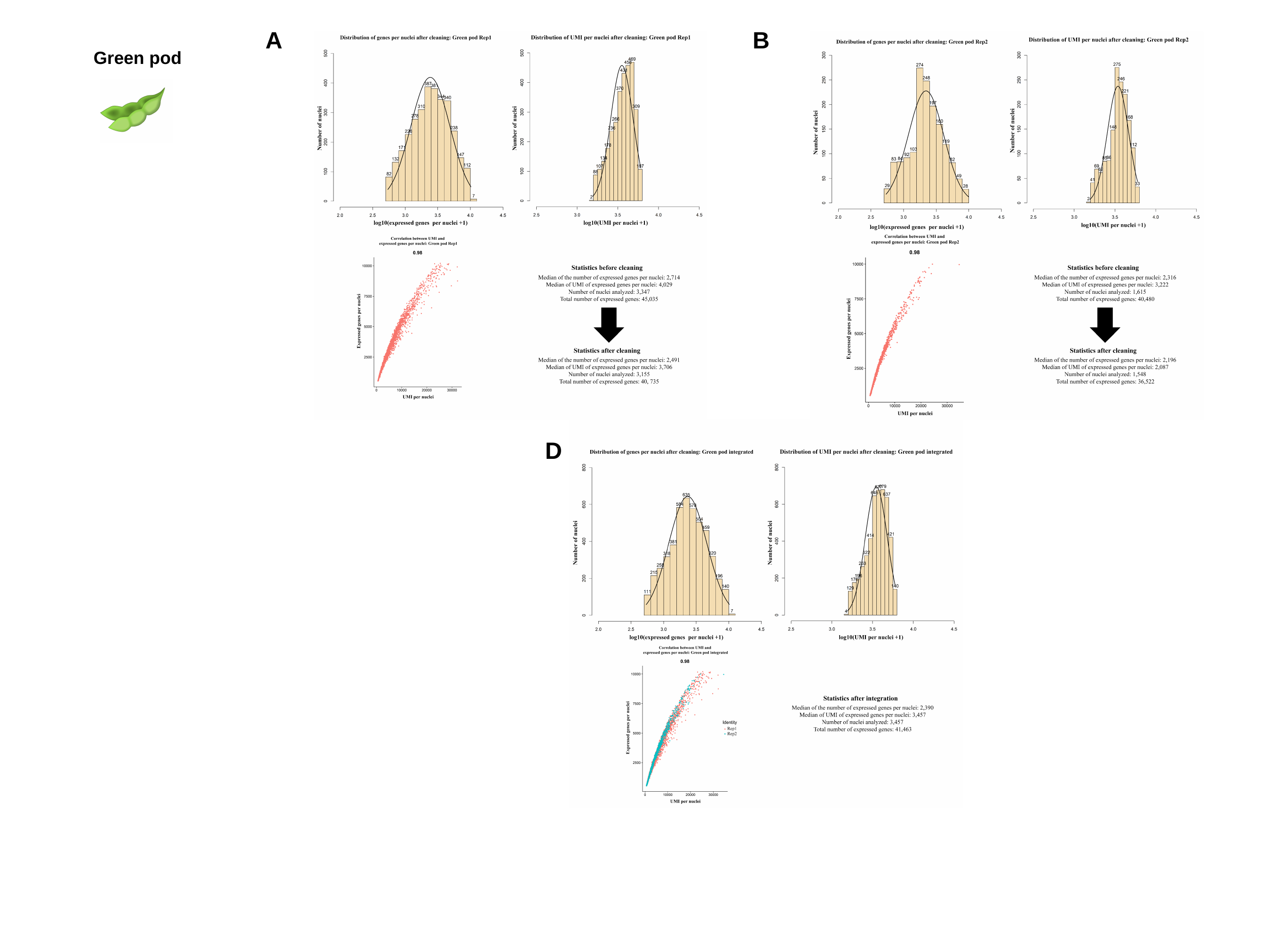

A
B
D
Green pod

### Slide 6
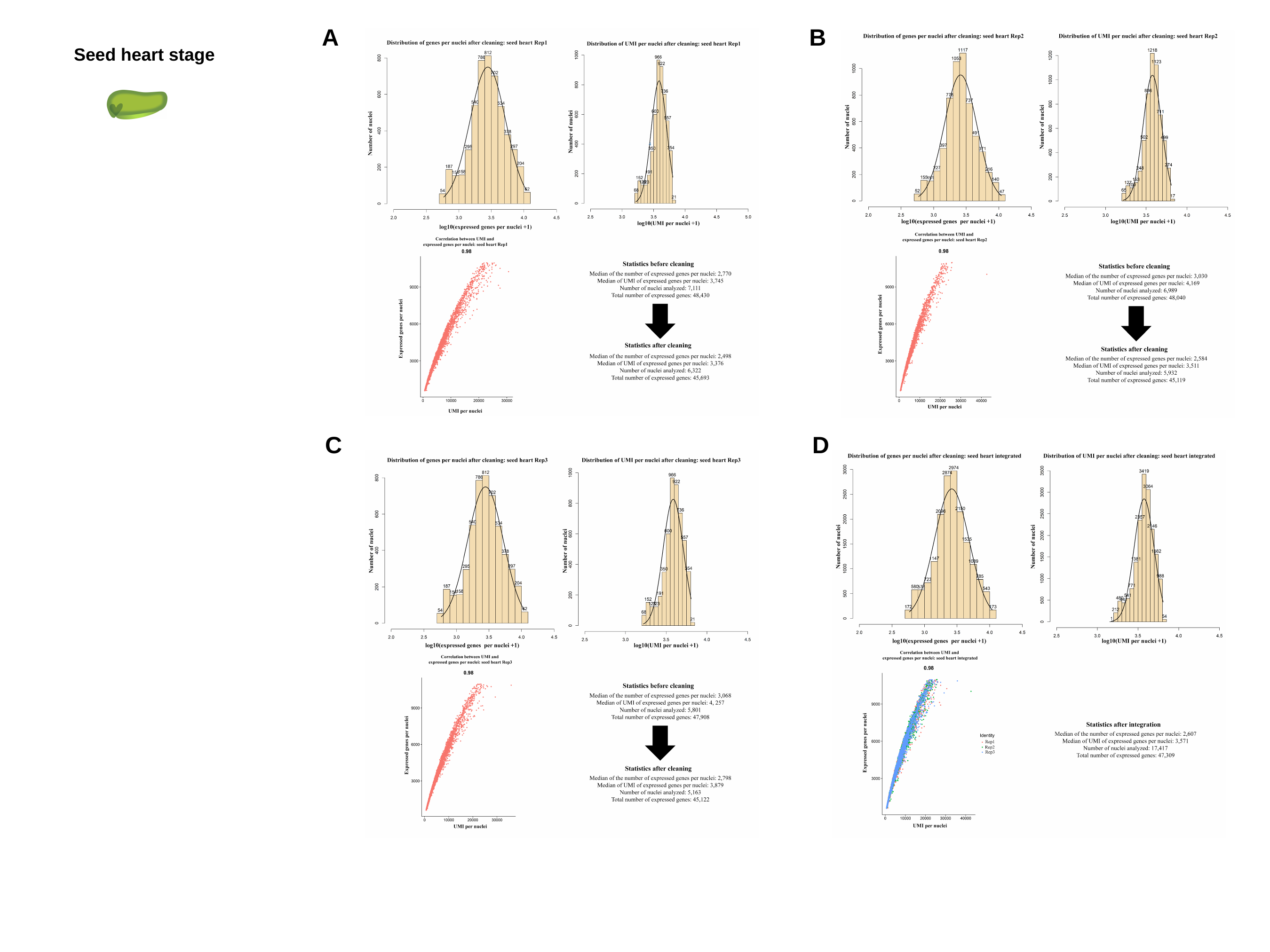

A
B
C
D
Seed heart stage

### Slide 7
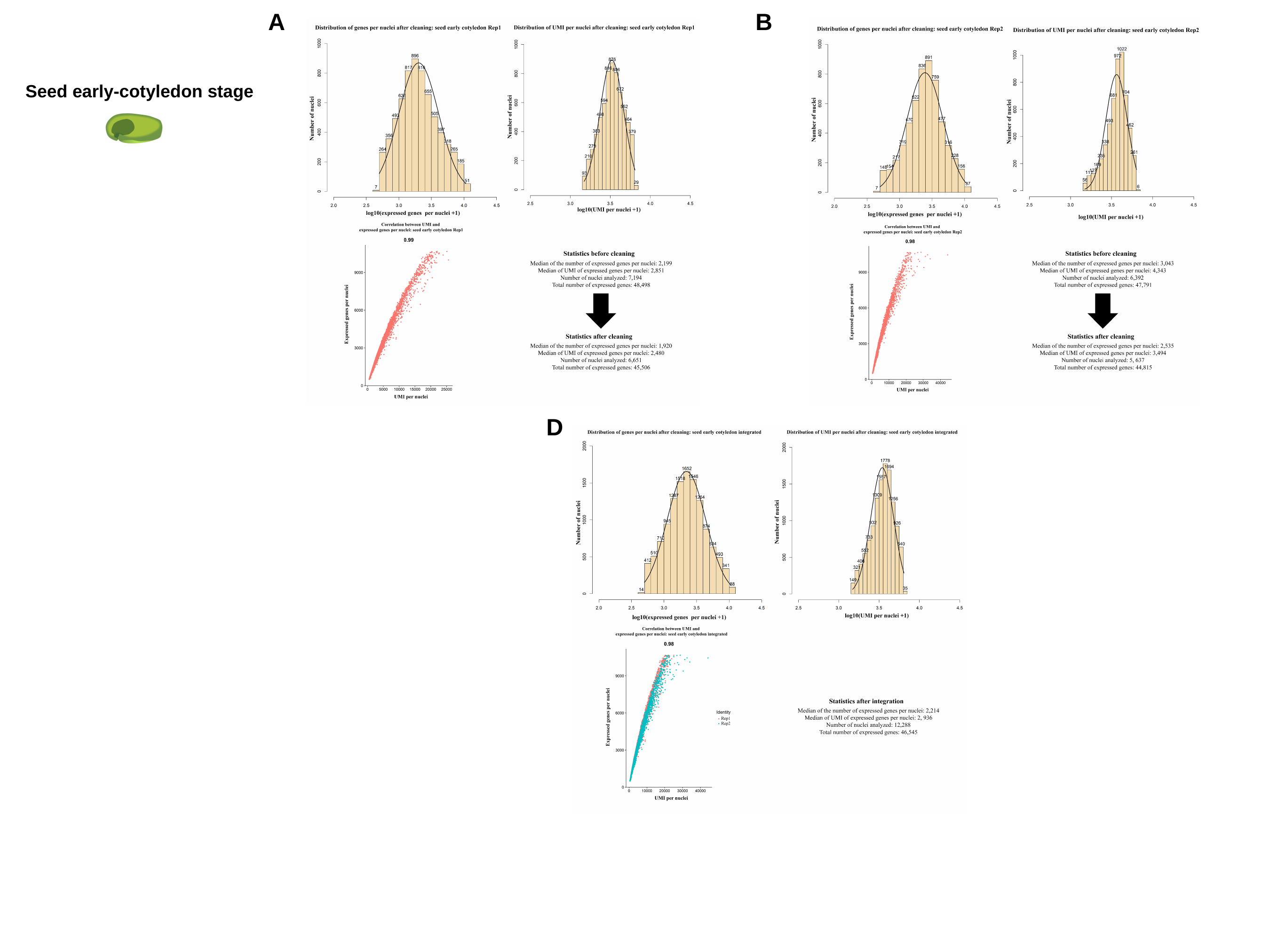

A
B
D
Seed early-cotyledon stage

### Slide 8
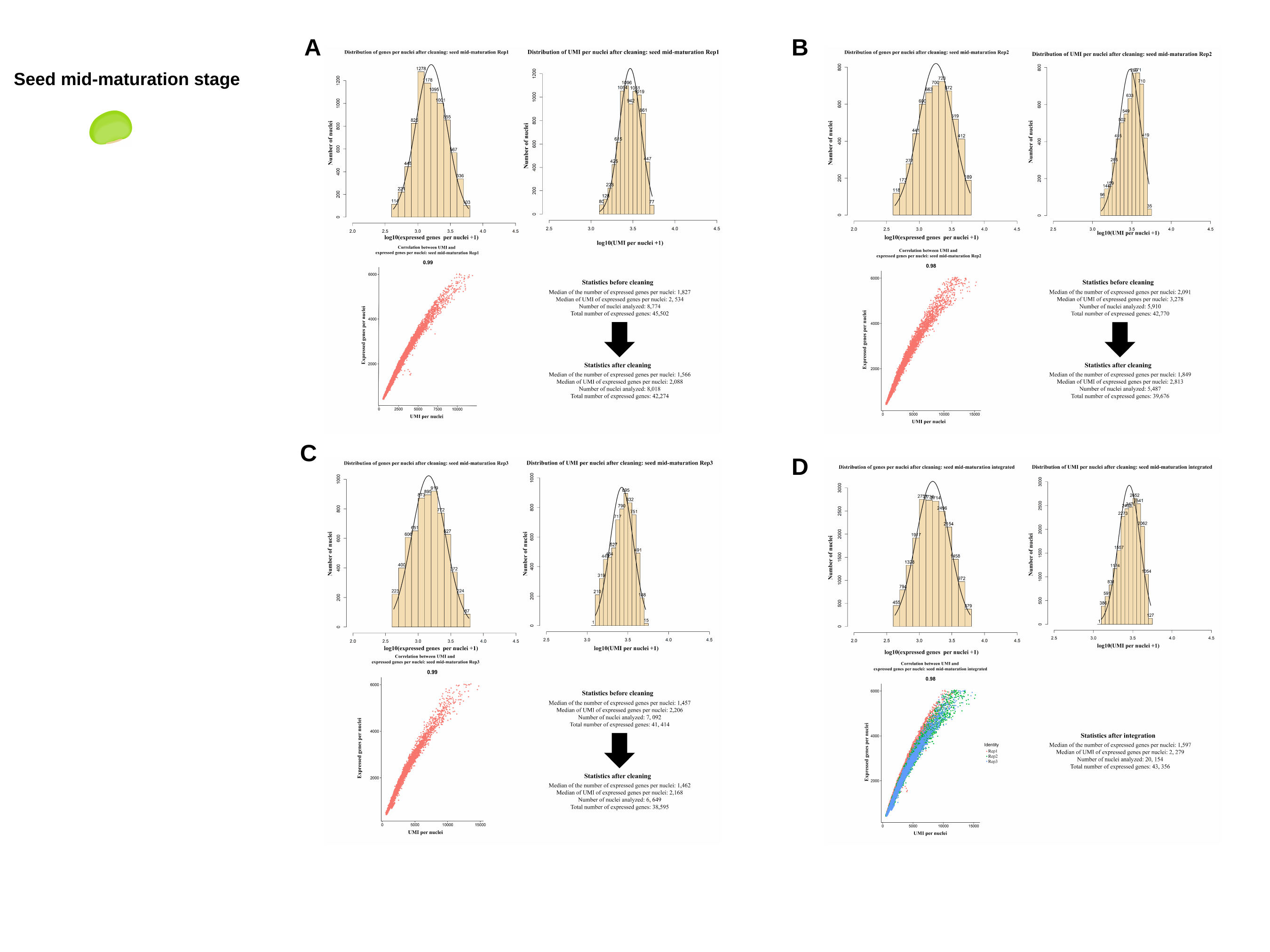

A
B
C
D
Seed mid-maturation stage
