## Supplementary material for "Tabula Glycine: The whole-soybean single-cell resolution transcriptome atlas": FigS4

### Slide 1
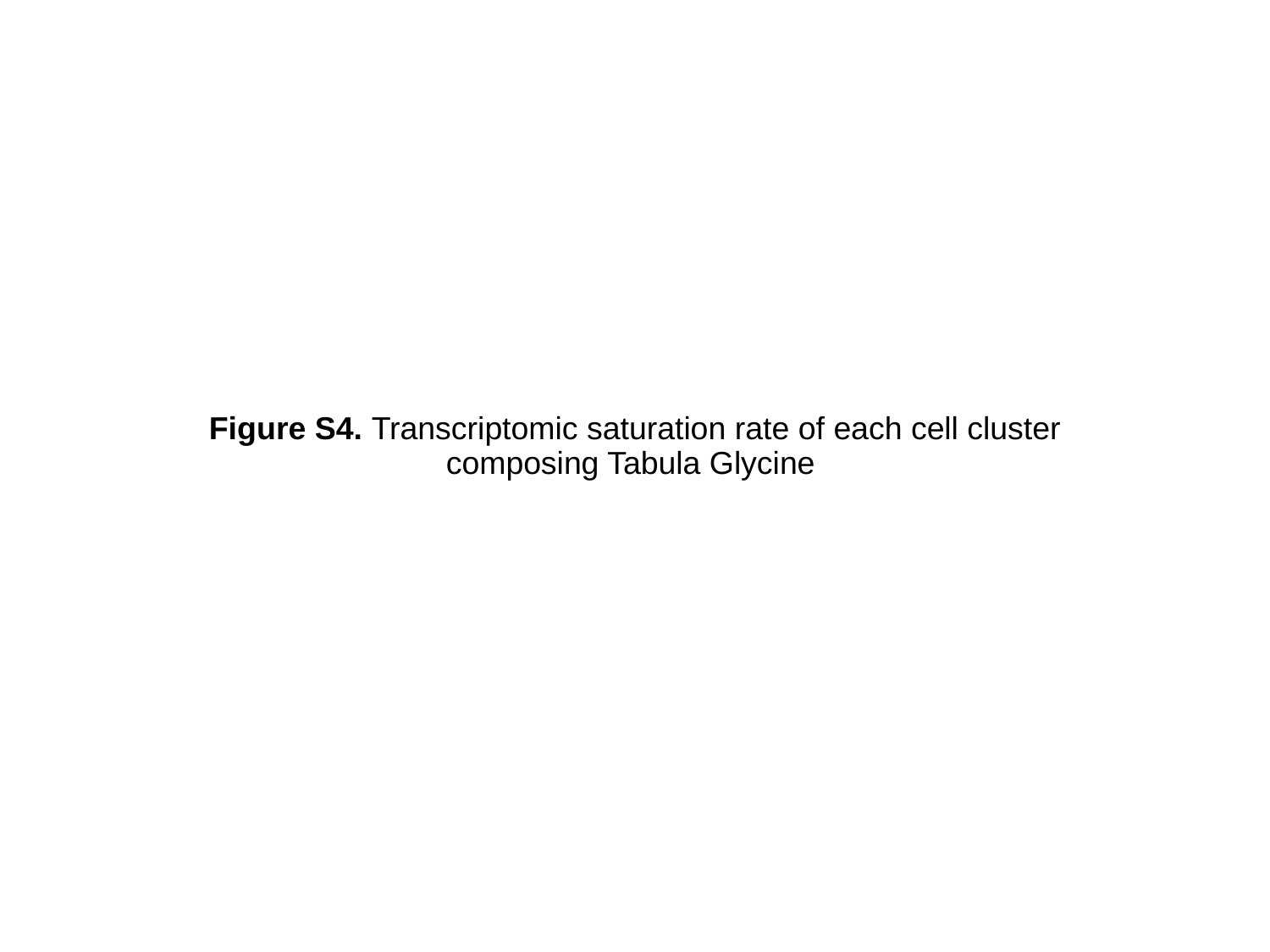

#### Figure S4. Transcriptomic saturation rate of each cell cluster composing Tabula Glycine

### Slide 2
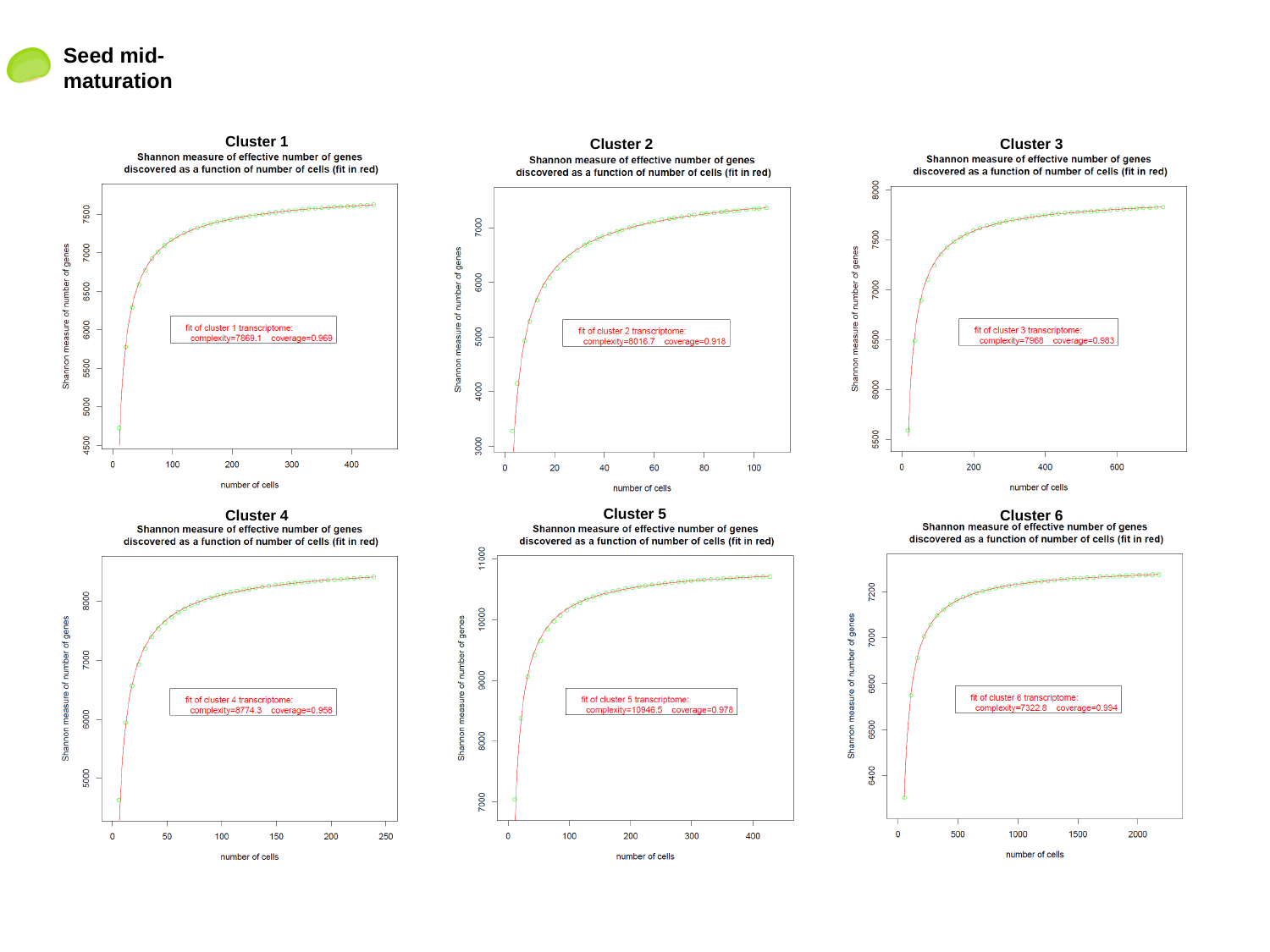

Seed mid-maturation
Cluster 1
Cluster 2
Cluster 3
Cluster 5
Cluster 4
Cluster 6

### Slide 3
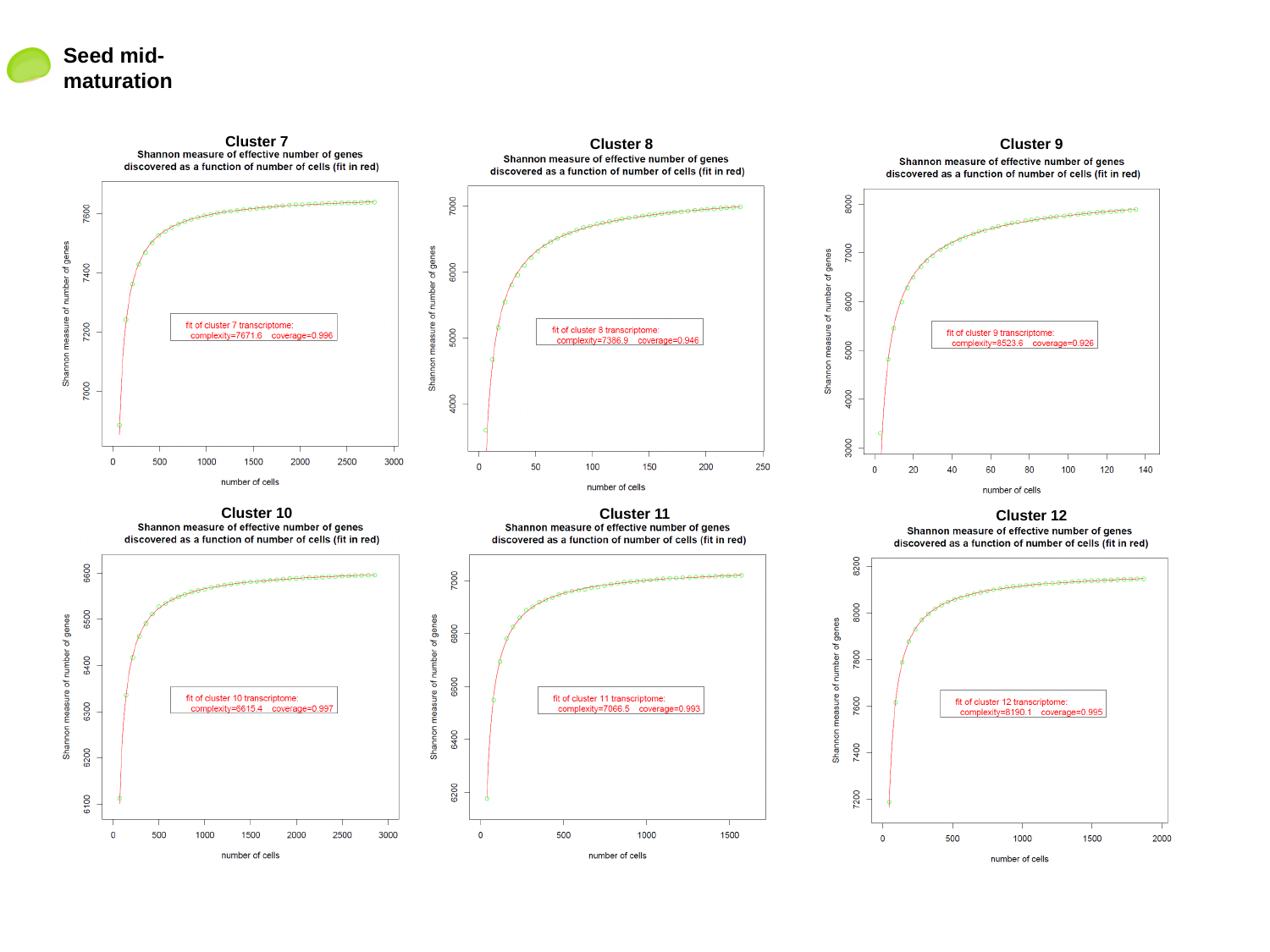

Seed mid-maturation
Cluster 7
Cluster 8
Cluster 9
Cluster 10
Cluster 11
Cluster 12

### Slide 4
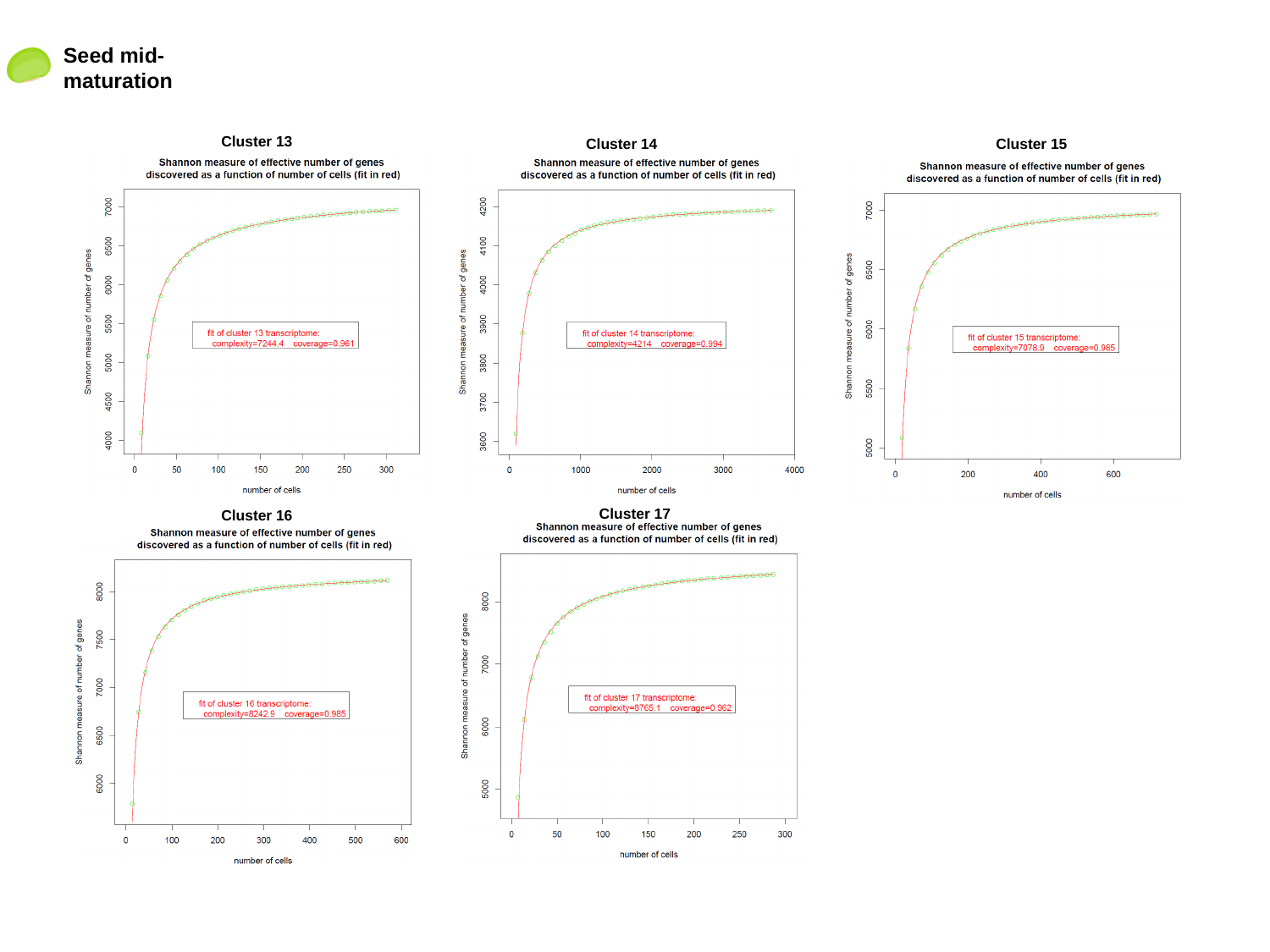

Seed mid-maturation
Cluster 13
Cluster 14
Cluster 15
Cluster 17
Cluster 16

### Slide 5
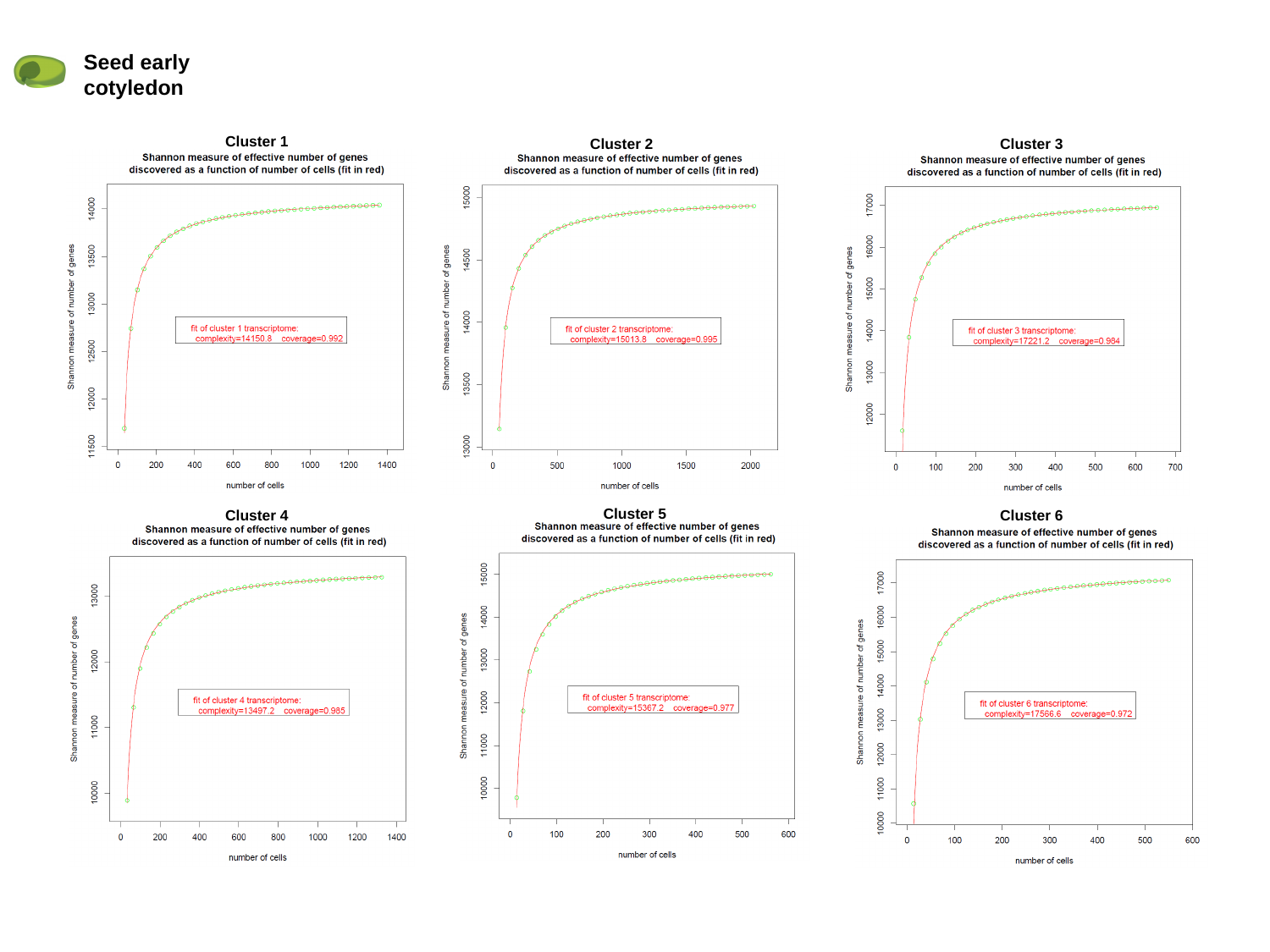

Seed early cotyledon
Cluster 1
Cluster 2
Cluster 3
Cluster 5
Cluster 4
Cluster 6

### Slide 6
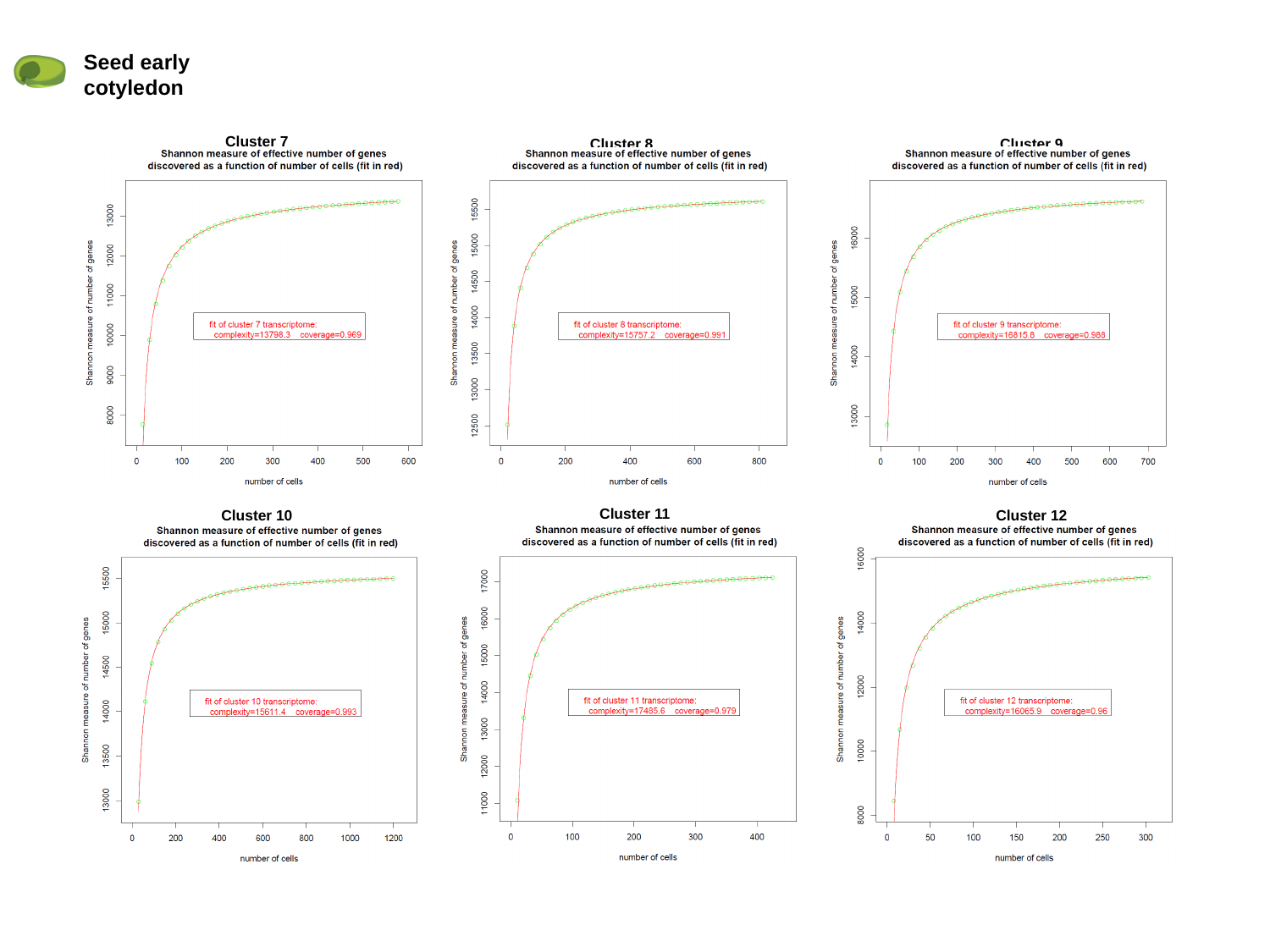

Seed early cotyledon
Cluster 7
Cluster 8
Cluster 9
Cluster 11
Cluster 10
Cluster 12

### Slide 7
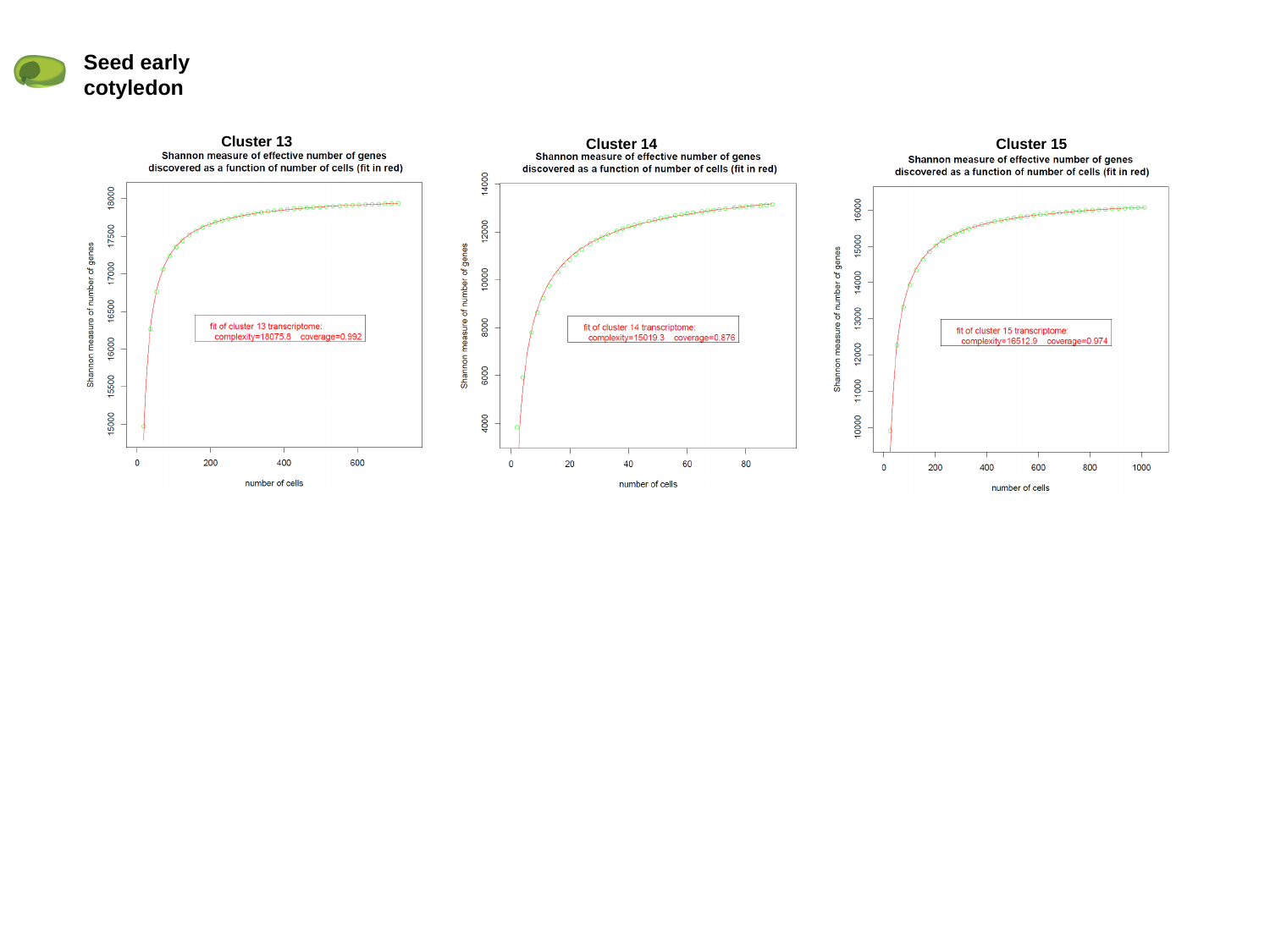

Seed early cotyledon
Cluster 13
Cluster 14
Cluster 15

### Slide 8
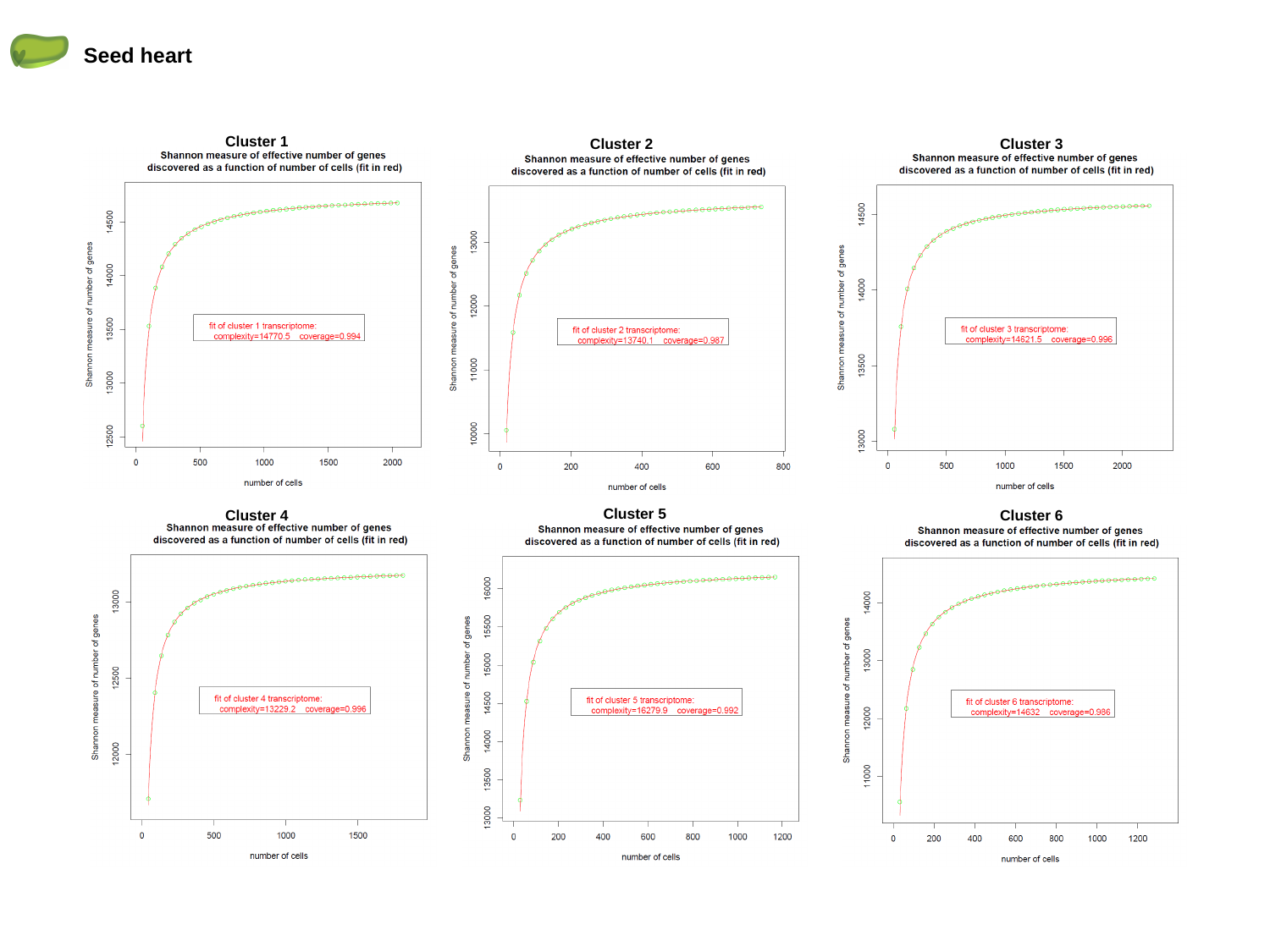

Seed heart
Cluster 1
Cluster 2
Cluster 3
Cluster 5
Cluster 4
Cluster 6

### Slide 9
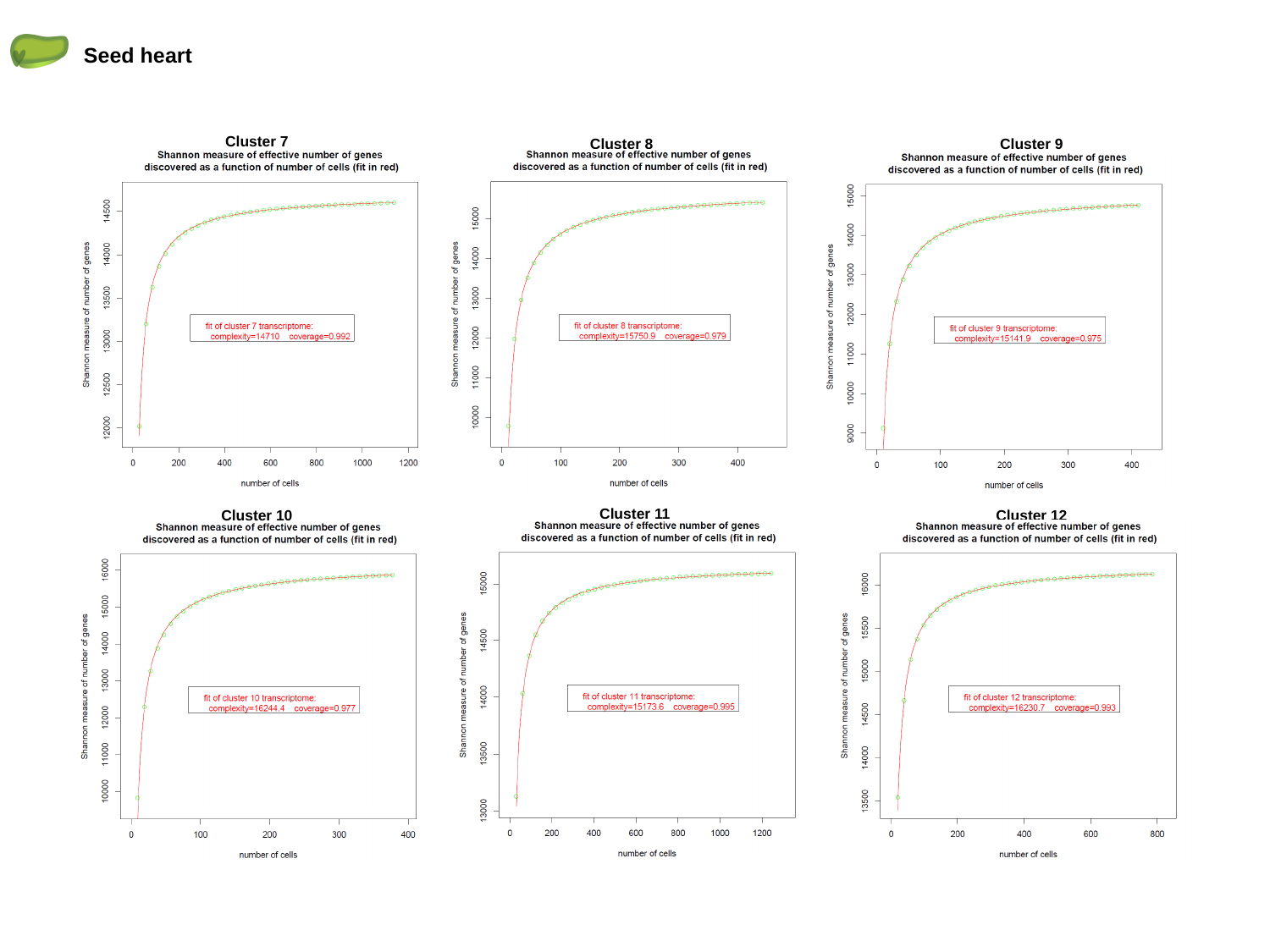

Seed heart
Cluster 7
Cluster 8
Cluster 9
Cluster 11
Cluster 10
Cluster 12

### Slide 10
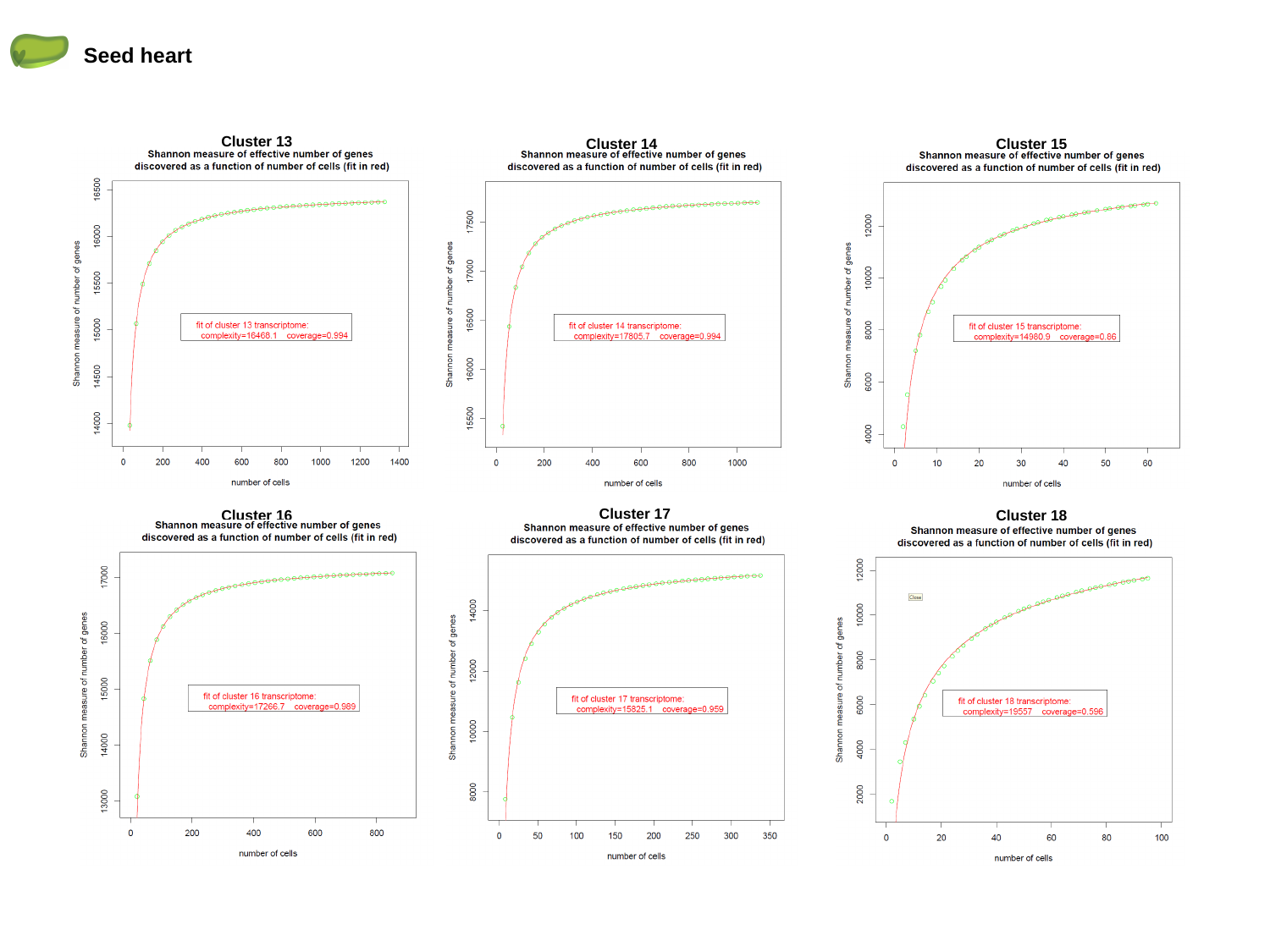

Seed heart
Cluster 13
Cluster 14
Cluster 15
Cluster 17
Cluster 16
Cluster 18

### Slide 11
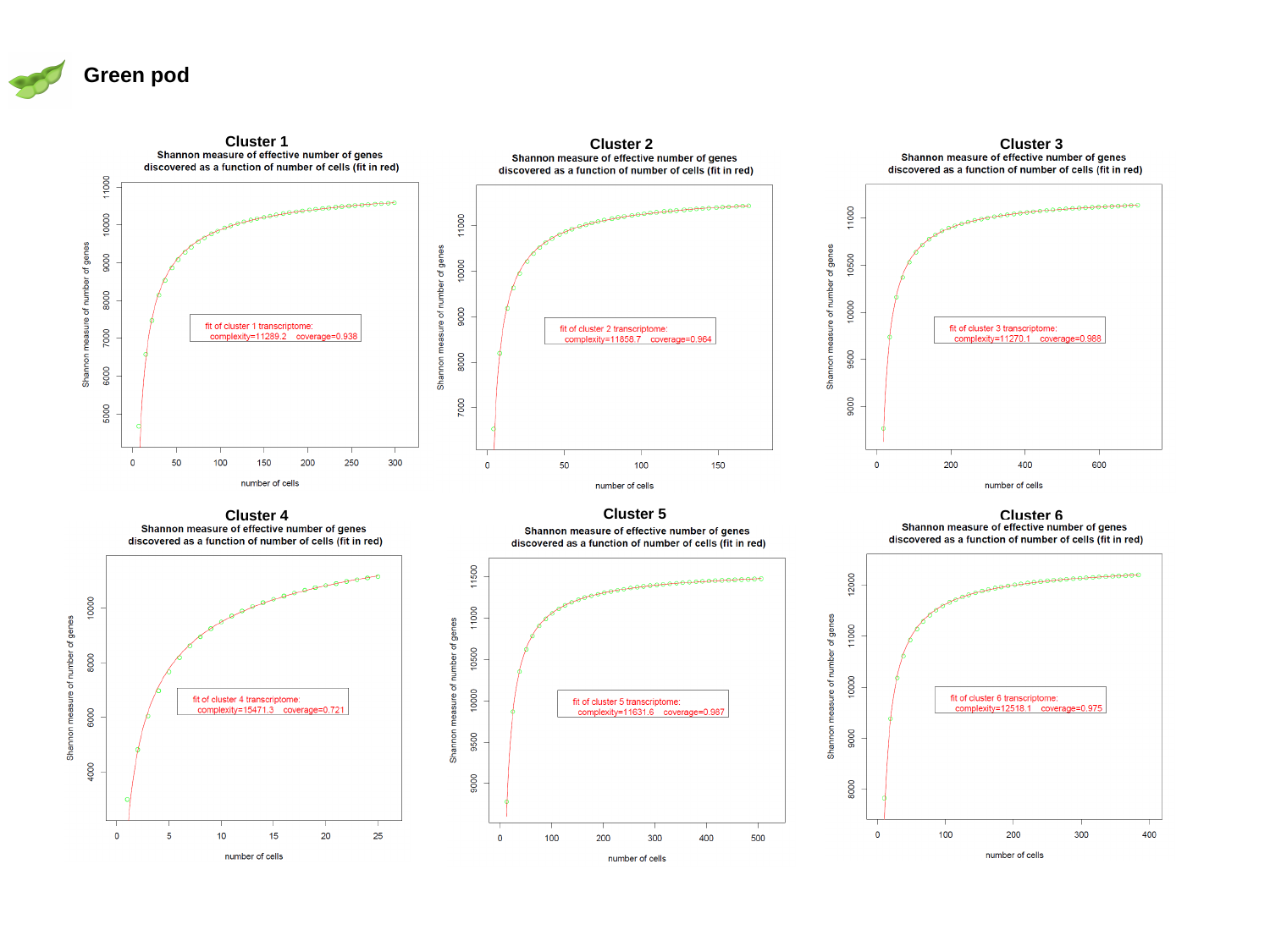

Green pod
Cluster 1
Cluster 2
Cluster 3
Cluster 5
Cluster 4
Cluster 6

### Slide 12
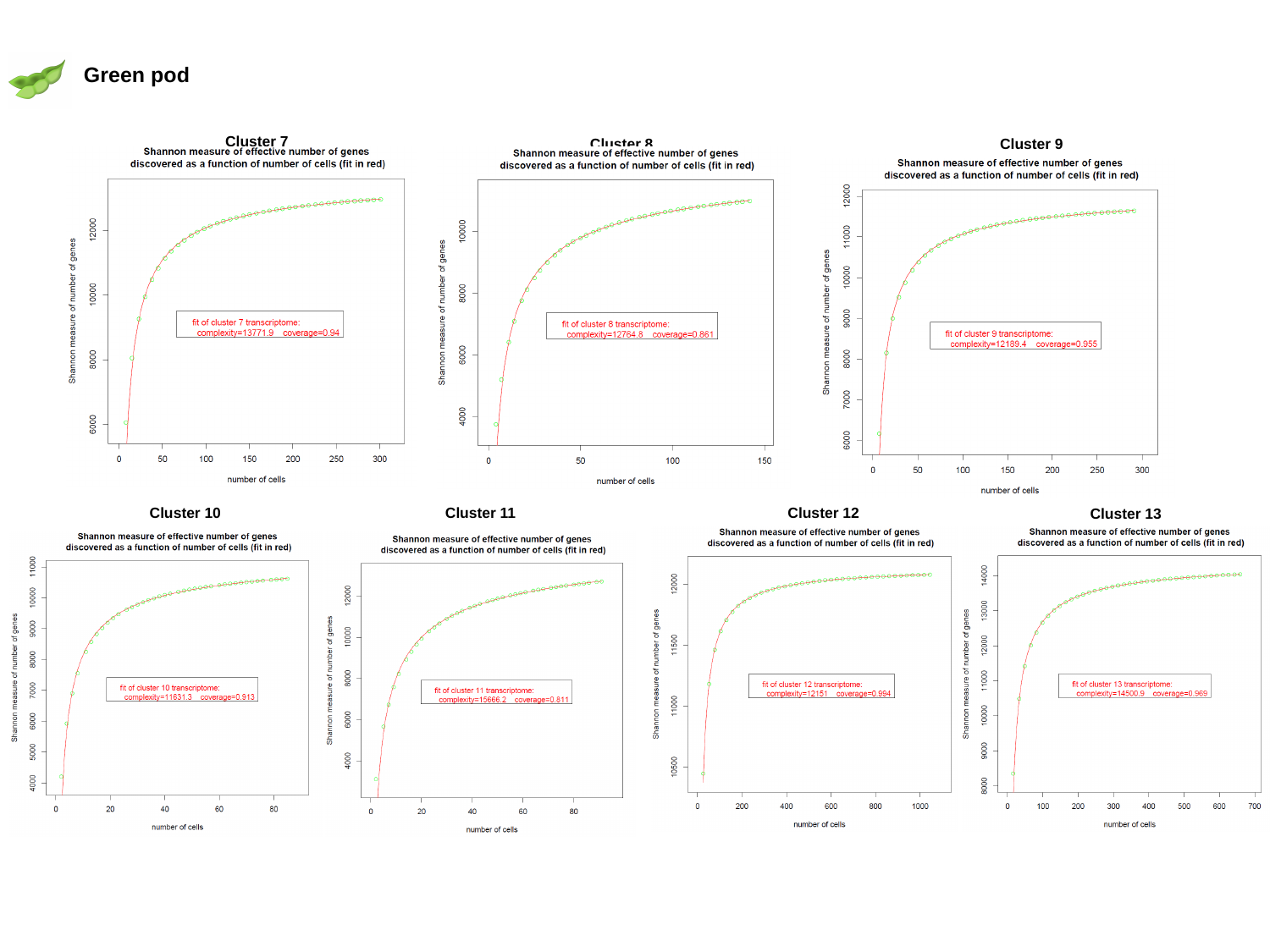

Green pod
Cluster 7
Cluster 8
Cluster 9
Cluster 11
Cluster 12
Cluster 10
Cluster 13

### Slide 13
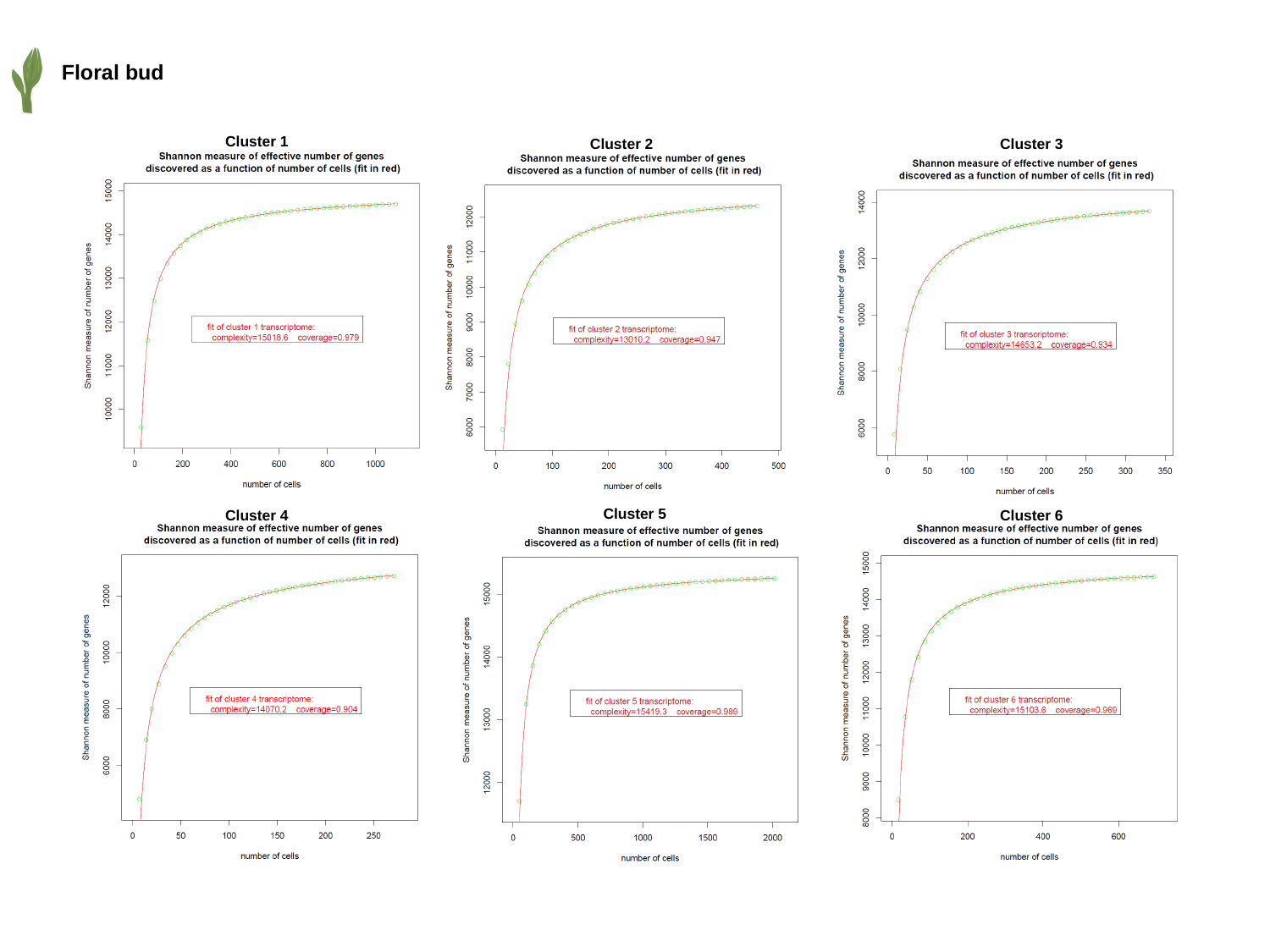

Floral bud
Cluster 1
Cluster 2
Cluster 3
Cluster 5
Cluster 4
Cluster 6

### Slide 14
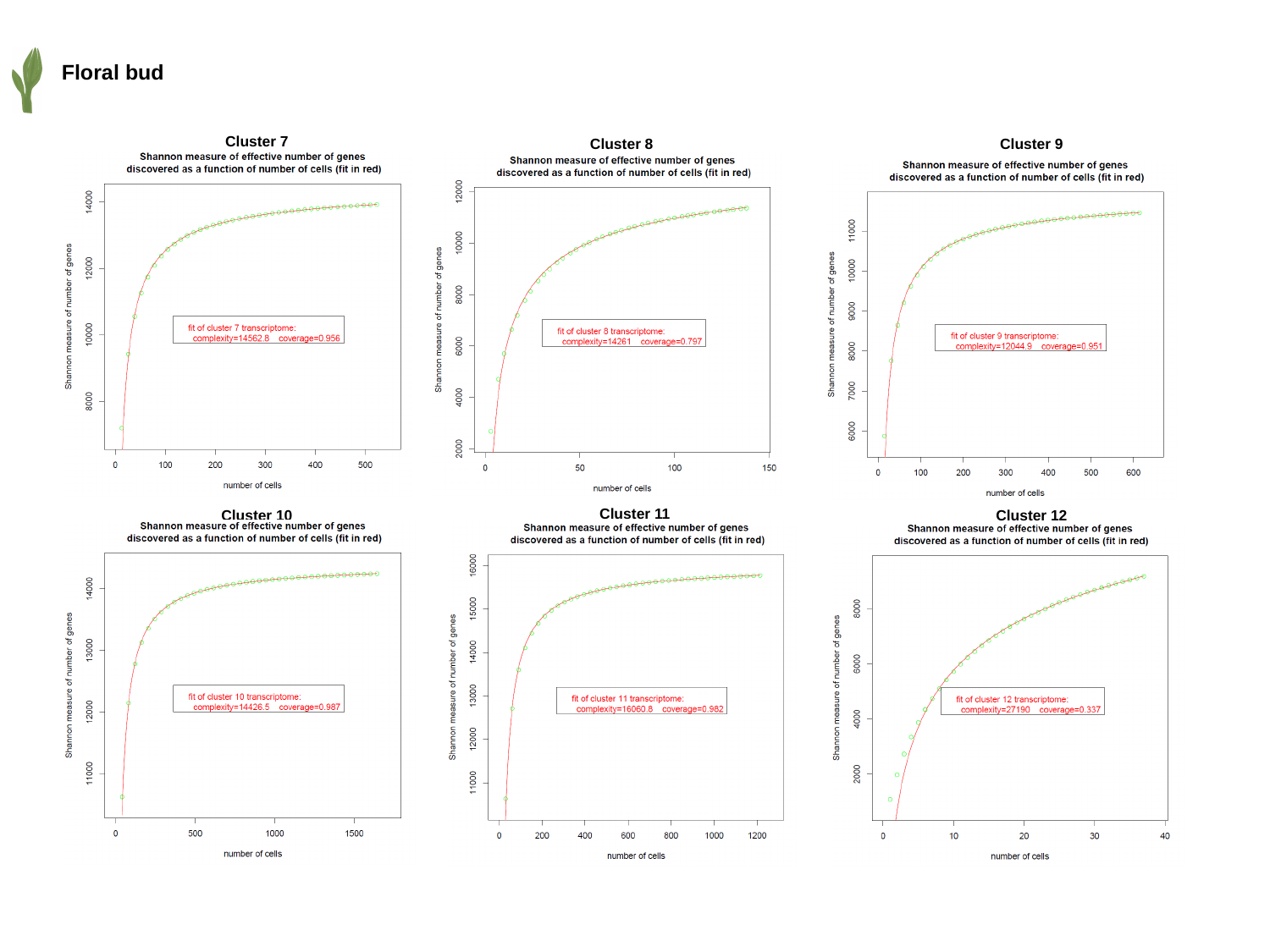

Floral bud
Cluster 7
Cluster 8
Cluster 9
Cluster 11
Cluster 10
Cluster 12

### Slide 15
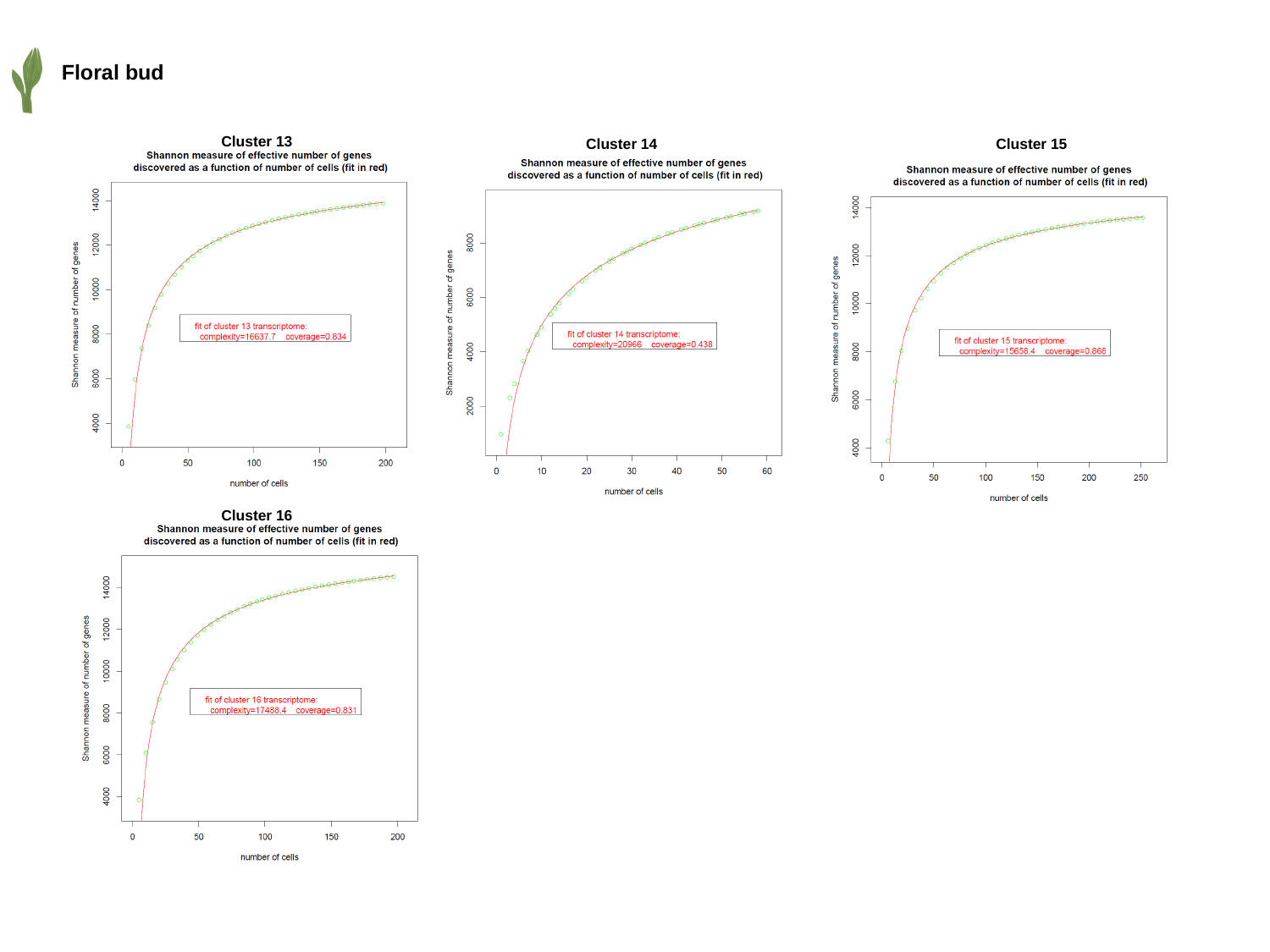

Floral bud
Cluster 13
Cluster 14
Cluster 15
Cluster 16

### Slide 16
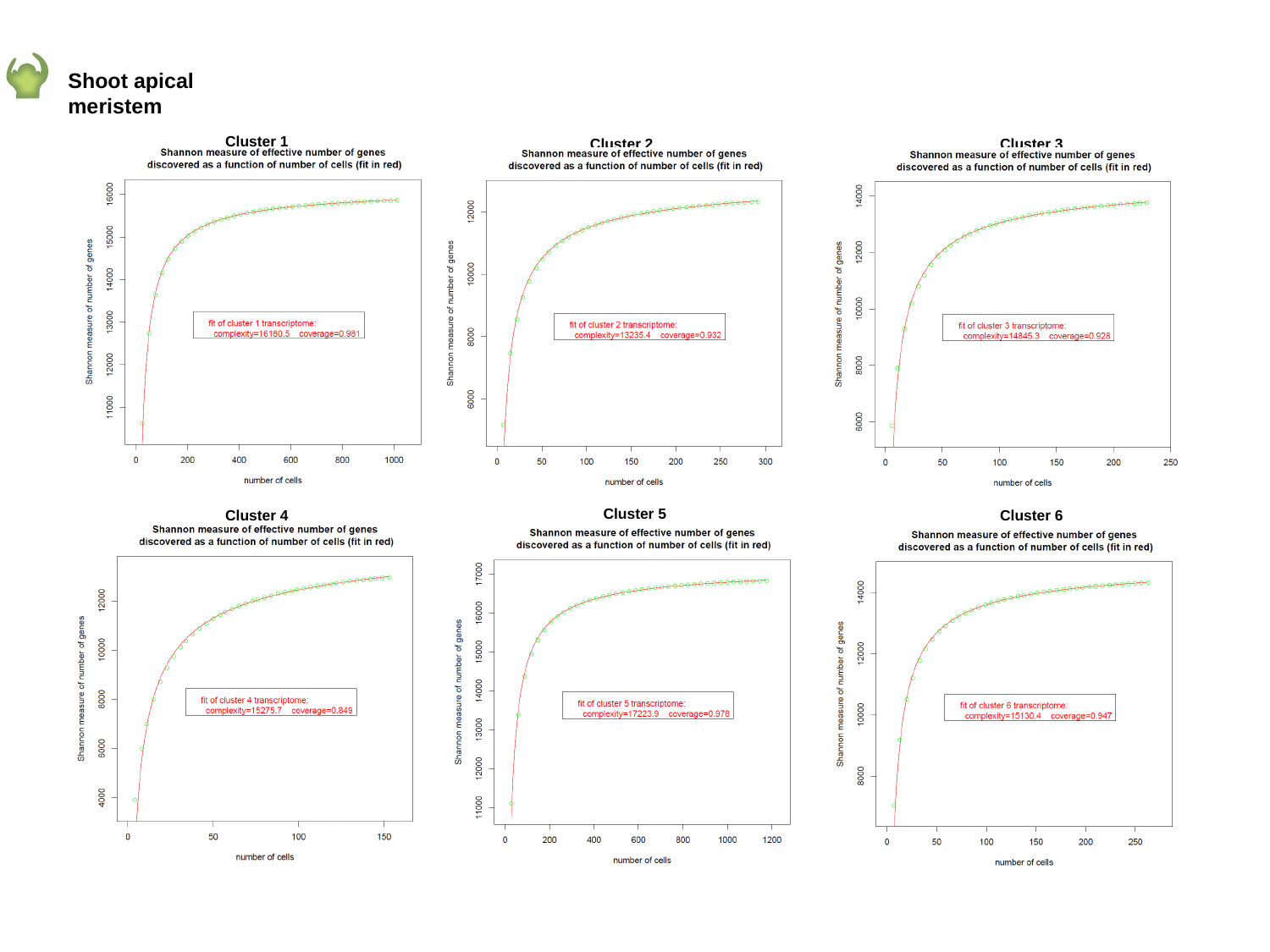

Shoot apical meristem
Cluster 1
Cluster 2
Cluster 3
Cluster 5
Cluster 4
Cluster 6

### Slide 17
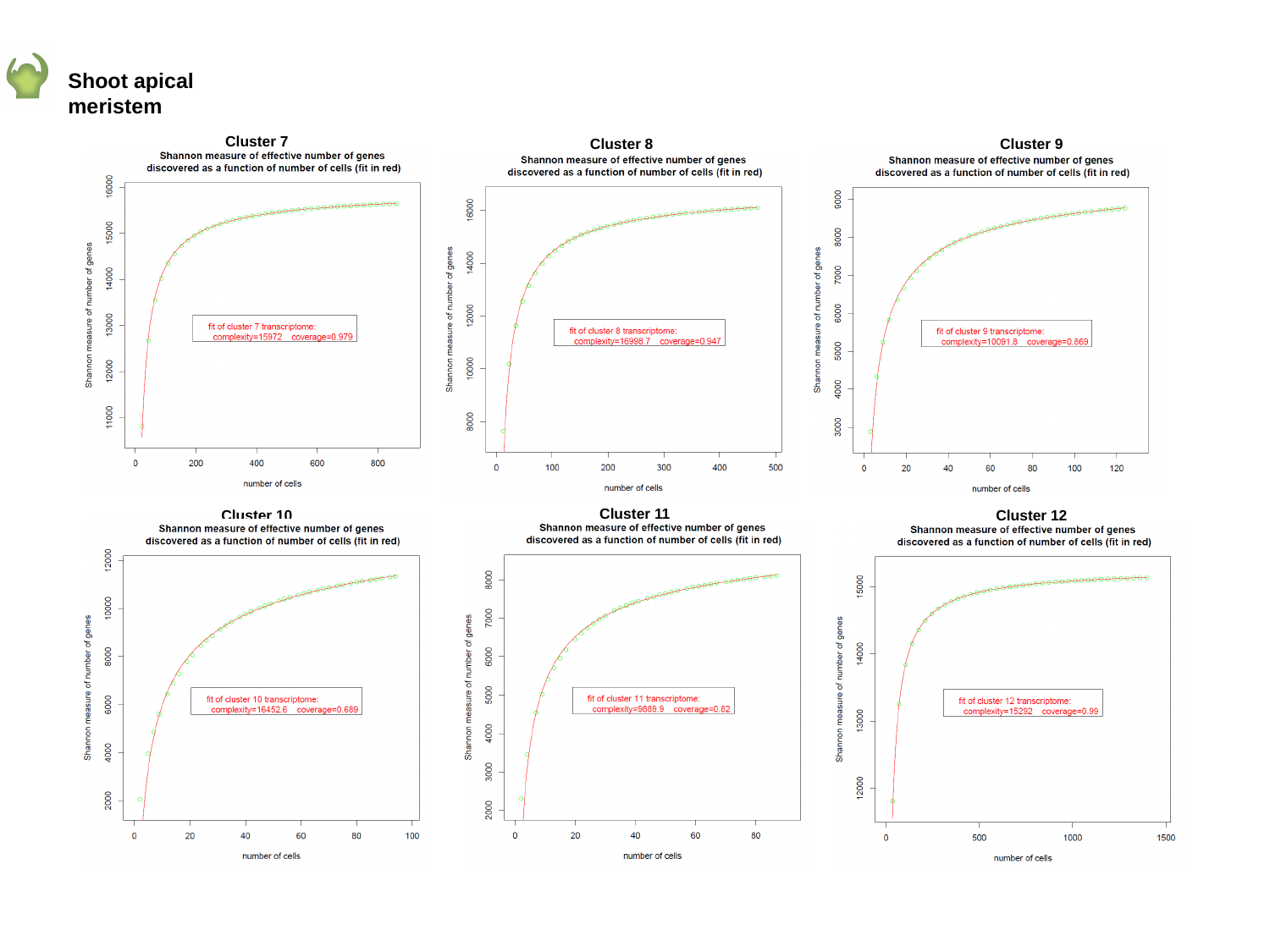

Shoot apical meristem
Cluster 7
Cluster 8
Cluster 9
Cluster 11
Cluster 10
Cluster 12

### Slide 18
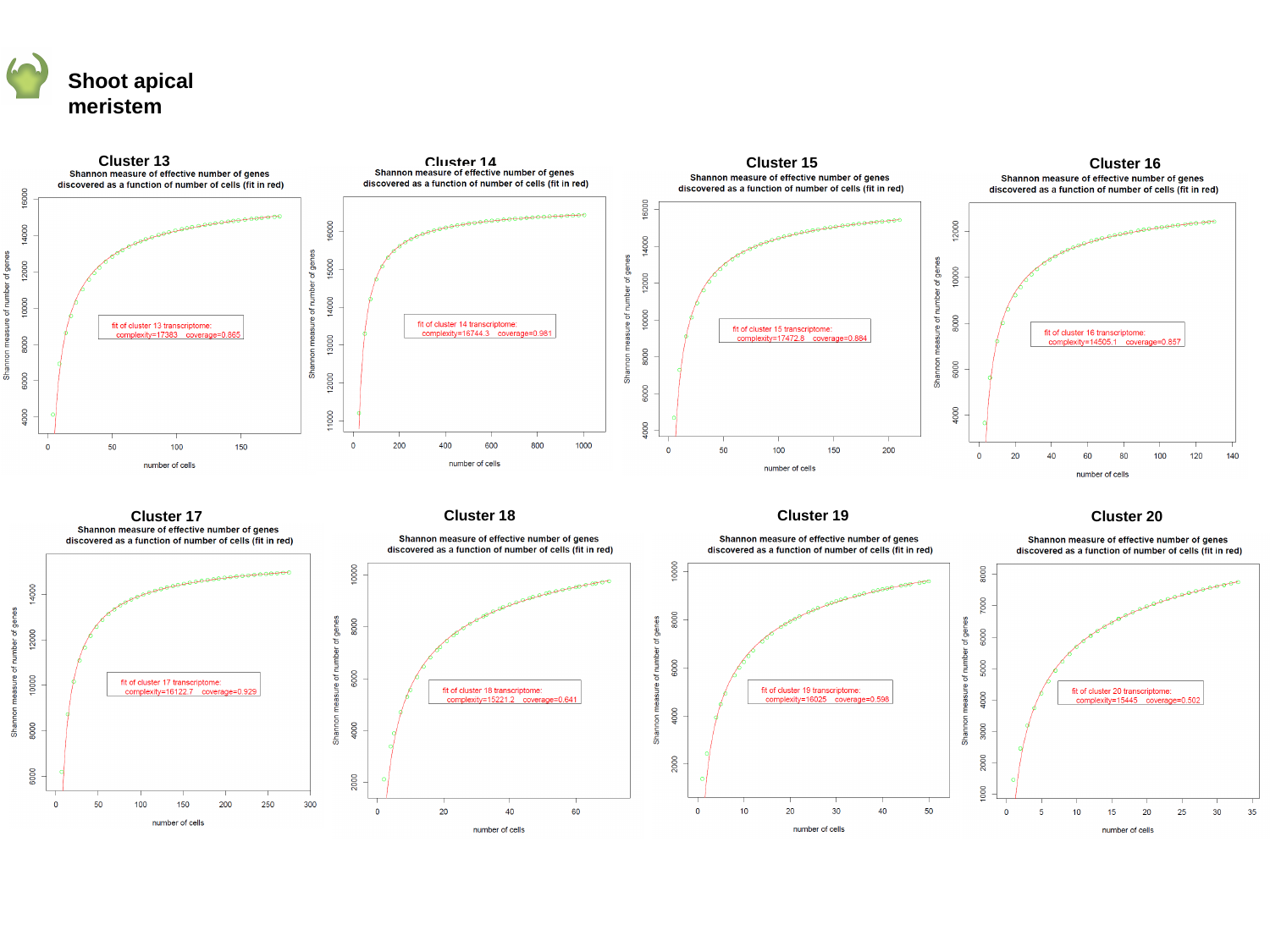

Shoot apical meristem
Cluster 13
Cluster 14
Cluster 15
Cluster 16
Cluster 18
Cluster 19
Cluster 17
Cluster 20

### Slide 19
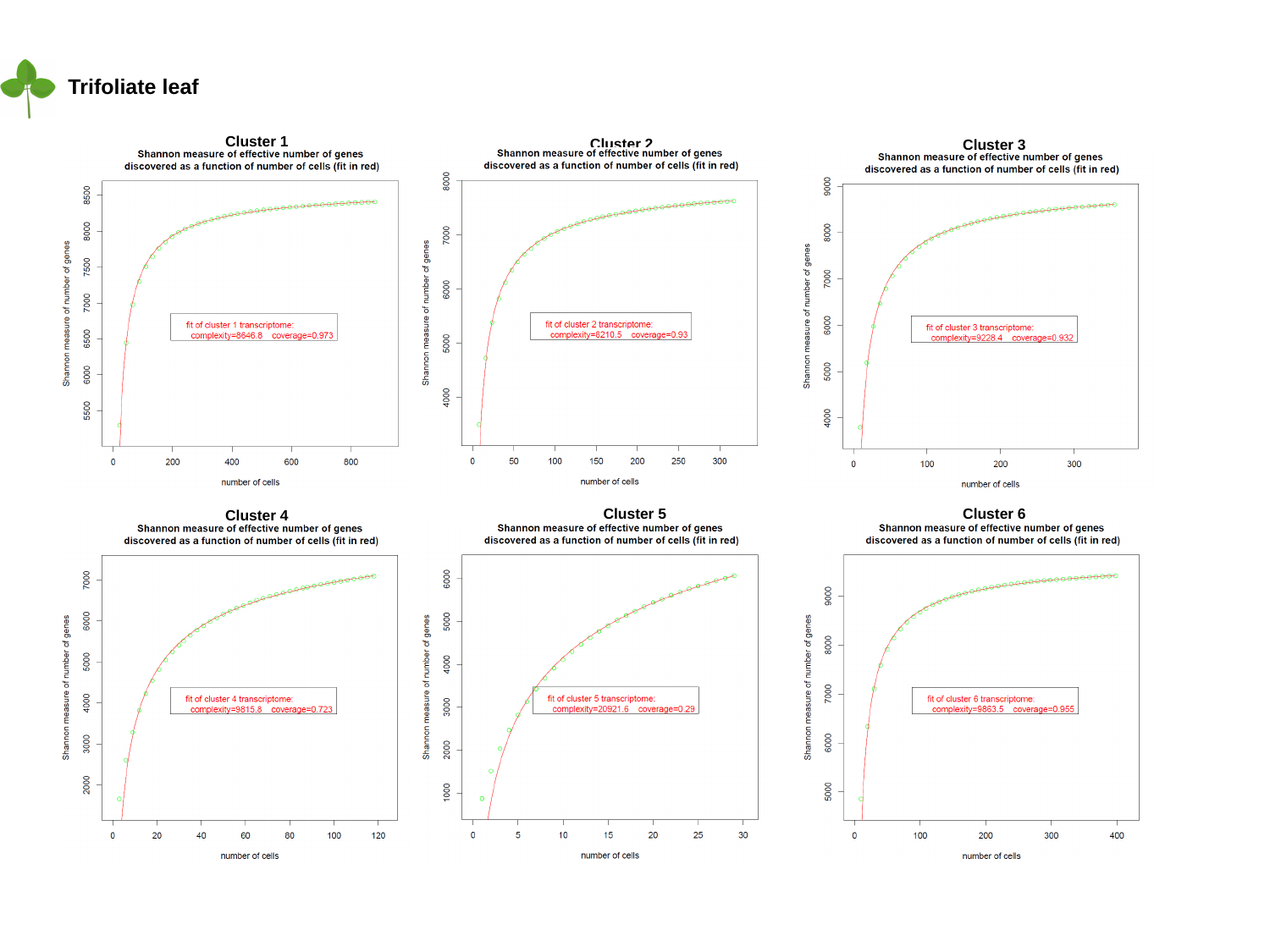

Trifoliate leaf
Cluster 1
Cluster 2
Cluster 3
Cluster 5
Cluster 6
Cluster 4

### Slide 20
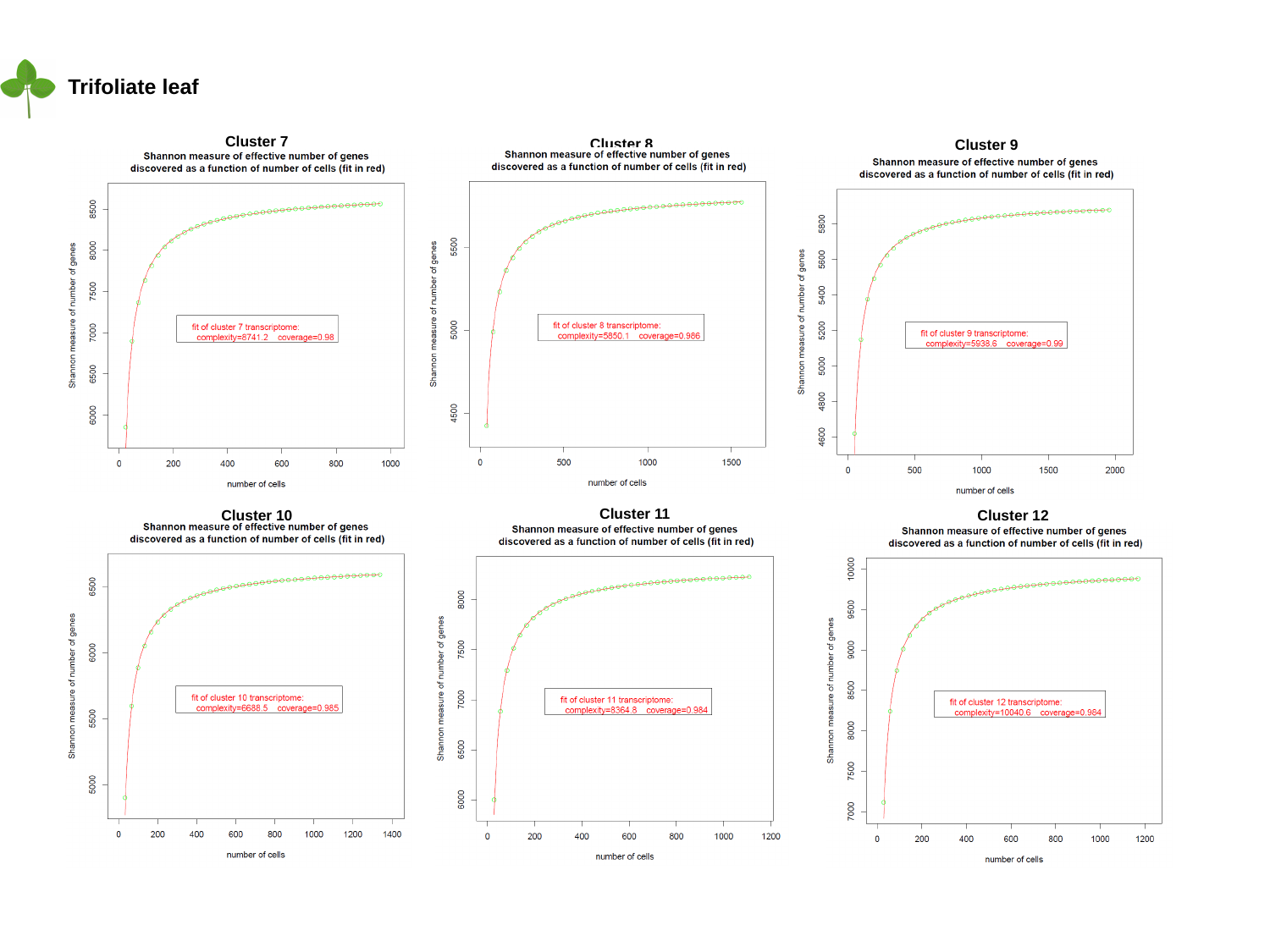

Trifoliate leaf
Cluster 7
Cluster 8
Cluster 9
Cluster 11
Cluster 10
Cluster 12

### Slide 21
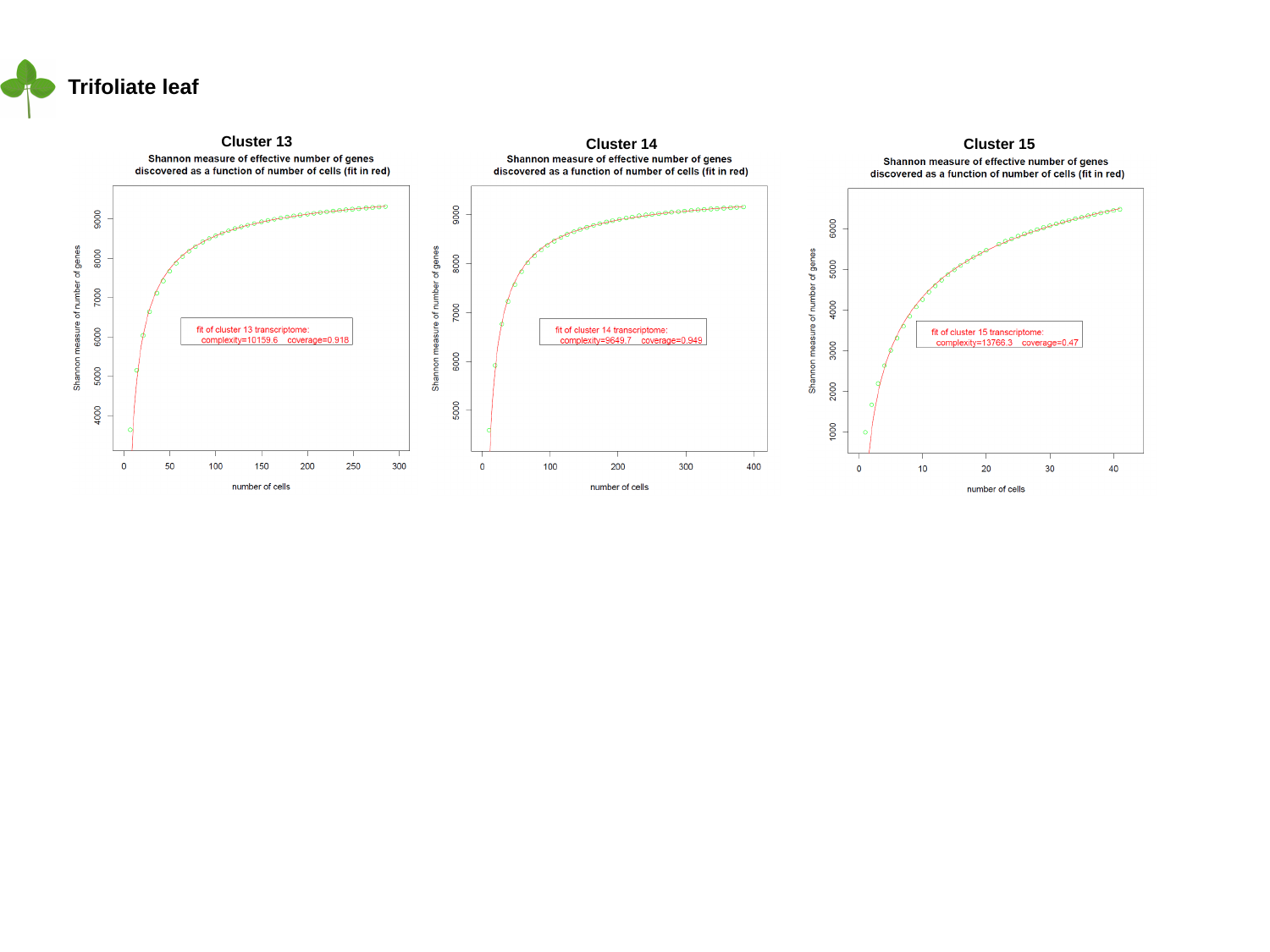

Trifoliate leaf
Cluster 13
Cluster 14
Cluster 15

### Slide 22
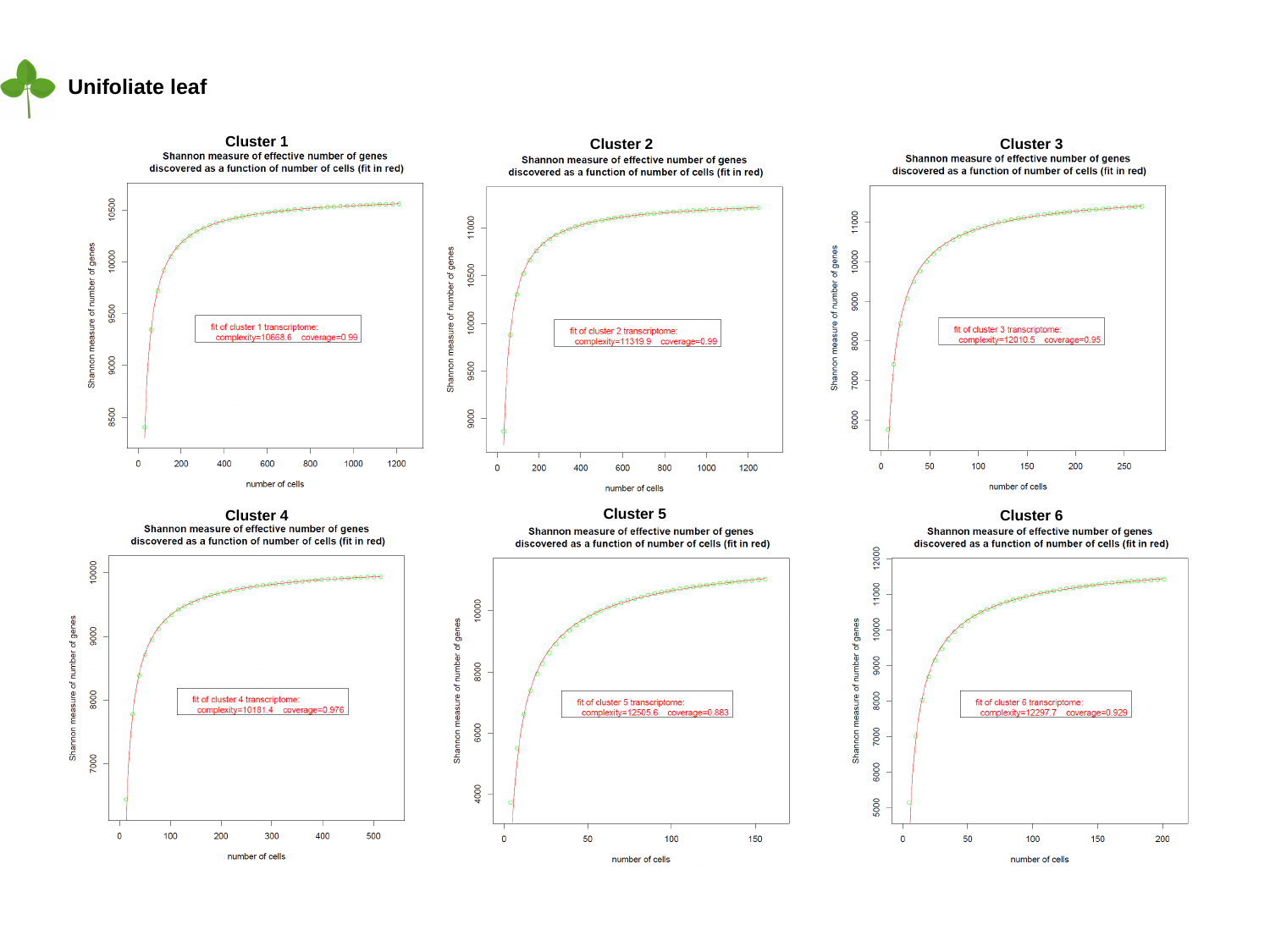

Unifoliate leaf
Cluster 1
Cluster 2
Cluster 3
Cluster 5
Cluster 4
Cluster 6

### Slide 23
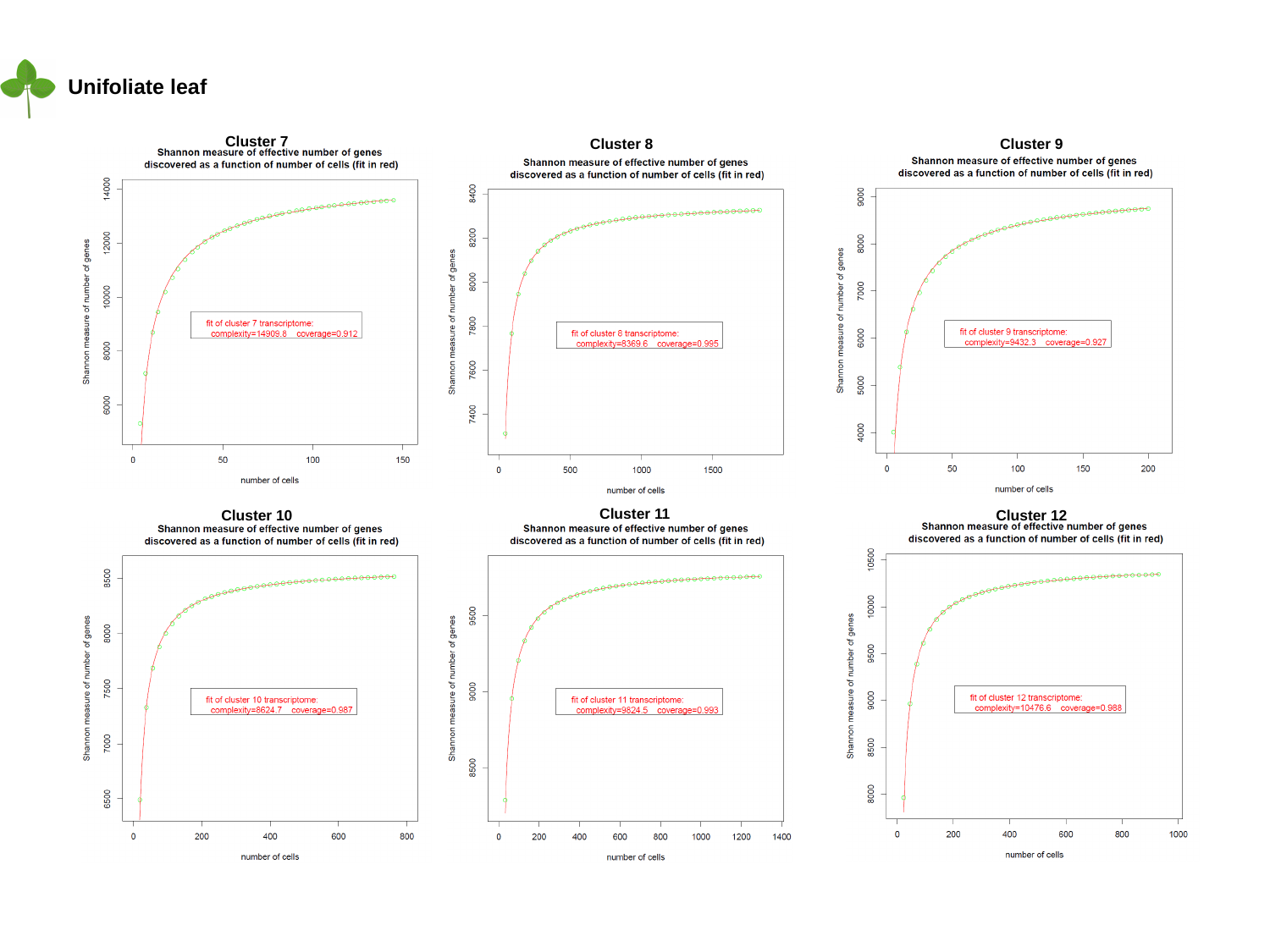

Unifoliate leaf
Cluster 7
Cluster 8
Cluster 9
Cluster 11
Cluster 10
Cluster 12

### Slide 24
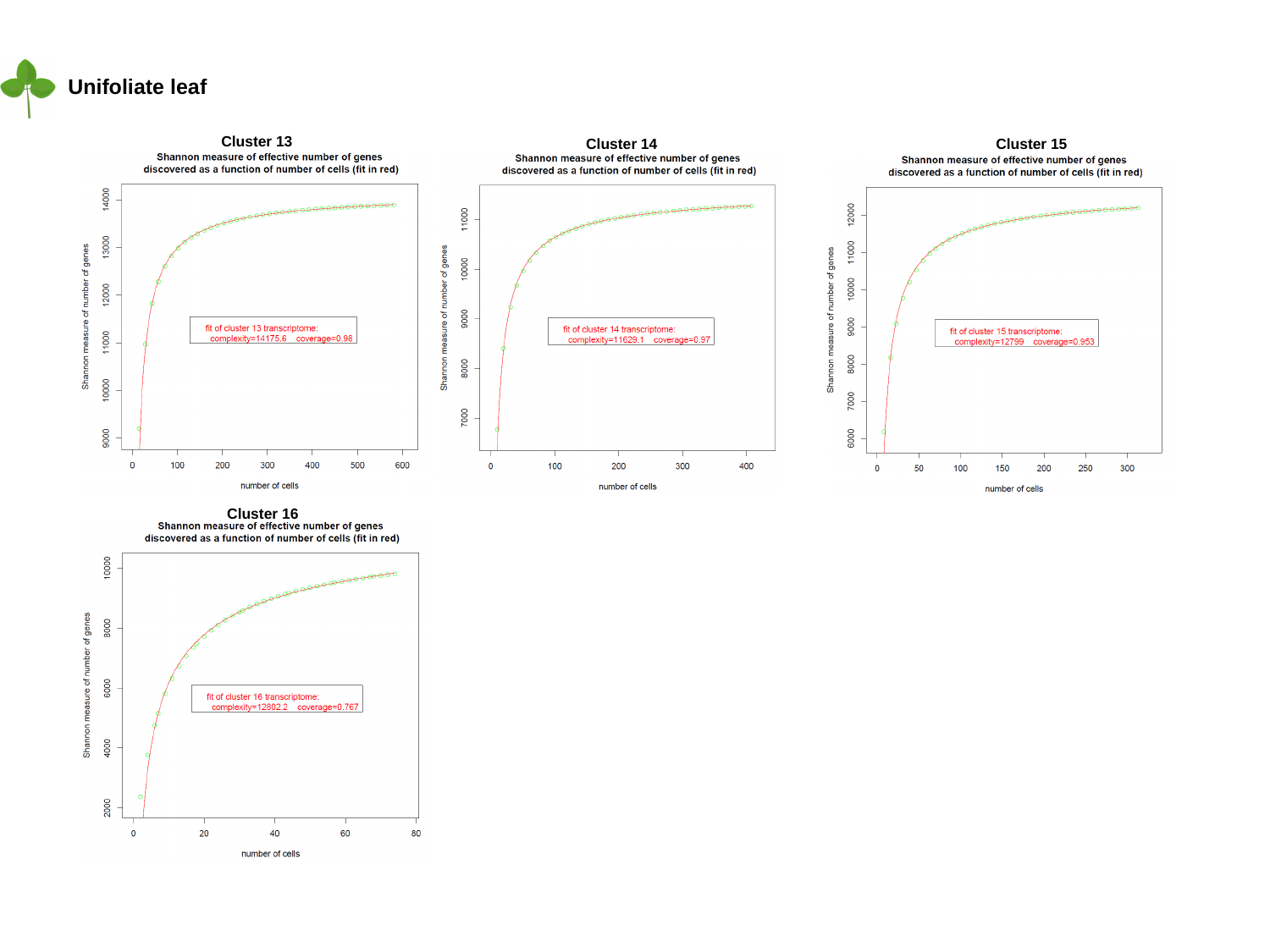

Unifoliate leaf
Cluster 13
Cluster 14
Cluster 15
Cluster 16

### Slide 25
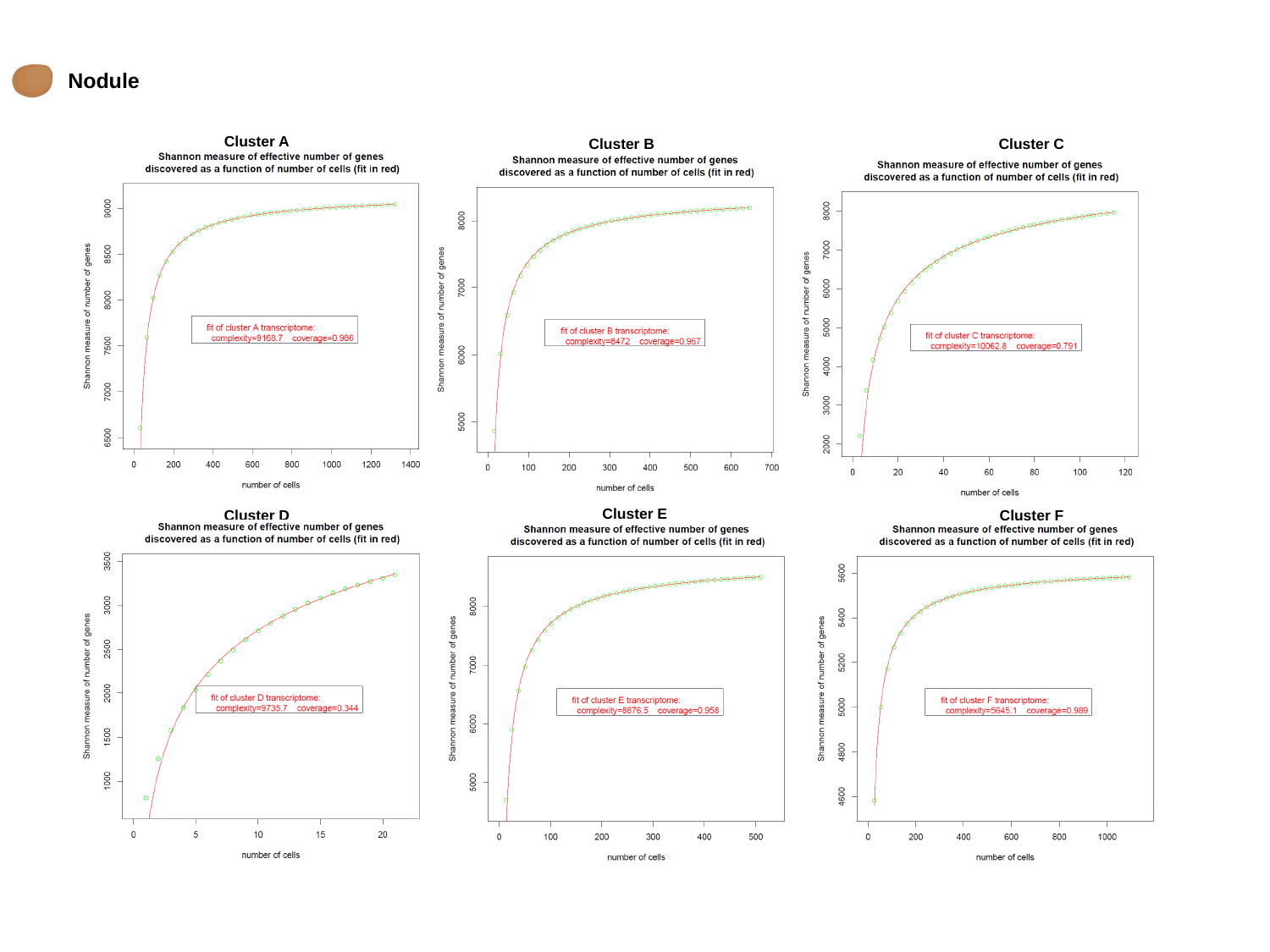

Nodule
Cluster A
Cluster B
Cluster C
Cluster E
Cluster D
Cluster F

### Slide 26
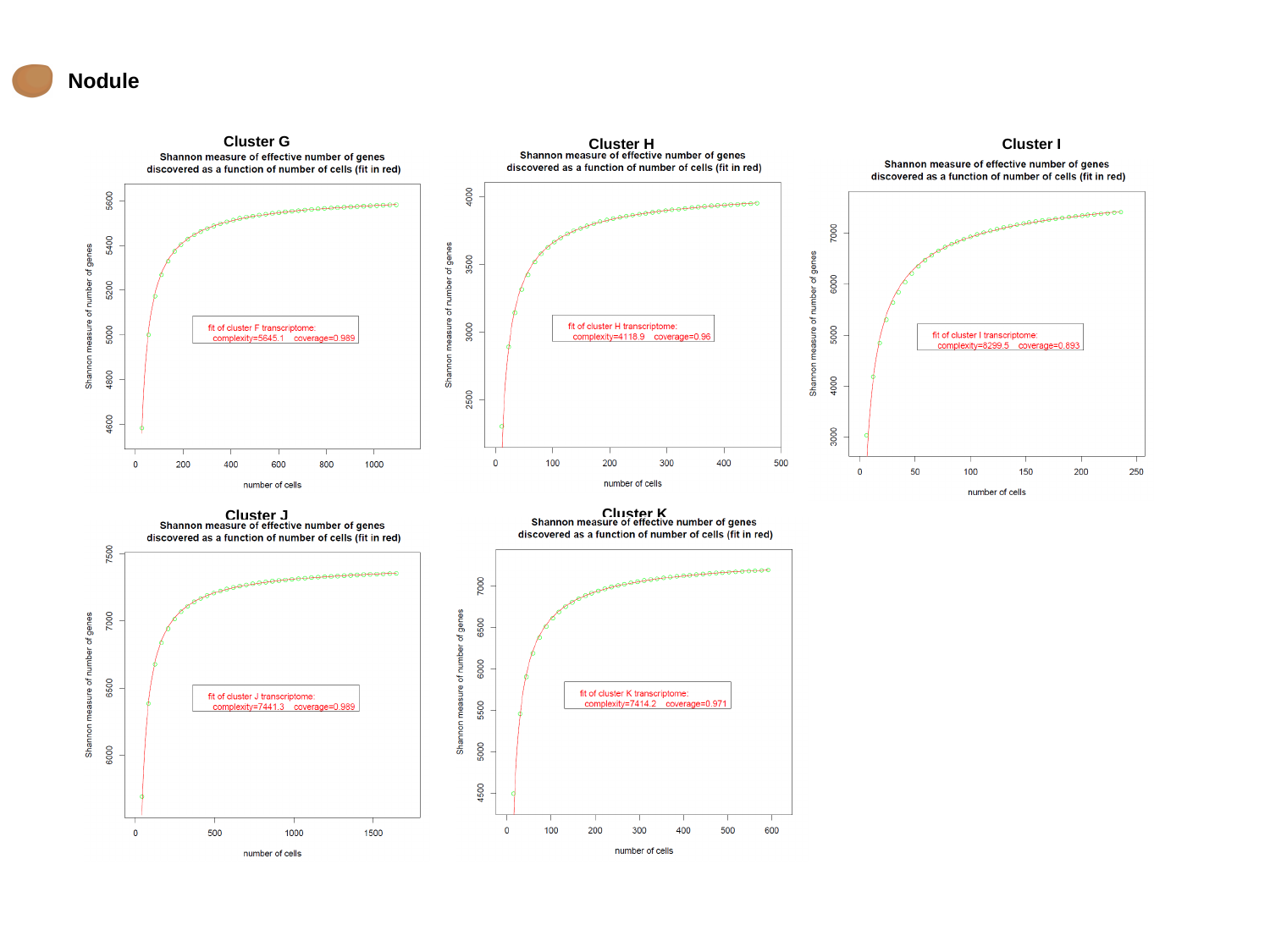

Nodule
Cluster G
Cluster H
Cluster I
Cluster K
Cluster J

### Slide 27
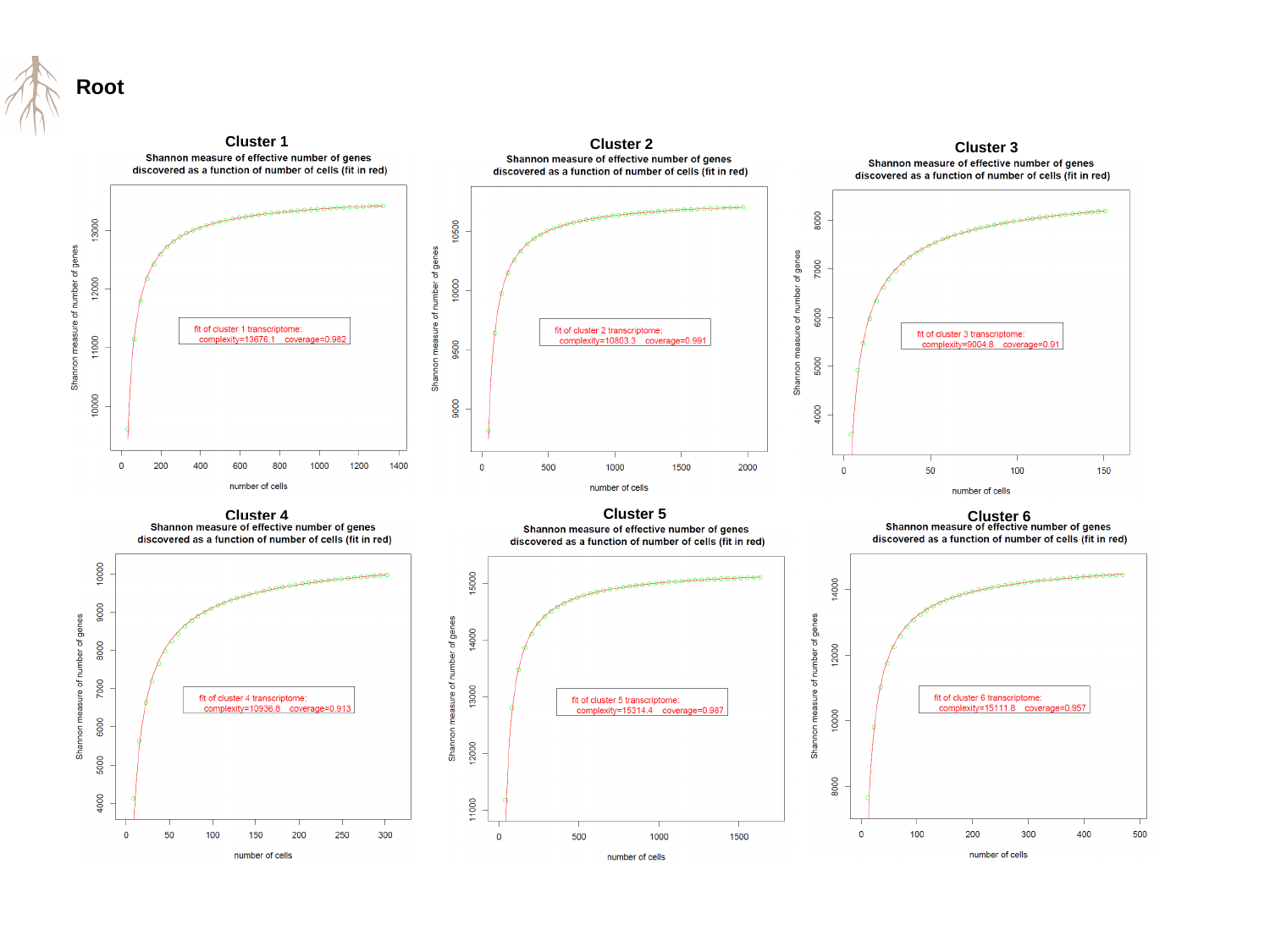

Root
Cluster 1
Cluster 2
Cluster 3
Cluster 5
Cluster 4
Cluster 6

### Slide 28
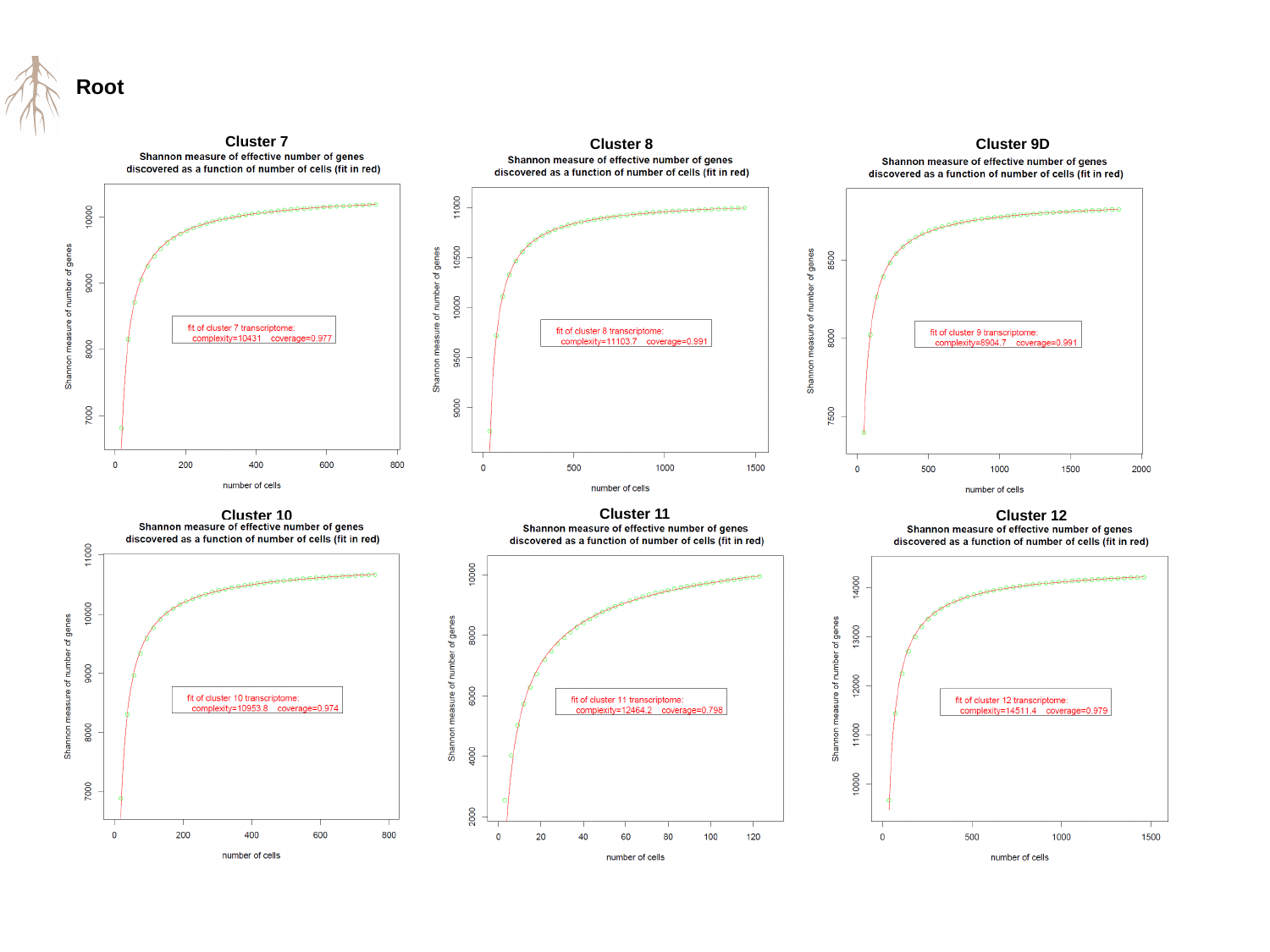

Root
Cluster 7
Cluster 8
Cluster 9D
Cluster 11
Cluster 10
Cluster 12

### Slide 29
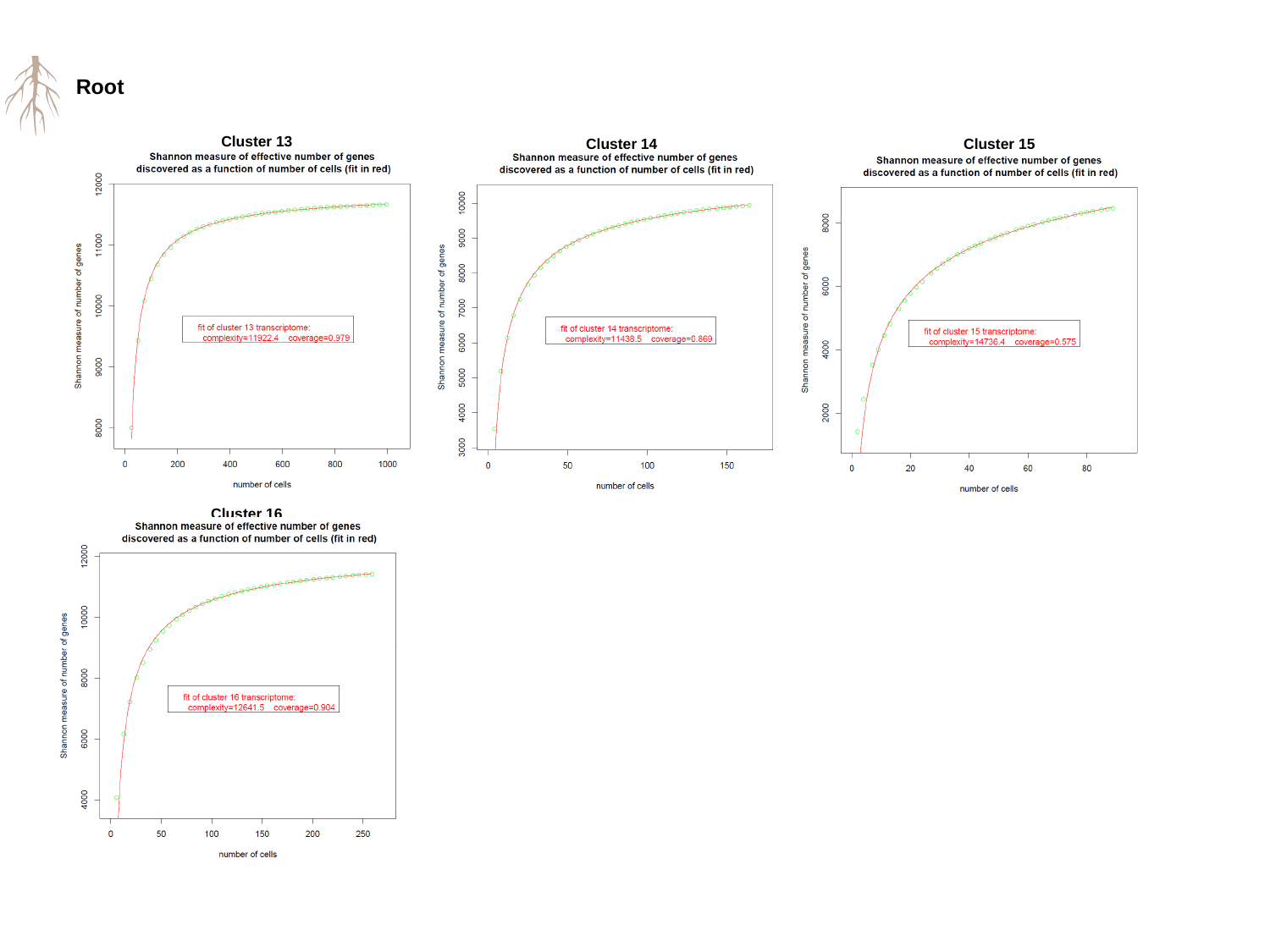

Root
Cluster 13
Cluster 14
Cluster 15
Cluster 16
