## Supplementary material for "Tabula Glycine: The whole-soybean single-cell resolution transcriptome atlas": Other supplemental figures

### Slide 1
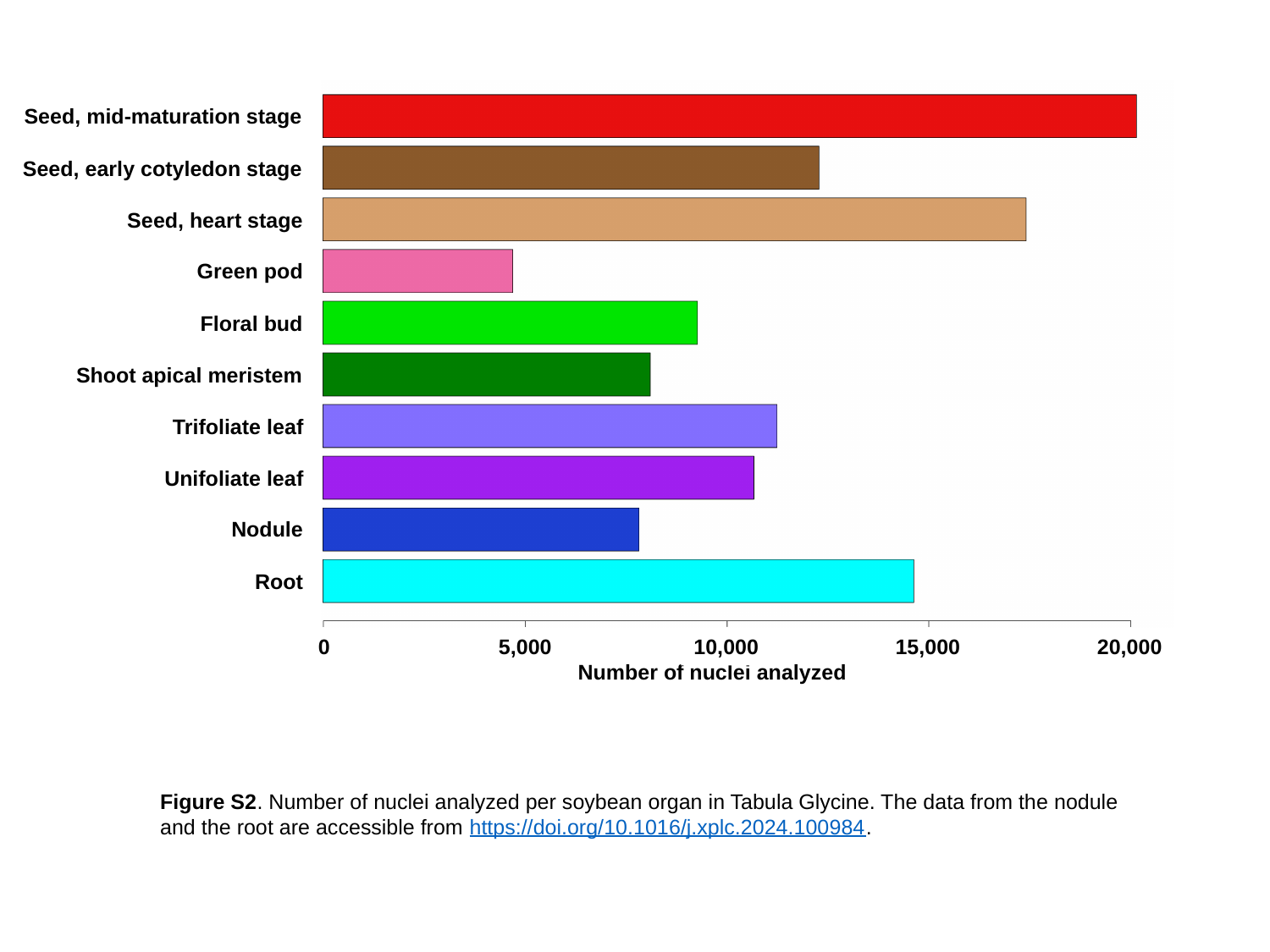

Seed, mid-maturation stage
Seed, early cotyledon stage
Seed, heart stage
Green pod
Floral bud
Shoot apical meristem
Trifoliate leaf
Unifoliate leaf
Nodule
Root
0
5,000
10,000
15,000
20,000
Number of nuclei analyzed
Figure S2. Number of nuclei analyzed per soybean organ in Tabula Glycine. The data from the nodule and the root are accessible from https://doi.org/10.1016/j.xplc.2024.100984.

### Slide 2
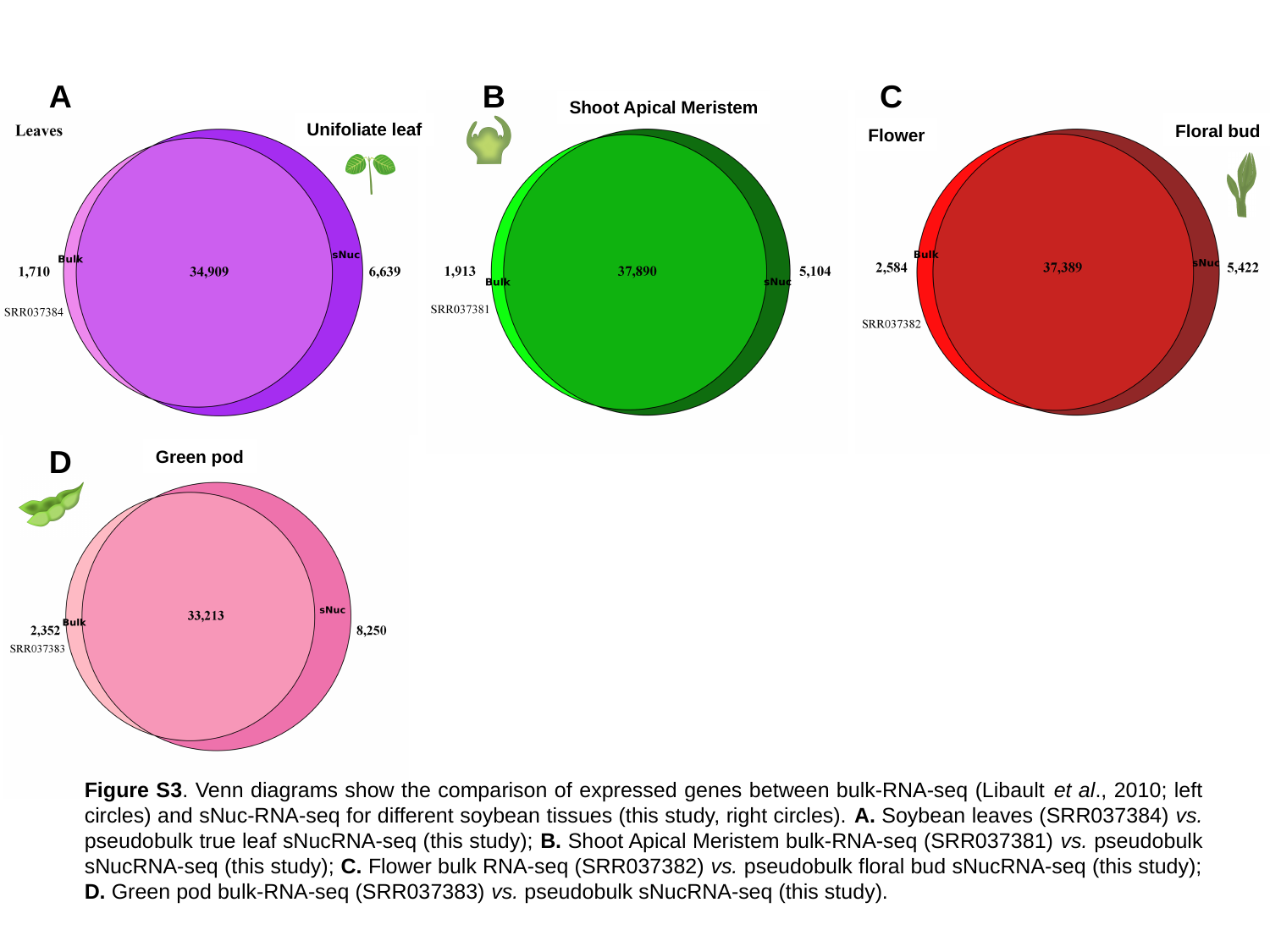

C
A
B
Shoot Apical Meristem
Unifoliate leaf
Floral bud
Flower
D
Green pod
Figure S3. Venn diagrams show the comparison of expressed genes between bulk-RNA-seq (Libault et al., 2010; left circles) and sNuc-RNA-seq for different soybean tissues (this study, right circles). A. Soybean leaves (SRR037384) vs. pseudobulk true leaf sNucRNA-seq (this study); B. Shoot Apical Meristem bulk-RNA-seq (SRR037381) vs. pseudobulk sNucRNA-seq (this study); C. Flower bulk RNA-seq (SRR037382) vs. pseudobulk floral bud sNucRNA-seq (this study); D. Green pod bulk-RNA-seq (SRR037383) vs. pseudobulk sNucRNA-seq (this study).

### Slide 3
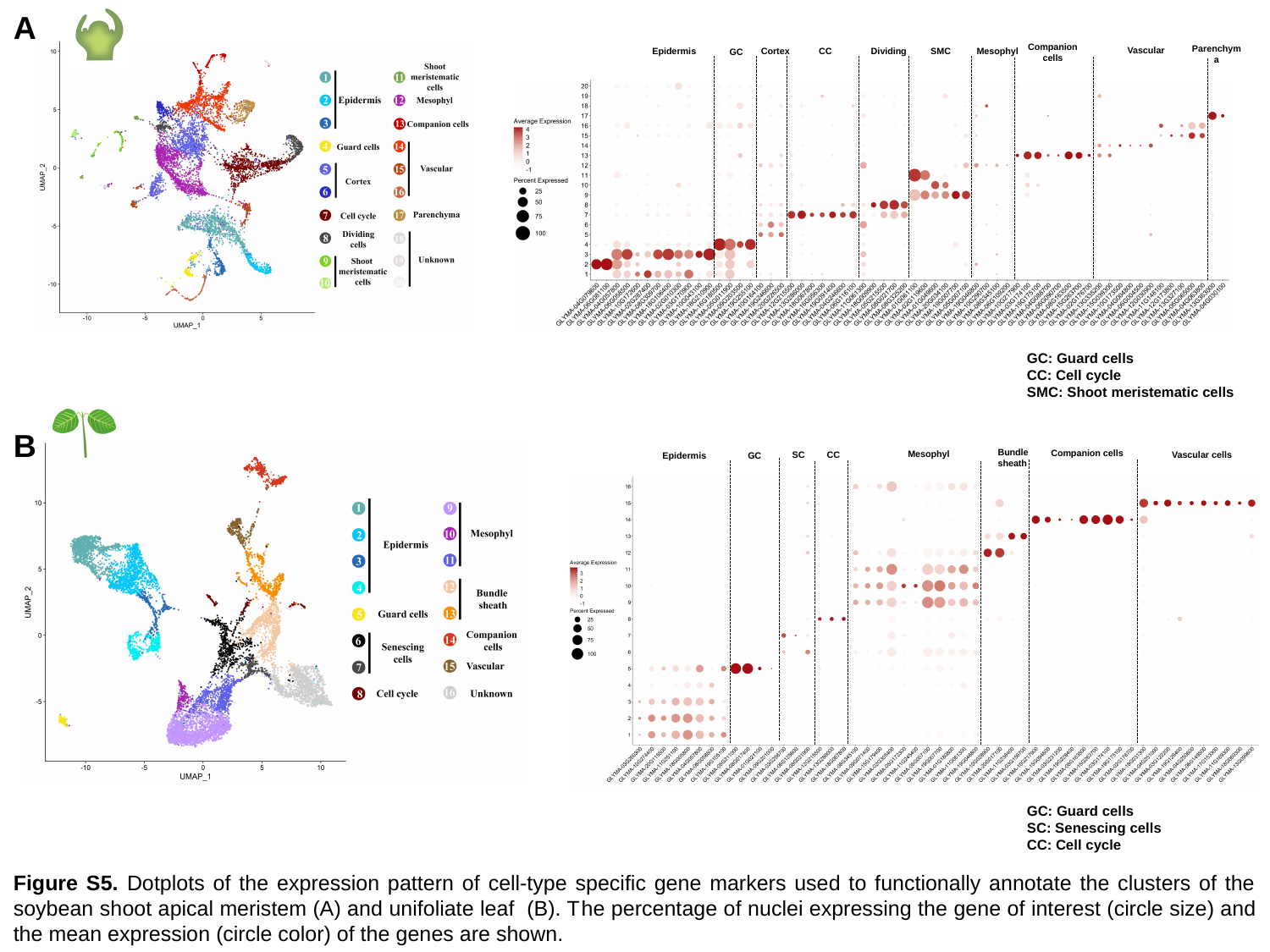

A
Companion cells
Parenchyma
Vascular
Mesophyl
CC
Dividing
SMC
Epidermis
Cortex
GC
GC: Guard cells
CC: Cell cycle
SMC: Shoot meristematic cells
B
Bundle
sheath
Companion cells
Mesophyl
Vascular cells
CC
SC
Epidermis
GC
GC: Guard cells
SC: Senescing cells
CC: Cell cycle
Figure S5. Dotplots of the expression pattern of cell-type specific gene markers used to functionally annotate the clusters of the soybean shoot apical meristem (A) and unifoliate leaf (B). The percentage of nuclei expressing the gene of interest (circle size) and the mean expression (circle color) of the genes are shown.

### Slide 4
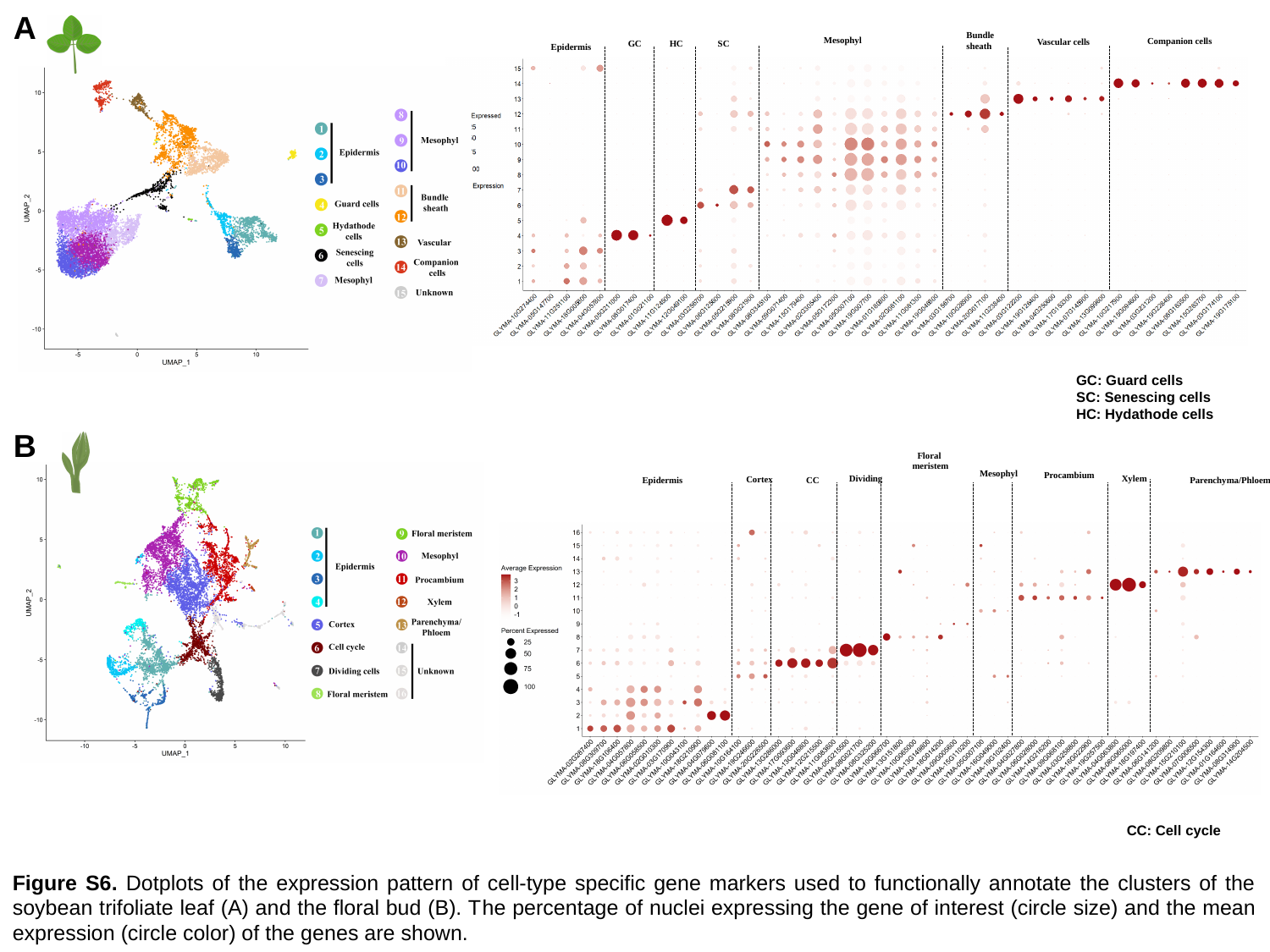

A
Bundle
sheath
Mesophyl
Companion cells
Vascular cells
GC
HC
SC
Epidermis
GC: Guard cells
SC: Senescing cells
HC: Hydathode cells
B
Floral
 meristem
Mesophyl
Procambium
Xylem
Dividing
Cortex
Epidermis
CC
Parenchyma/Phloem
CC: Cell cycle
Figure S6. Dotplots of the expression pattern of cell-type specific gene markers used to functionally annotate the clusters of the soybean trifoliate leaf (A) and the floral bud (B). The percentage of nuclei expressing the gene of interest (circle size) and the mean expression (circle color) of the genes are shown.

### Slide 5
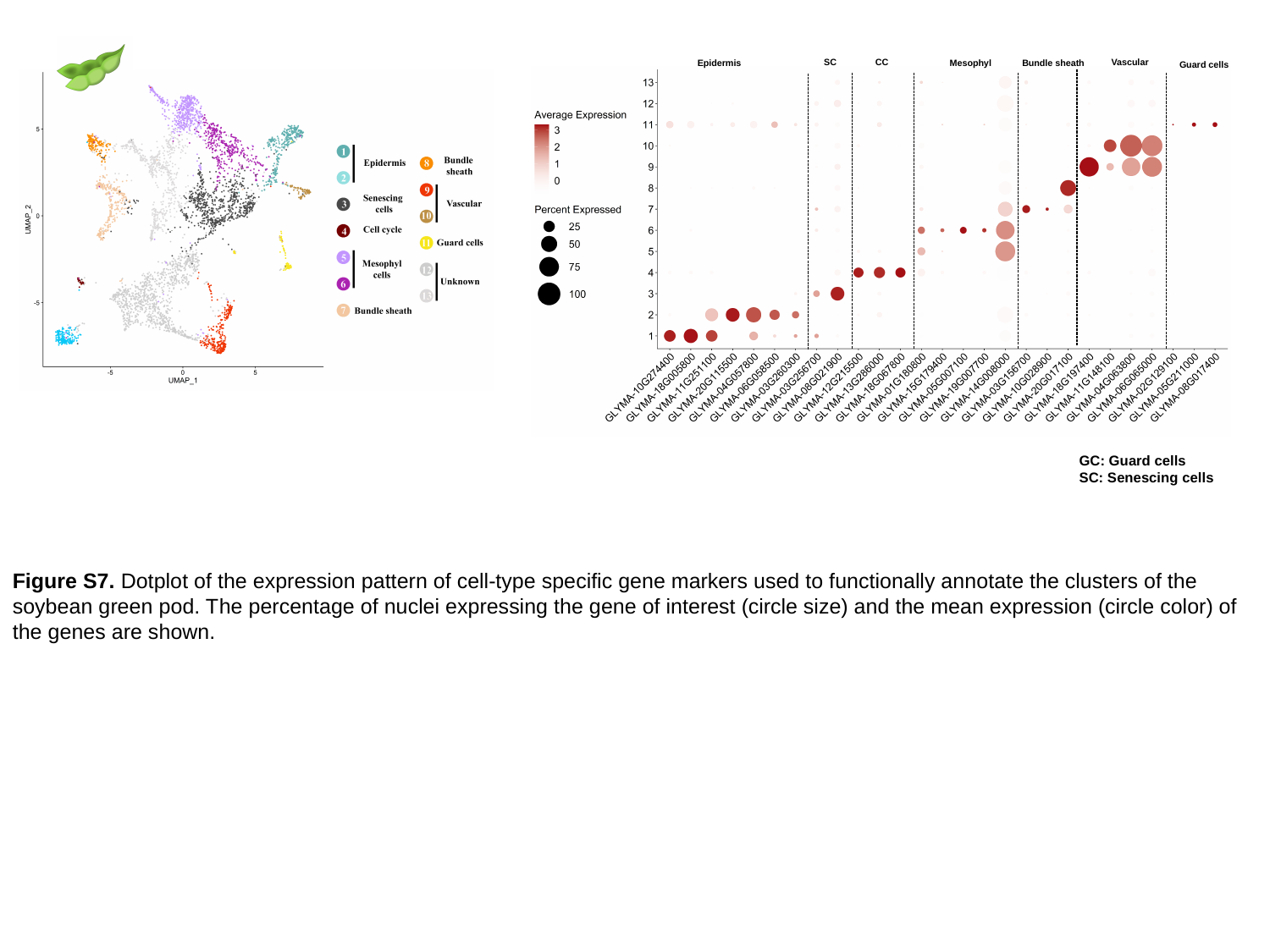

SC
CC
Vascular
Epidermis
Mesophyl
Bundle sheath
Guard cells
GC: Guard cells
SC: Senescing cells
Figure S7. Dotplot of the expression pattern of cell-type specific gene markers used to functionally annotate the clusters of the soybean green pod. The percentage of nuclei expressing the gene of interest (circle size) and the mean expression (circle color) of the genes are shown.

### Slide 6
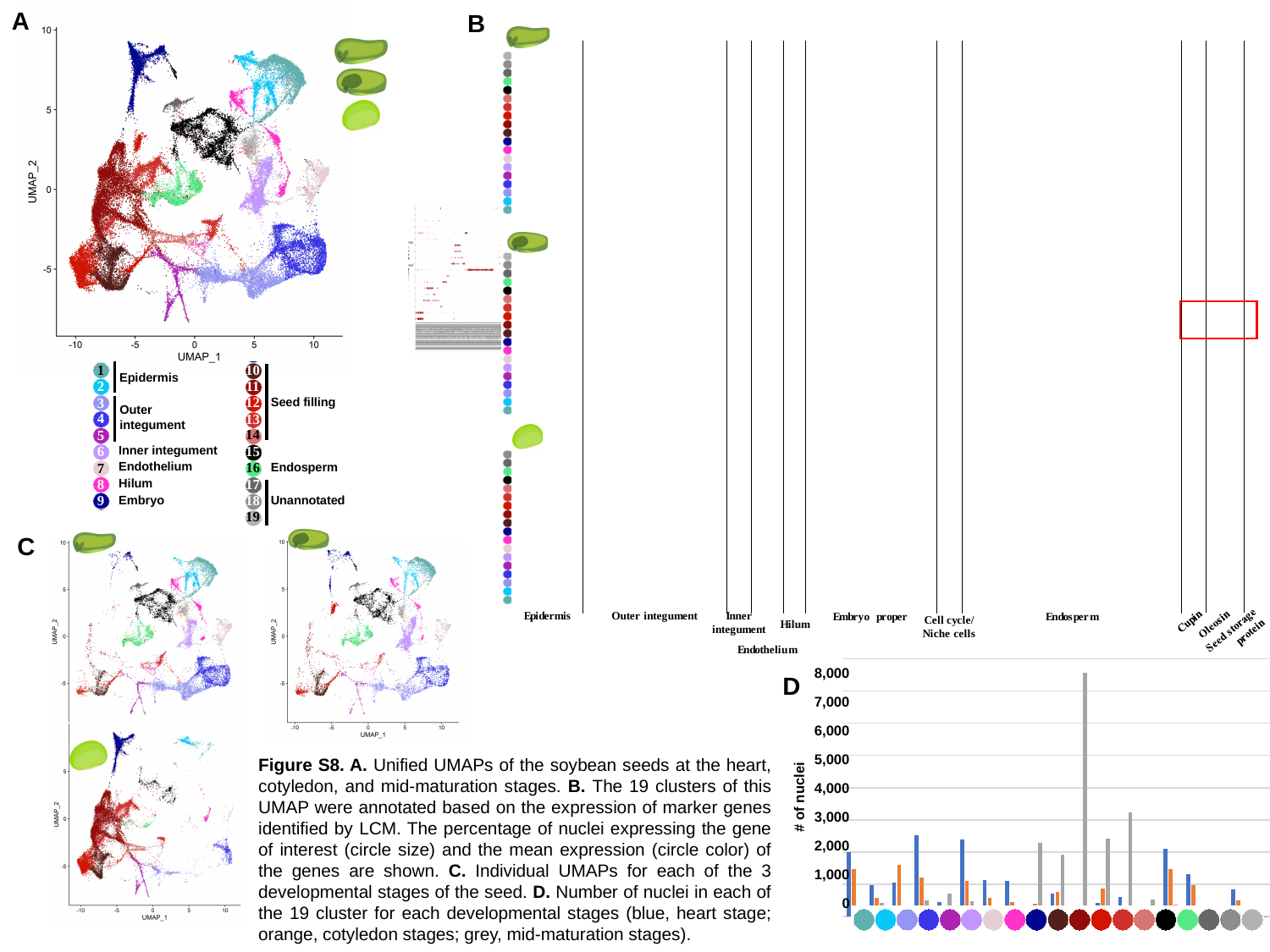

A
B
1
Epidermis
2
3
Outer
integument
4
5
6
Inner integument
Endothelium
7
8
Hilum
9
Embryo
10
11
12
Seed filling
13
14
15
16
Endosperm
17
18
Unannotated
19
C
###### Chart
| Category | a_Seed heart | b_Seed cotyledon | Seed mid-maturation |
|---|---|---|---|
| 1 | 1984.0 | 1461.0 | 96.0 |
| 2 | 973.0 | 565.0 | 412.0 |
| 3 | 1051.0 | 1607.0 | 8.0 |
| 4 | 2528.0 | 1208.0 | 503.0 |
| 5 | 437.0 | 224.0 | 706.0 |
| 6 | 2384.0 | 1089.0 | 479.0 |
| 7 | 1118.0 | 581.0 | 52.0 |
| 8 | 1105.0 | 451.0 | 265.0 |
| 9 | 306.0 | 398.0 | 2285.0 |
| 10 | 701.0 | 750.0 | 1919.0 |
| 11 | 202.0 | 230.0 | 7553.0 |
| 12 | 410.0 | 872.0 | 2418.0 |
| 13 | 615.0 | 257.0 | 3234.0 |
| 14 | 224.0 | 96.0 | 518.0 |
| 15 | 2092.0 | 1465.0 | 371.0 |
| 16 | 1316.0 | 968.0 | 266.0 |
| 17 | 274.0 | 171.0 | 30.0 |
| 18 | 850.0 | 510.0 | 0.0 |
| 19 | 9.0 | 14.0 | 4.0 |8,000
7,000
6,000
5,000
4,000
3,000
2,000
1,000
0
D
#### of nuclei
				Figure S8. A. Unified UMAPs of the soybean seeds at the heart, cotyledon, and mid-maturation stages. B. The 19 clusters of this UMAP were annotated based on the expression of marker genes identified by LCM. The percentage of nuclei expressing the gene of interest (circle size) and the mean expression (circle color) of the genes are shown. C. Individual UMAPs for each of the 3 developmental stages of the seed. D. Number of nuclei in each of the 19 cluster for each developmental stages (blue, heart stage; orange, cotyledon stages; grey, mid-maturation stages).
Supplemental Figure S11. A-C. Ridge plot distributions of the expression of the soybean Oleosin family (A.) and Cupin family genes (B.), and protein storage genes(C.) (Table Sx).
Supplemental Figure S12. A-C. Dot plot of oleosin genes (A.), cupin gene family (B.), and protein storage genes(C.) (Table Sx). D. Re-clustering of seed-filling clusters (Clusters #11-14) showing clustering by developmental stage.
Vascular tissues
Embryo
Epidermis
Seed storage
Outer integument
Suspensor
Hilum
Inner integument
Endothelium
Embryo proper

### Slide 7
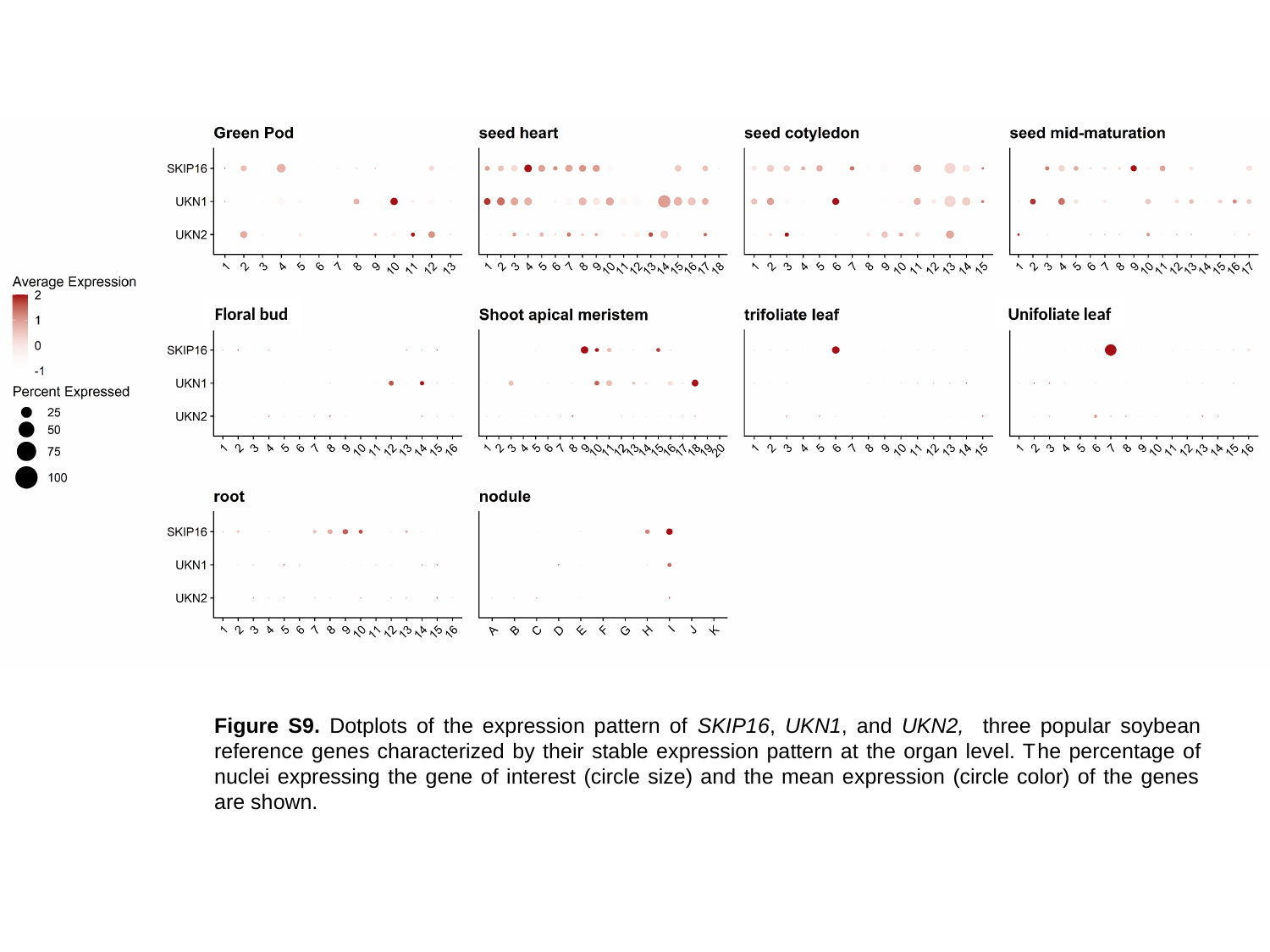

Floral bud
Unifoliate leaf
Figure S9. Dotplots of the expression pattern of SKIP16, UKN1, and UKN2, three popular soybean reference genes characterized by their stable expression pattern at the organ level. The percentage of nuclei expressing the gene of interest (circle size) and the mean expression (circle color) of the genes are shown.

### Slide 8
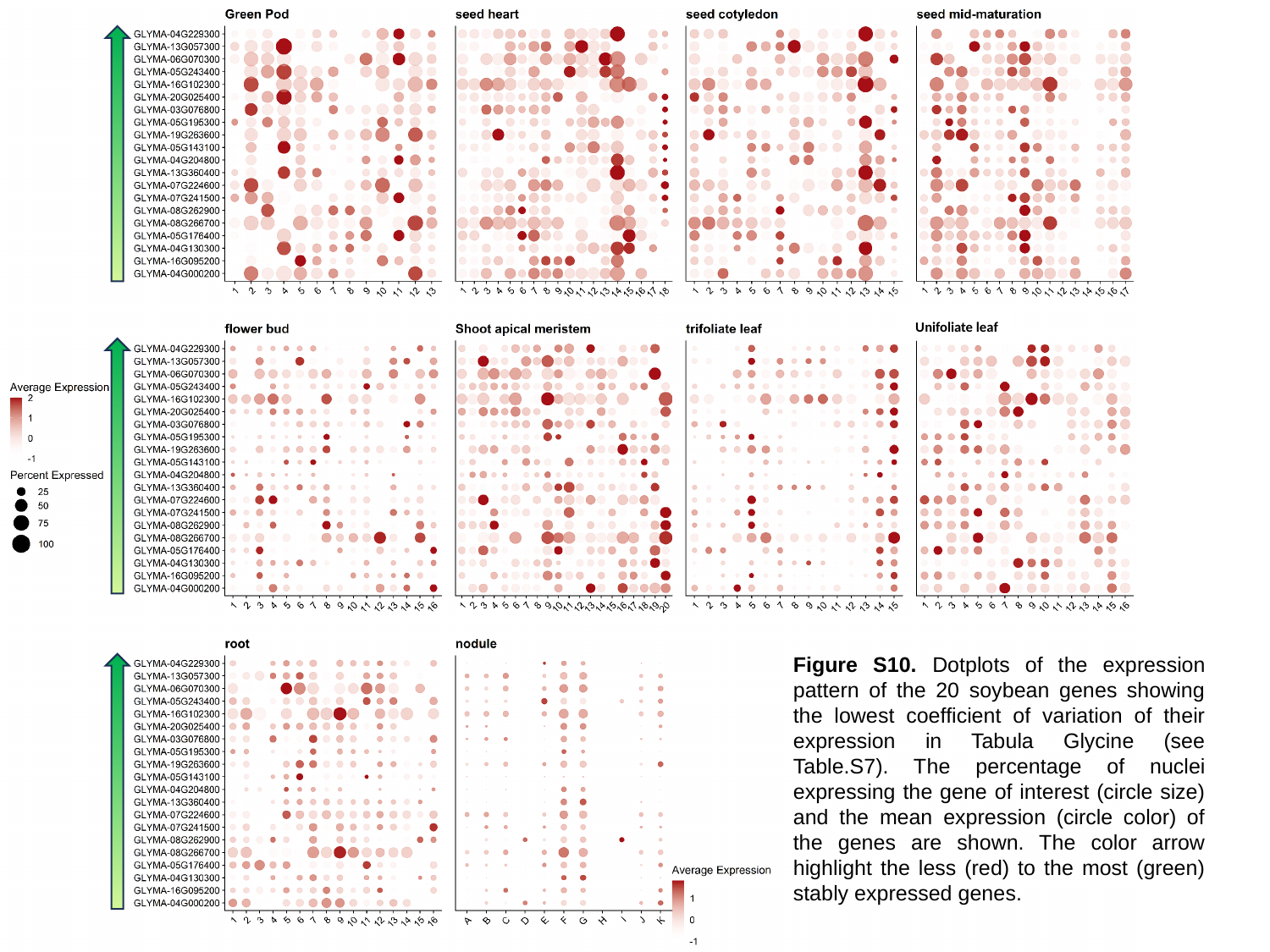

Unifoliate leaf
Figure S10. Dotplots of the expression pattern of the 20 soybean genes showing the lowest coefficient of variation of their expression in Tabula Glycine (see Table.S7). The percentage of nuclei expressing the gene of interest (circle size) and the mean expression (circle color) of the genes are shown. The color arrow highlight the less (red) to the most (green) stably expressed genes.

### Slide 9
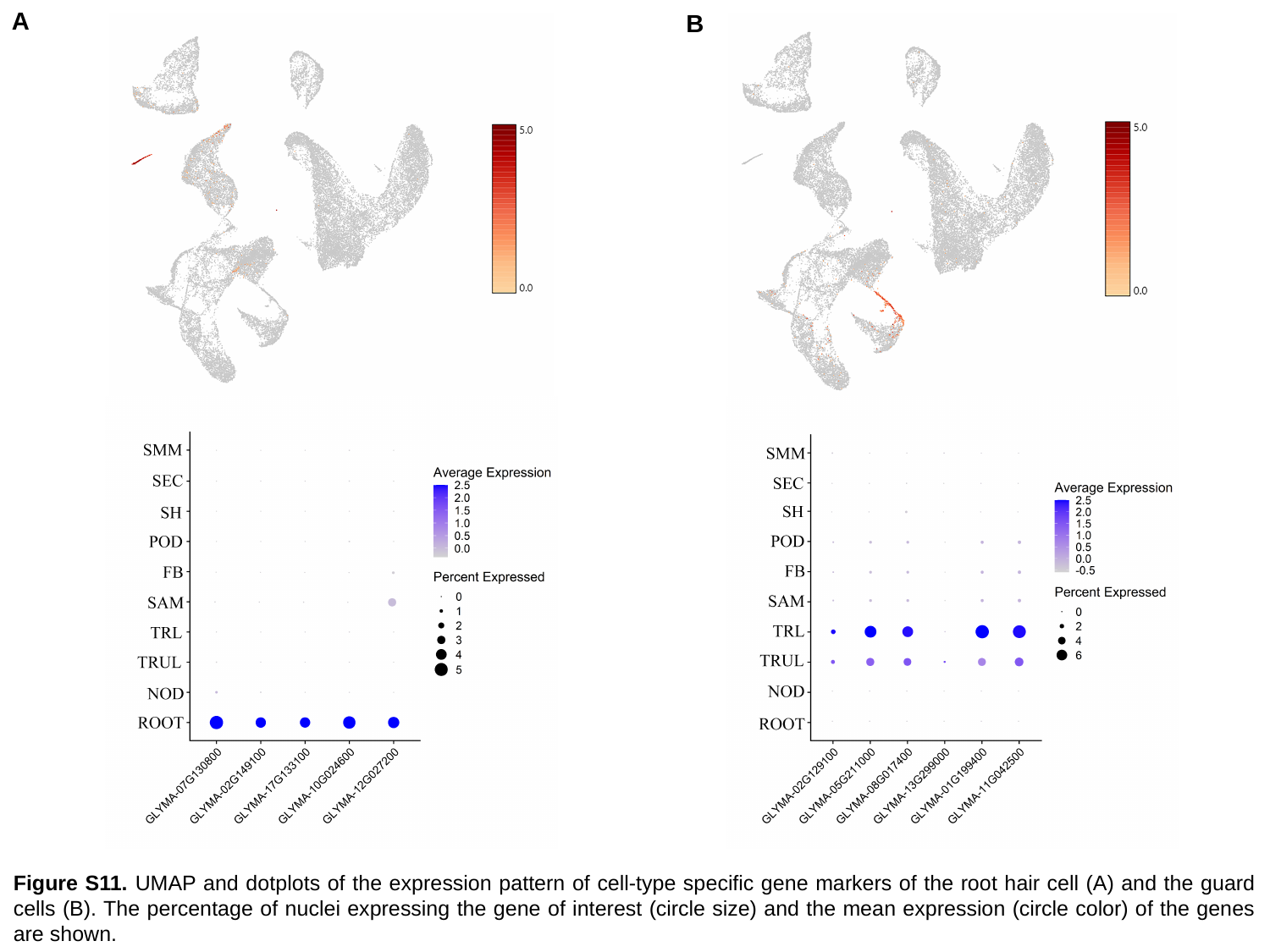

A
B
Figure S11. UMAP and dotplots of the expression pattern of cell-type specific gene markers of the root hair cell (A) and the guard cells (B). The percentage of nuclei expressing the gene of interest (circle size) and the mean expression (circle color) of the genes are shown.

### Slide 10
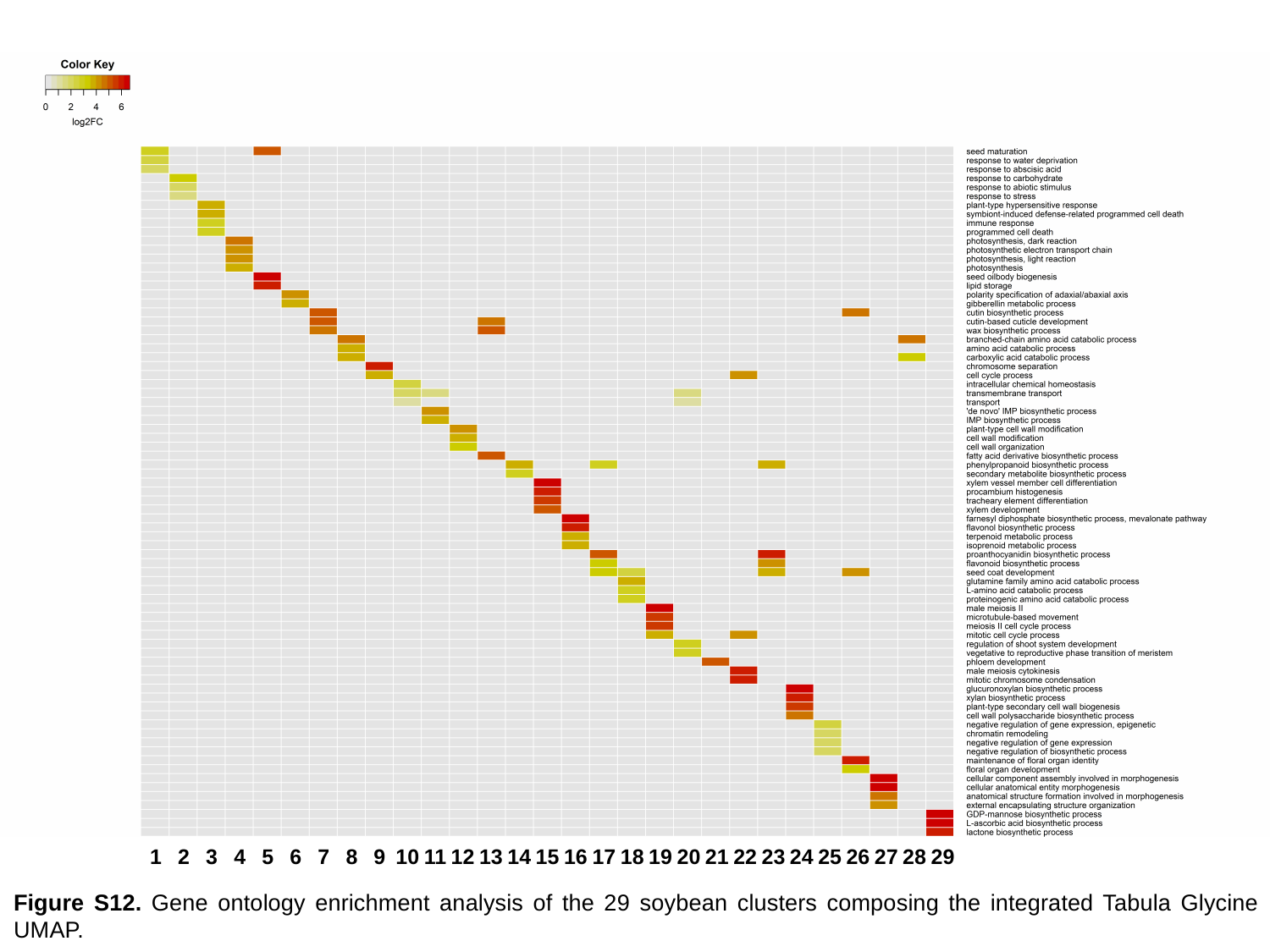

1
2
3
4
5
6
7
8
9
10
11
12
13
14
15
16
17
18
19
20
21
22
23
24
25
26
27
28
29
Figure S12. Gene ontology enrichment analysis of the 29 soybean clusters composing the integrated Tabula Glycine UMAP.
