## Supplementary material for "Tabula Glycine: The whole-soybean single-cell resolution transcriptome atlas": Table S10

**Table S10: Necessary reported information to allow evaluation and repetition of a plant single cell/nucleus experiment.**

|  | **Details** | **Experimental information** |
| --- | --- | --- |
| **Biological material** | | |
| Species | Glycine max |  |
| Accession | Williams 82 |  |
| Tissue type | Root, nodule, true leaves, trifoliate, pods, seeds, shoot apical and floral meristems. |  |
| Detailed growth conditions | Greenhouse (day/night: 16/8 hrs, 26/20C) |  |
| Harvest conditions | Collected and process within 5 minutes |  |
| **Sample preparation** | | |
| Isolation protocol | Thibivilliers et al., 2021 |  |
| Tissue dissection | N.A. |  |
| Fixation | N.A. |  |
| Cell/nuclei enrichment | Nuclei purified by FANS |  |
| Total sample preparation time | 1-2 hours |  |
| Estimated cell/nuclei number loaded | Estimated of 5,000 targeted |  |
| Instrument/Method/Kit | Thibivilliers et al., 2021 |  |
| Cell viability test | N.A. |  |
| **Libraries** | | |
| Library construction | sNucRNa-seq 3’ |  |
| Amplification method | 10X Genomics |  |
| **Sequence results** | | |
| Instrument/method | Novaseq S4 |  |
| Library layout/paired-end | Paired-end dual indexes |  |
| N° sequenced reads | Targeted 30-50,000 reads per nucleus |  |
| **Raw data** | | |
| Reference genome | Glycine_max v2.1 ([Index of /pub/plants/release-52/fasta/glycine_max/dna (ebi.ac.uk)](https://ftp.ensemblgenomes.ebi.ac.uk/pub/plants/release-52/fasta/glycine_max/dna/)) |  |
| Annotation version | Glycine max v2.1.52 ([Index of /pub/plants/release-52/gtf/glycine_max (ebi.ac.uk)](https://ftp.ensemblgenomes.ebi.ac.uk/pub/plants/release-52/gtf/glycine_max/)) |  |
| Mapping method (incl. software, customized settings) | CellRanger v6.1.2. This program uses STAR predetermined software. Customized: --include-introns=true |  |
| Mapping efficiency | Seed mid-maturation avg: 82.4%  Seed early cotyledon avg: 86.6%  Seed heart avg: 88.2%  Green Pod avg: 77.65%  Flower bud avg: 60.55%  SAM avg: 72%  Trifoliate leaves avg: 89.75%  True leaves avg: 86.4%  Root avg:78.4%  Nodule avg: 58.2% |  |
| Sequencing saturation | Seed mid-maturation avg: 42.4%  Seed early cotyledon avg: 57.25%  Seed heart avg: 60.3%  Green Pod avg: 70.5%  Flower bud avg: 77.2%  SAM avg: 81.2%  Trifoliate leaves avg: 80.35%  True leaves avg: 75.9%  Root avg:83.2%  Nodule avg: 45.7% |  |
| Estimation of ambient RNA | SoupX (Young and Behjati, 2020. [10.1093/gigascience/giaa151](https://doi.org/10.1093/gigascience/giaa151)) |  |
| Imputation method and settings | N.A. |  |
| **Processed data** | | |
| N° captured cells/nuclei | Seed mid-maturation: 21,776  Seed early cotyledon: 13,586  Seed heart:19,901  Green pod:4,962  Flower bud:10,317  SAM:8,637  Trifoliate leave:16,183  True leave:11,373  Nodule:19,217  Root:14,776 |  |
| N° high quality cells/nuclei | Seed mid-maturation: 20,154  Seed early cotyledon:12,288  Seed heart:17,417  Green pod:3,457  Flower bud:9,728  SAM:8,108  Trifoliate leave:11,251  True leave:10,667  Nodule:7,830  Root:14,639 |  |
| Filter criteria: % mitochondrial reads/cell or nucleus | N.A |  |
| Filter criteria: % chloroplast reads/cell or nucleus | N.A |  |
| Filter criteria: Minimum N° UMI/cell or nucleus | >=500 UMI |  |
| N° total detected transcripts | Number of expressed genes  Seed mid-maturation: 43,356  Seed early cotyledon:46,545  Seed heart:47,309  Green pod:41,463  Flower bud:42,811  SAM:42,994  Trifoliate leave:43,984  True leave:41,458  Nodule:37,118  Root:42,391 |  |
| Doublet rate | N.A (We used DoubletDetection) |  |
| Replicate comparisons | Correlation |  |
| Batch correction method for merging (incl. reasoning for batch correction) | N.A |  |
| **Validation** | Method of automatic annotation of clusters | N.A |
|  | Method of manual annotation (markers, gene function info) | See supplemental |
|  | Verification in planta (e.g. Number of markers used for validation) | Using molecular cartography (See Supplemental) |
| **Data availability** | Analysis scripts & codes (GitHub) | N.A |
|  | Excel Tables DEG for each cluster | Included at www.tabulaglycine.org |
|  | Objects/count matrix in repository (which one, where?) | GEO/NCBI  Bioprojects: PRJNA983388  PRJNA938968  GSE226149 (mtx file)  GSE226149 (mtx file) |
|  | On-line tool/browser URL | www.tabulalycine.org |
|  | Cell-level metadata table | Included at www.tabulaglycine.org |
| **Additional** | additional comments from the authors |  |
