## Supplementary figures and images for "Tabula Glycine: The whole-soybean single-cell resolution transcriptome atlas"

### Video S1

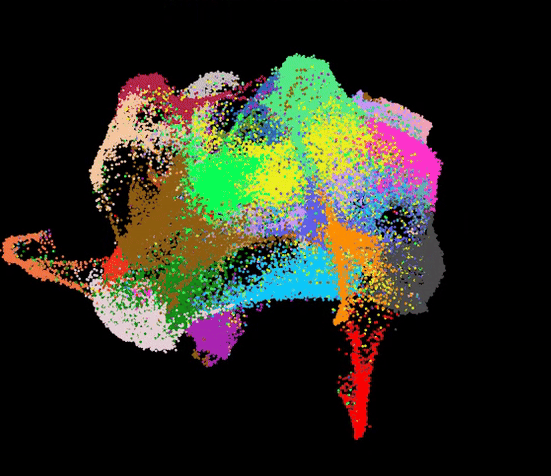
